# Quantifying the effects of cell death and agar density on yeast colony biofilms using an extensional-flow mathematical model

**DOI:** 10.1101/2025.10.04.680491

**Authors:** Alexander K. Y. Tam, Daniel J. Netherwood, Jennifer M. Gardner, Jin Zhang, Campbell W. Gourlay, Vladimir Jiranek, Benjamin J. Binder, J. Edward F. Green

**Affiliations:** UniSA STEM, The University of South Australia, Mawson Lakes SA 5095, Australia; School of Computer and Mathematical Sciences, The University of Adelaide, Adelaide SA 5005, Australia; Department of Wine Science, School of Agriculture, Food and Wine, The University of Adelaide, Glen Osmond SA 5064, Australia; Kent Fungal Group, School of Biosciences, The University of Kent, Canterbury, Kent, United Kingdom; School of Biological Sciences, The University of Southampton, Southampton, United Kingdom

**Keywords:** Lubrication theory, thin-film approximation, viscous flow, multiphase flow, slip, *Saccharomyces cerevisiae*, parameter estimation

## Abstract

We use a combination of experiments, mathematical modelling, and parameter estimation to better understand how agar density affects colony biofilm growth of the yeast species *Saccharomyces cerevisiae*. We obtained 15 total experimental replicates on rectangular plates filled with 0.6%, 0.8%, 1.2%, and 2.0% agar. In the experiments, we measured the horizontal expansion over time, the number of living cells, and the colony-biofilm aspect ratio. These measurements quantify the colony-biofilm size, composition, and shape, respectively. We modelled colony-biofilm expansion using a thin-film extensional-flow mathematical model. By fitting five unknown model parameters to mean experimental data, we show that nutrient uptake decreases and biofilm–substratum adhesion strength increases with an increase in agar density. Sensitivity analysis, fitting to individual replicates, and synthetic-data analysis confirmed that increased biofilm–substratum adhesion is the most consistent effect of increased agar density. This finding aligns with similar results reported for bacteria, and suggests that substratum properties are important for yeast-colony-biofilm growth.

## 1 Introduction and Background

Biofilms are surface-dwelling communities of cells embedded within a viscous extracellular matrix (ECM) composed of extracellular polymeric substances (EPS) [1–3]. The bakers’ yeast *Saccharomyces cerevisiae* is an important model organism in cell biology [4, 5], because of its ease of use in experiments and its similarity to higher eukaryotic cells [6]. Reynolds and Fink [7] developed an assay for *S. cerevisiae* biofilm formation, which requires the Flo11p glycoprotein known to promote cell-to-surface adhesion. Grown on agar media, the yeast forms a confluent mat that can fill a 90mm Petri dish in under two weeks [8–10]. These mats, which we refer to as colony biofilms, adopt circular shapes with smooth or undulate boundaries [7, 9], rather than the filamentous or dendritic structures formed by other *S. cerevisiae* colonies [11]. Colony biofilms are less complex than naturally-occurring biofilms, but still involve extracellular matrix deposition. Due to similarities between yeast and the bacterium *Mycobacterium smegmatis* [12], Reynolds and Fink [7] hypothesised that the yeast colony biofilm expands by sliding motility. This term refers to passive growth [13], where the cells and fluid expand as a unit, with low surface tension and weak adhesion to the substratum. Mathematical modelling, when combined with experimental observations, enables us to better understand these expansion mechanisms and dynamics.

Mathematical modelling of colony-biofilm growth is an established field, encompassing agent-based, reaction–diffusion, and continuum-mechanical approaches [14, 15]. The earliest and most common models consider growth in a liquid culture medium from which the biomass obtains nutrients [16–18]. In contrast, our yeast colony biofilms spread over an agar medium, bounded above by air. This scenario has been investigated previously by several authors [19–26]. For *S. cerevisiae* specifically, Tam et al. [19] developed a thin-film extensional-flow mathematical model for colony-biofilm expansion by sliding motility [19]. However, a limitation of this model is that Tam et al. [19] mostly neglected cell death when comparing the model with experiments. In harsh environments, yeast cells will die by accidental cell death [27, 28], but can also undergo regulated cell death [29, 30], which releases nutrient for the benefit of the colony. Another limitation is that Tam et al. [19] only considered experiments on low-density (0.3%) agar. Recent work [31–33] has explored how agar density and cell–substratum adhesion impact growth in bacterial colonies [31, 32, 34, 35]. These works have revealed that high-density substrates tend to reduce nutrient uptake [35], increase cell–substrate friction [32], promote wrinkle formation [34], and favour vertical over horizontal growth, increasing the biofilm–agar contact angle [32]. Whilst we expect these effects of varying agar density to occur in *S. cerevisiae* colonies as well, their prevalence and underlying mechanisms remain largely unexplored in fungal colonies.

We combine experiments and mathematical modelling to quantify the effects of cell death and agar density in *S. cerevisiae* yeast colony biofilms. In Section 2, we describe our rectangular experimental assay and the image processing procedure. These methods yielded a total of 15 experimental colony biofilms on 0.6%, 0.8%, 1.2%, and 2.0% agar. All replicates were stained using Phloxine B dye (10 µm), to indicate locations of cell death [36, 37]. We obtained measurements for colony-biofilm width over time, cell-viability data at three spatial locations, and a measure of the aspect ratio. Like in bacterial colonies [32], in our experiments higher-density agar favoured vertical growth over horizontal growth.

To interpret these new experimental results, we modify the thin-film extensional-flow model of Tam et al. [19] to incorporate colony-biofilm composition data and the impact of cell–substrate adhesion. The model assumes that the agar substratum is rigid. Our objective is to quantify how agar density affects parameters related to cell proliferation, nutrient uptake and consumption, and biofilm–substratum adhesion. Using numerical optimisation, we fit five unknown model parameters to the experimental data, obtaining a set of optimal parameters for each agar density. Consistent with previous studies in bacteria [32, 35], increased agar density reduces nutrient uptake and increases cell–substratum friction. In contrast, biomass-production rate, cell-death rate, and nutrient-consumption rate exhibit less variability across agar densities. We confirm the robustness of these results using parameter-sensitivity analysis and synthetic simulations. These analyses suggest that differences in cell–substratum adhesion are robust, whereas differences in nutrient uptake are less robust. Thus, we confirm that cell–substratum adhesion is an important mechanism explaining differentiated growth of *S. cerevisiae* colony biofilms on varying density media.

## 2 Experiments and Image Processing

### 2.1 Experimental Method

Σ1278b, a diploid prototrophic strain of *Saccharomyces cerevisiae*, was used in these experiments. Yeast Peptone Dextrose (YPD) medium (10 g L^−1^ yeast extract, 20 g L^−1^ peptone, 20 g L^−1^ glucose) either as a liquid or with 0.6%, 0.8%, 1.2%, and 2.0% agar for solid media was prepared by filter sterilising 2 × YPD and mixing with an equal volume of molten 12, 16, 24 or 40 g L^−1^ (BD Bacto™ Agar, Becton, Dickinson and Company, New Jersey, USA). Phloxine B was also added to YPD agar at 10 µm. A 5 mL yeast culture was grown for 24 hours in liquid YPD prior to inoculation on plates. A straight streak of yeast culture was applied to YPD agar in a 100mm square Petri dish with a 4.5mm plastic inoculation loop guided along a suspended ruler edge. The initial cell density was 1.2 × 10^8^ cells mL^−1^ and the streak width approximately 2mm. Plates were cling wrapped and incubated agar side down for up to 21 days at 25 °C. Macroscopic plate images were captured on Days 2, 4, 6, 8, 10, 12, 14, and 21, using an Apple iPhone 12 Pro. Cross sections that spanned the entire width of the colony-biofilm streak (and 5mm tall) were cut from plates with a scalpel and imaged using a Nikon Eclipse 50i microscope and Nikon camera (DS-2MBW, Nikon, Japan). Multiple images were taken across each section and stitched together with Fiji software (ImageJ).

The proportion of stained cells was measured for samples from two plates (one each on 0.6% and 2.0% agar), on Day 14 of the experiments. This was achieved using flow cytometry with a Guava 12 HT system and Guava EasyCyte™ software (Luminex, Yokohama, Japan; formerly Millipore, Darmstadt, Germany). The flow cytometry was equipped with Violet (405nm), Blue (488nm, 150 mW), Red (642nm) excitation lasers, along with 450/45nm and 664/20nm emission filters. Cell samples were collected with sterile pipette tips or washed from 2 cm^2^ or 4 cm^2^ agar slices spanning the entire width of the area containing yeast and diluted in 5mL phosphate buffered saline. Cells fluorescent due to the uptake of Phloxine B were considered non-viable. In parallel, the total number of cells was also enumerated on Day 21 using a hemocytometer and a Nikon Eclipse 50i microscope. Total and viable cell enumeration was performed on a representative sample of cells washed from a 4 cm^2^ agar slice. Counting was performed by both Flow Cytometry, as well as manual counting with a hemocytometer. After washing, plates were inspected by microscopy and it was noted that all cells washed off and there were no cells invading the agar. Photographs of the sampled regions used to obtain the cell viability and cell count measurements are available in the Supporting Material.

### 2.2 Image Processing and Quantification

We obtained a time series of photographs of the yeast colony biofilm for each experimental replicate. Example photographs from one replicate on 0.6% agar and one replicate on 2.0% agar are shown in Figure 1. To obtain the colony-biofilm width in an experimental photograph, we used the known Petri-dish size of 100mm to determine the physical distance corresponding to a pixel. After obtaining this scaling factor, we manually rotated and cropped each photograph to remove the Petri dish from the image. We then used Julia’s `FileIO` package and `Images` library to import the photograph and apply Otsu’s method to obtain a binary image. We then computed the total number of visible pixels in the colony biofilm, divided by the number of pixels in the *y*-direction (as per the axes shown on Figure 1) of the photograph, and applied the scaling factor, to determine an approximation for the colony-biofilm width. The mean colony-biofilm half-widths over time for all agar densities are shown in Figure 2.x

**Figure 1:**
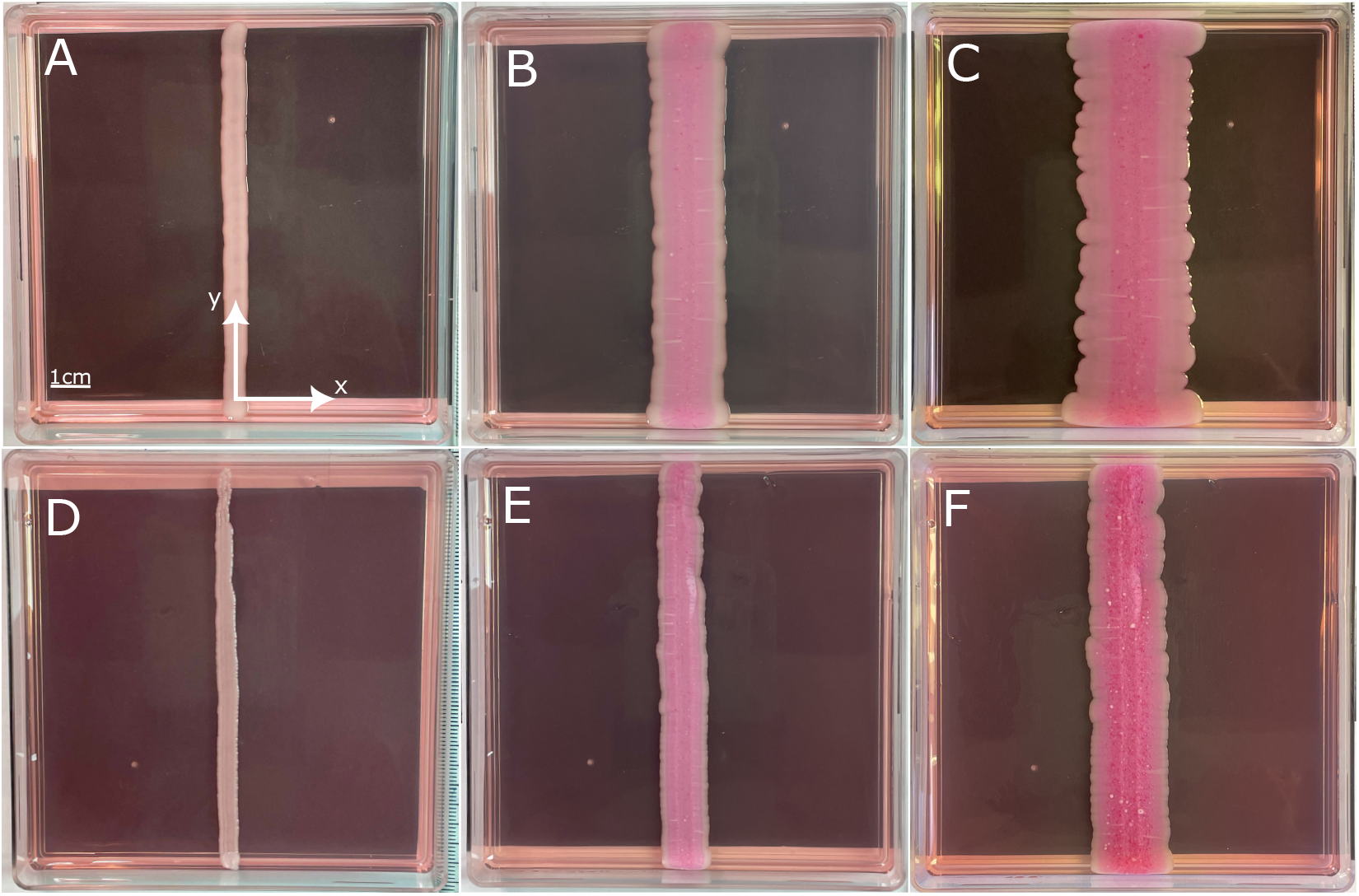
Experimental photographs of yeast-colony-biofilm growth over 21 days on 0.6% and 2.0% agar. Darker pink indicates regions of dead cells stained with Phloxine B. (A) Colony biofilm grown on 0.6% agar (Day 2), with axes and scale bar included. (B) Colony biofilm grown on 0.6% agar (Day 10). (C) Colony biofilm grown on 0.6% agar (Day 21). (D) Colony biofilm grown on 2.0% agar (Day 2). (E) Colony biofilm grown on 2.0% agar (Day 10). (F) Colony biofilm grown on 2.0% agar (Day 21).

**Figure 2:**
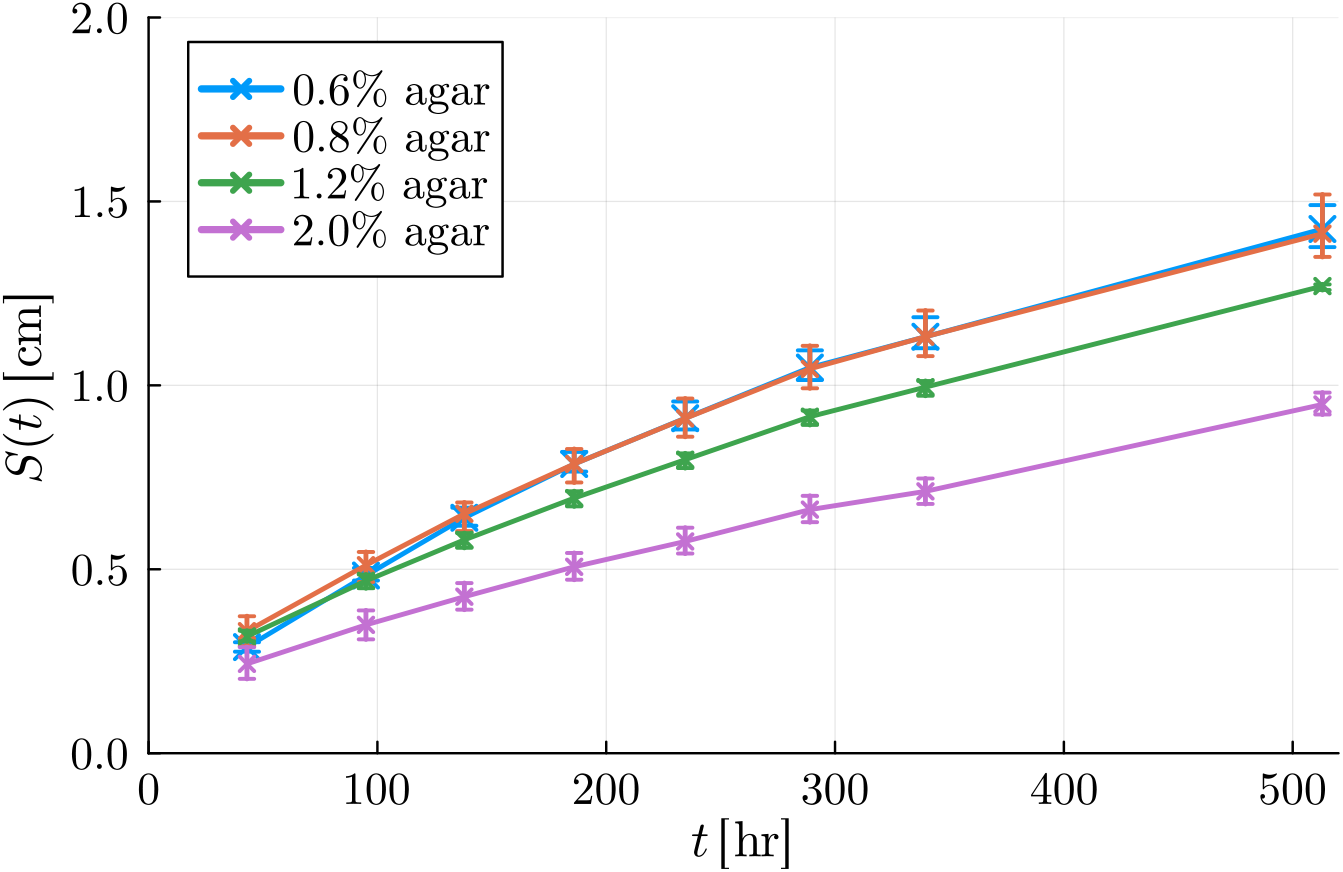
Mean experimental half-width, *S*(*t*), for colony biofilms grown on different density agar media, plotted against time. Error bars indicate the range observed across all experiments (*n* = 4 for all agar densities, except *n* = 3 for 2.0% agar).

Our experiments also provide measurements of the proportions of living and dead cells on Day 14, for one replicate on 0.6% agar and one replicate on 2.0% agar. Photographs of colonies stained with Phloxine B (Figure 1) suggest that the proportion of living cells is maximum at the leading edge, and decreases towards the centre of the colony biofilm. In sampling from a 2 cm^2^ region of agar across the full width of the colony biofilm, we found 89.5% total cell viability on low-density (0.6%) agar, and 89.9% on high-density (2.0%) agar. Since there was a small difference in overall viability between agar densities, we assumed that each colony biofilm has the same composition regardless of replicate and agar density. We then obtained smaller samples from the same two colonies at three localised regions of the colony biofilm, to quantify how cell viability depended on position. Averaging across both 0.6% and 2.0% agar, we obtained the results shown in Table 1. In previous experiments, some colony biofilms stained with Phloxine B included a red ring of elevated cell death near the rim, which could be due to regulated cell death [30]. However, we did not observe an analogous region in these experiments. Instead, the proportion of living cells is lowest in the centre of the colony biofilm, and increases towards the leading edge (Table 1).

**Table 1:** Mean proportion of living cells, Φ(*x*_*k*_), at three spatial locations along the colony biofilm. The proportions are averaged from one replicate each on 0.6% and 2.0% agar, at *t* = *t*_Φ_, which corresponds to Day 14.

| | **Centre** ($x_1 = 0$) | **Middle** ($x_2 = S(t_\Phi)/2$) | **Edge** ($x_3 = S(t_\Phi)$) |
| --- | --- | --- | --- |
| $\Phi(x_k)$ | 0.855 | 0.874 | 0.918 |

We used cross-sectional microscopy and cell-count data to estimate the colony-biofilm thickness. One limitation of our experimental method is that measuring the vertical profile was only possible after cutting the colony biofilm and imaging the cross-section. During cutting, the biofilm–substratum and biofilm–air interfaces of colonies grown on low-density agar changed shape significantly. Consequently, we could not obtain reliable thickness measurements for colony biofilms grown on low-density agar. Given these experimental challenges, we only measured the physical aspect ratio from micrographs of colony-biofilm cross-sections on high-density (2.0%) agar, such as in Figure 3. This yielded an aspect ratio of 0.0683 for 2.0% agar.

**Figure 3:**
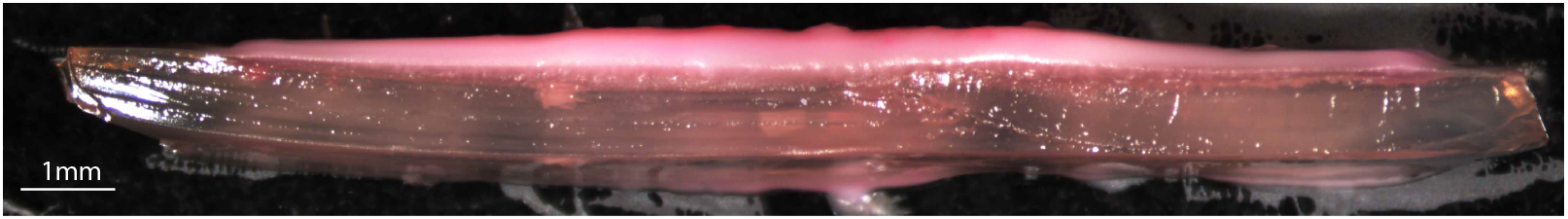
Cross-section of a colony biofilm grown on 2.0% agar, after 21 days of growth. We used this micrograph to measure the physical aspect ratio for 2.0% agar. We subsequently estimated the aspect ratio for 0.6%, 0.8%, and 1.2% agar using cell-count data.

We used total cell counts to estimate the aspect ratios for 0.6%, 0.8%, and 1.2% agar. Cell-count data were collected using a hemocytometer and a Nikon Eclipse 50i microscope on Day 21, providing an accurate measure of biomass volume. For each experiment, we used a 1 cm strip to count the mean number of cells per mm in the *y*-direction of the dish. The ratio of cell count to mean width then provided an estimate for the colony-biofilm thickness, in cells per unit area in the (*x, y*) plane. Provided that the cell sizes and colony-biofilm compositions are similar for each agar density, the ratio of thickness in cells mm^−2^ to half-width in mm will be approximately proportional to the physical aspect ratio. Consequently, we estimated the physical aspect ratio by scaling the cell-count aspect ratio by a factor chosen to ensure that the aspect ratio for 2.0% agar is the measured value, 0.0683. Table 2 shows these calculations, with the results showing that the quantitative trends are consistent with the observations that colonies are thicker on higher-density agar than on lower-density agar.

**Table 2:** Aspect-ratio calculations based on the experimental cell-count data from Day 21 on agar media of different density.

| Agar<br>Density, $a$<br>(%) | $n$ | Mean<br>Count<br>(cells mm<sup>-1</sup>) | Mean Half-Width<br>(mm) | Mean<br>Thickness<br>(cells mm<sup>-2</sup>) | Aspect<br>Ratio, $A(a)$ |
| --- | --- | --- | --- | --- | --- |
| 0.6 | 4 | $1.06 \times 10^8$ | 14.3 | $7.44 \times 10^7$ | 0.0306 |
| 0.8 | 4 | $1.40 \times 10^8$ | 14.1 | $9.91 \times 10^7$ | 0.0411 |
| 1.2 | 4 | $1.60 \times 10^8$ | 12.7 | $1.25 \times 10^8$ | 0.0577 |
| 2.0 | 3 | $1.12 \times 10^8$ | 9.5 | $1.14 \times 10^8$ | 0.0683 |

To summarise the experimental measurements and enable comparison with the model, we introduce the following notation:

- *S*_*i*_(*t*_*j*_; *a*): Experimental colony-biofilm half-width for replicate *i* = 1, 2, 3, 4 on agar density *a*, at time *t*_*j*_, for *j* = 1, …, 8, where *t*_*j*_ represents the time of the *j*-th photograph.
- Φ_*k*_ = Φ(*x*_*k*_): Living-cell volume fraction at spatial position *x*_*k*_, for *k* = 1, 2, 3, averaged over two replicates (one on 0.6% agar, and one on 2.0% agar). The three measurements were taken at approximately *x*_1_ = 0, *x*_2_ = *S*(*t*_Φ_)*/*2, and *x*_3_ = *S*(*t*_Φ_), where *t*_Φ_ corresponds to Day 14.
- *A*(*a*): Colony-biofilm aspect ratio on agar density *a*, averaged over all replicates on agar density *a*. The aspect ratio is defined as the colony biofilm’s maximum vertical thickness divided by its mean half-width. We obtained one estimate per experiment at time *t* = *t*_*A*_, corresponding to Day 21.

When comparing the model with experiments, we nondimensionalise the experimental quantities by scaling the experimental results in accordance with the model nondimensionalisation in Section 3.2.

## 3 Mathematical Model

We develop a thin-film continuum-mechanical model for colony-biofilm growth, similar to those used by several authors [18–20, 24]. The base model considers colony-biofilm growth along and perpendicular to a rigid substratum of width *X*_*s*_ and depth *H*_*s*_. The colony biofilm exists in the region Ω_*b*_(*t*) bounded by 0 < *x* < *S*(*t*), and 0 < *z* < *h*(*x, t*), where *S*(*t*) is the colony-biofilm half-width, *t* is time, and *z* = *h*(*x, t*) is the biofilm–air interface, which is a free surface. We model cellular material and the EPS matrix as viscous fluids [38], and partition the biomass into two phases: a living phase that contributes to growth and nutrient consumption, and an inactive phase of passive fluid. To incorporate both phases into the model, we introduce the volume fractions [39] of the living phase, *ϕ*_*n*_(*x, z, t*), and the inactive phase, *ϕ*_*m*_(*x, z, t*). The living biomass accounts for proliferating cells, and the inactive biomass phase represents dead cells. Since the proportion of cells to ECM in our experiments is unknown, we assume that both phases also contain extracellular fluid, such that the amount of ECM in each phase is proportional to the volume of cellular material. The usual no-voids assumption, *ϕ*_*n*_ + *ϕ*_*m*_ ≡ 1, applies throughout the domain.

The colony biofilm expands by cell proliferation and ECM production, driven by the consumption of nutrients supplied from the substratum. We assume that the nutrients occupy no volume and denote the nutrient concentration in the substratum as *g*_*s*_(*x, z, t*). This variable is defined in the substratum domain Ω_*s*_, which is the fixed region bounded by 0 < *x* < *X*_*s*_ and −*H*_*s*_ < *z* < 0. During growth, the colony biofilm takes up nutrients from the substratum through the biofilm–substratum interface. We denote the nutrient concentration in the colony biofilm as *g*_*b*_(*x, z, t*), defined in Ω_*b*_(*t*). A schematic illustrating the model geometry is shown in Figure 4.

**Figure 4:**
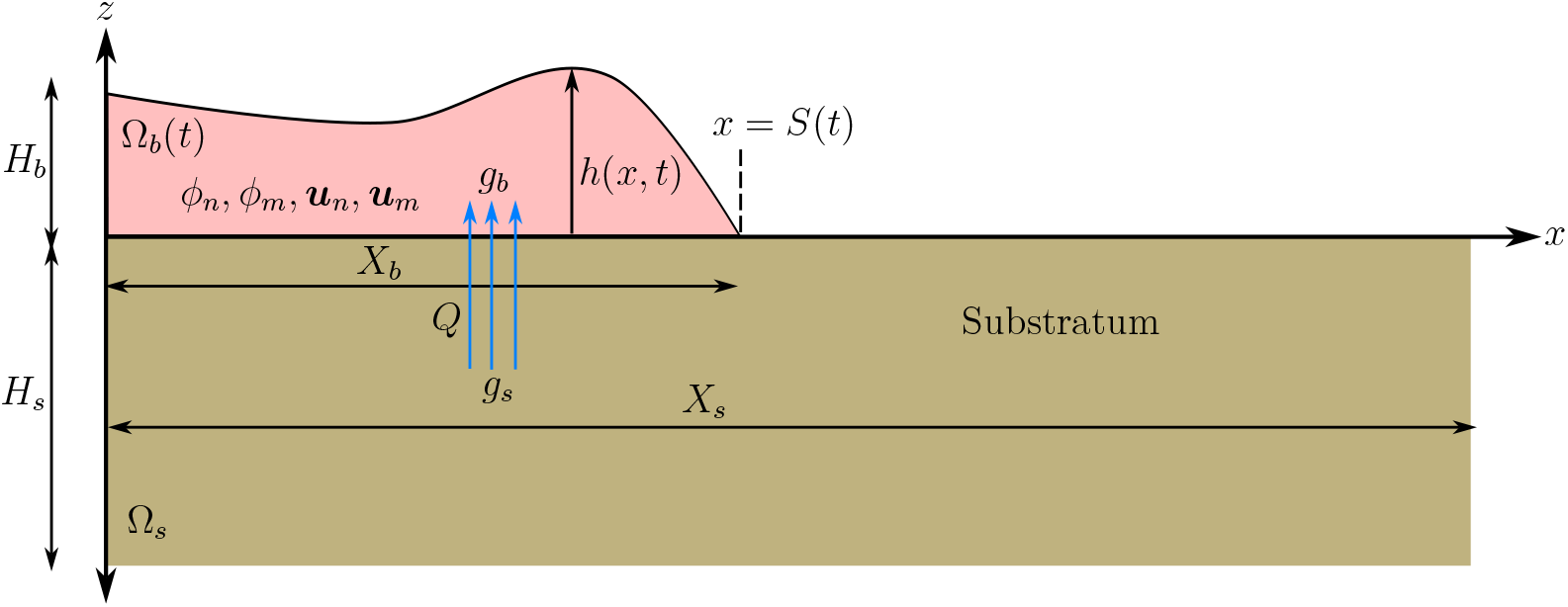
Schematic illustrating the geometry, variables, and parameters of the mathematical model.

### 3.1 Governing Equations and Boundary Conditions

The governing equations arise from mass and momentum conservation, and their derivation closely follows Tam et al. [19]. We summarise the key assumptions here, and present a detailed derivation in the Supporting Material. Yeast cells are non-motile. Unlike bacteria [40, 41], yeasts do not respond actively to environmental cues, such as agar density. Instead, colony-biofilm expansion occurs due to cell proliferation facilitated by catabolism of glucose [42]. We adopt first-order kinetics for nutrient consumption and biomass production. We assume that cell death occurs at a rate proportional to the volume fraction of living cells, which represents random accidental cell death. On death, we assume that cells immediately form part of the inactive phase, without any corresponding change in volume [18]. This assumption is consistent with the hypothesis that cell death influences the ECM composition [28]. We assume that nutrients disperse by diffusion in the substratum, and by diffusion and advection with both fluid phases in the colony biofilm, where they become available for consumption.

Following these model assumptions, we obtain mass-balance equations for the volume fractions and nutrient concentrations:

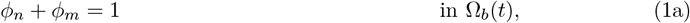

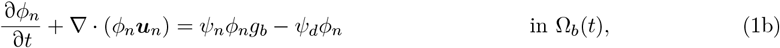

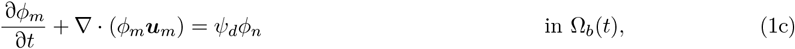

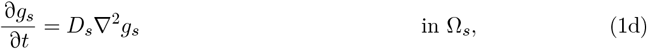

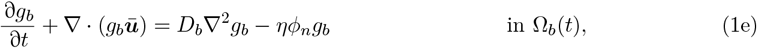

where ***ū*** = *ϕ*_*n*_***u***_*n*_ + *ϕ*_*m*_***u***_*m*_ is the mixture-averaged fluid velocity, ∇ = ( *∂*_*x*_, *∂*_*z*_), the parameter *ψ*_*n*_ is the biomass-production rate, *ψ*_*d*_ is the cell-death rate, *D*_*s*_ and *D*_*b*_ are the diffusivities of nutrients in the substratum and colony biofilm, respectively, and *η* is the nutrient-consumption rate.

The colony-biofilm rheology is difficult to determine precisely. Since the cells and the extracellular fluid are both primarily composed of water, for simplicity we assume that both phases are incompressible viscous Newtonian fluids [19]. Conservation of momentum then yields

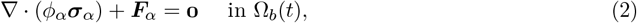

Where

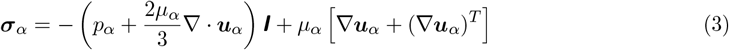

is the Cauchy stress tensor, ***F***_*α*_ represents external sources of momentum for phase *α* = *n, m*, and ***I*** is the identity matrix. For both fluid phases we neglect an explicit contribution to the pressure due to growth, instead assuming that material incompressibility drives expansion in response to cell proliferation [18, 19]. Owing to the mass source terms that represent cell proliferation, cell death, and ECM production, the velocity field is in general not divergence free. Thus, we need to retain a bulk viscosity term in the stress tensor (3). We choose the coefficient of this term, −2*µ*_*α*_*/*3, in accordance with Stokes’ hypothesis [18, 19, 43, 44]. The momentum sources, ***F***_*α*_, consist of interphase drag and interfacial forces [19, 45–47]. These momentum sources are given explicitly by ***F***_*α*_ = −*k*(***u***_*n*_ − ***u***_*m*_) + *p*_*α*_∇*ϕ*_*α*_ for *α* = *n, m*, where *k*(*ϕ*_*n*_, *ϕ*_*m*_) is the interphase drag coefficient. Since the interphase drag is large on the length scales relevant to biofilm growth [48–50], we assume that both fluid phases move with the common velocity ***u***_*n*_ = ***u***_*m*_ = ***u*** = (*u, w*). To simplify the model further, we assume that both phases share a common pressure *p*_*n*_ = *p*_*m*_*p* and dynamic viscosity *µ*_*n*_ = *µ*_*m*_ = *µ* [19].

We impose no-flux conditions for *g*_*s*_ at the Petri-dish boundaries *x* = *X*_*s*_ and *z* = −*H*_*s*_, and a no-flux condition for *g*_*b*_ on the biofilm–air interface *z* = *h*(*x, t*). We obtain another boundary condition by assuming that nutrients enter the colony biofilm at a rate proportional to the local concentration difference across the biofilm–substratum interface, neglecting osmotic effects. This yields the conditions

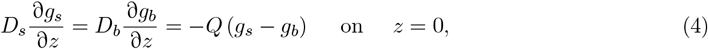

where *Q* is a mass-transfer coefficient describing the rate of nutrient uptake. We also apply standard boundary conditions from fluid mechanics, including no penetration on the biofilm–agar interface, and a kinematic condition as well as tangential and normal stress conditions on the biofilm–air free surface. In our model, surface tension might represent the typical fluid surface tension, or the strength of cell–cell adhesion at the biofilm–air interface [51]. Since sliding motility involves weak cell–cell adhesive forces [12, 13], we neglect surface tension in the normal stress condition on the free surface. In the work of Tam et al. [19], surface tension had minimal impact on horizontal expansion. The surface-tension coefficient does affect the colony-biofilm shape, and can inhibit ridge formation on the colony-biofilm surface. Our experimental results do not contain ridges, and have similar profiles to the solutions of Tam et al. [19] with zero surface tension. Hence, we neglect surface tension in this work.

For the tangential stress balance on the biofilm–substratum interface, we impose a general slip condition [47, 52], leading to

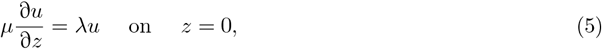

where *λ* is a slip parameter representing the strength of biofilm–substratum adhesion, and is assumed constant. Whilst the general tangential stress condition (5) was included in the models of Tam et al. [19] and Tam et al. [20], those works focused on the weak adhesion and perfect slip (*λ* = 0), and strong adhesion and no slip (*λ* → ∞) scenarios, respectively. In this work, we consider the generalised case in which *λ* varies between these two limits, and estimate the value of *λ* using experimental data. The full dimensional model then consists of the governing equations (where all PDEs apply in Ω_*b*_(*t*), except for equation (6c) which applies in Ω_*s*_)

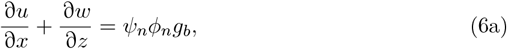

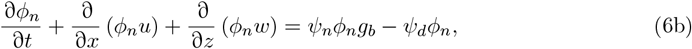

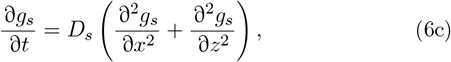

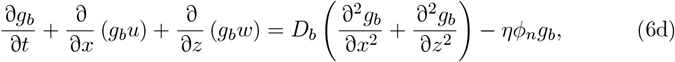

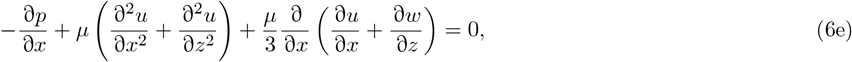

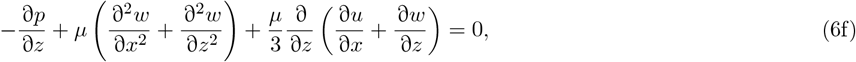

the boundary conditions

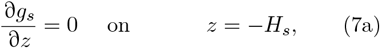

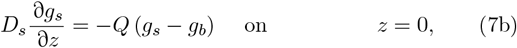

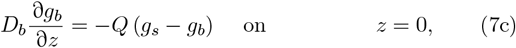

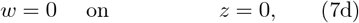

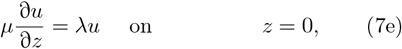

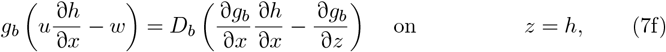

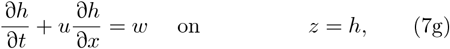

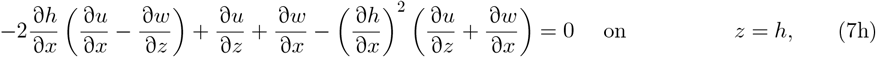

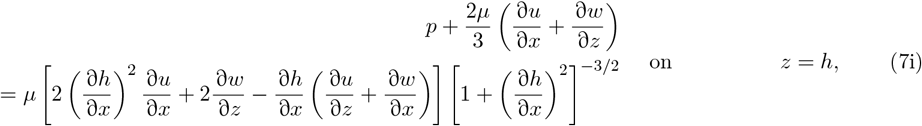

and initial conditions, which are specified in Section 3.3.

### 3.2 Extensional-Flow Scaling and Nondimensionalisation

After establishing the mass and momentum balances, we simplify the governing equations using the thin-film approximation. We assume that the aspect ratios of the colony biofilm and agar substratum are equal, and exploit their thin geometries by writing

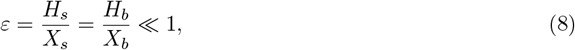

where *H*_*b*_ and *X*_*b*_ are the characteristic thickness and characteristic half-width of the colony biofilm, respectively. The characteristic colony-biofilm thickness is chosen to be *H*_*b*_ = *X*_*b*_*H*_*s*_*/X*_*s*_, which is not necessarily the initial colony biofilm thickness. This assumption is valid because the aspect ratio of the substratum in the experiments, *ε* = *H*_*s*_*/X*_*s*_ ≈ 0.06 ≪ 1, lies within the aspect-ratio range of mature colony biofilms (between 0.032 and 0.068 depending on agar density).

We nondimensionalise the governing system by scaling variables as follows

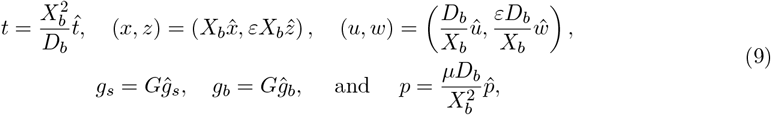

where *G* is the initial nutrient concentration in the substratum. As the thin-film approximation in Section 3.3 will outline, our scaling leads to an extensional flow [18, 53], where the nutrient concentrations and the horizontal component of the fluid velocity are independent of the depth *z* to leading order in *ε*. The pressure scaling in the extensional flow regime differs from the common lubrication approximation which scales pressure as 1*/ε*^2^ [18]. Surface tension [13] and biofilm–agar friction [7, 12] are low in colony biofilms, whereas the typical lubrication scaling is associated with high surface tension and no slip. The extensional-flow regime also produces more realistic colony-biofilm shapes than the lubrication regime [19, 20], which assumes a comparatively large pressure difference across the biofilm–air interface, balanced by a larger surface tension.

After scaling, the dimensionless governing equations become (dropping hats)

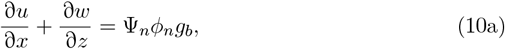

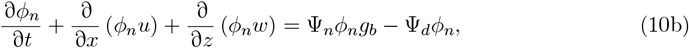

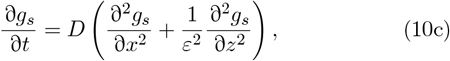

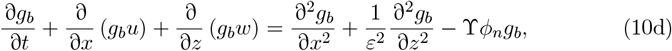

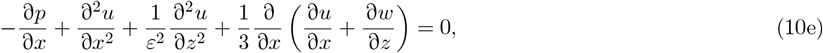

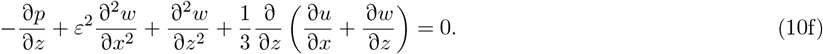

The dimensionless boundary conditions are

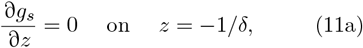

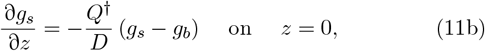

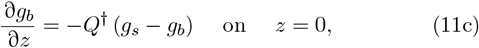

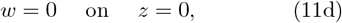

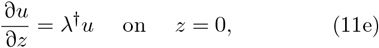

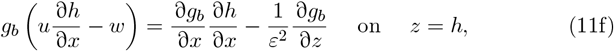

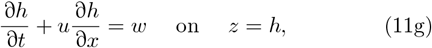

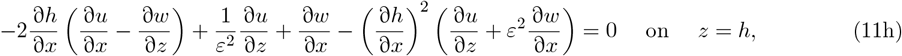

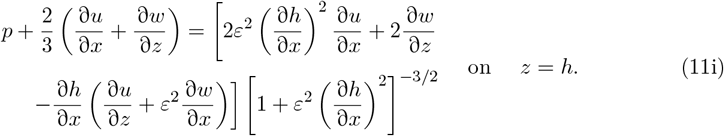

The dimensionless model (10) and (11) contains seven distinct dimensionless parameters,

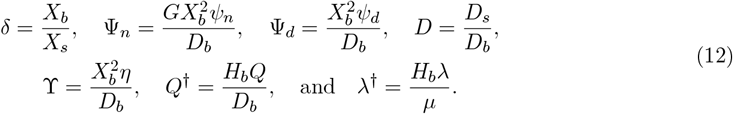

The parameter *δ* is the ratio of the colony-biofilm length scale to the Petri-dish length scale. The parameters Ψ_*n*_ and Ψ_*d*_ are the dimensionless biomass-production rate and cell-death rate, respectively, *D* is the nutrient-diffusivity ratio, and ϒ is the nutrient-consumption rate. The parameter *Q*^†^ is the nutrient-uptake rate by the colony biofilm, and *λ*^†^ is the dimensionless biofilm–substratum adhesion strength. In this work, we consider the distinguished limit in which *Q*^†^ and *λ*^†^ are of 𝒪(*ε*^2^) size as *ε* → 0, under the assumption that they themselves are small, but contribute to effects that are non-negligible. Accordingly, we rescale *Q*^†^ and *λ*^†^ by setting *Q*^†^ ∼ *ε*^2^*Q*^*^ and *λ*^†^ ∼ *ε*^2^*λ*^*^, where *Q*^*^ and *λ*^*^ are 𝒪(1) quantities as *ε* → 0. Nutrient uptake and slip then enter the leading-order model through 𝒪(*ε*^2^) corrections to the boundary conditions. This regime represents weak biofilm–substratum adhesion, which is suitable for expansion by sliding motility [19]. Our parameter estimation (see Section 4.1) will predict that *Q*^*^, *λ*^*^, and the other dimensionless quantities in (12) are of 𝒪(1) size, indicating that this scaling regime is suitable.

### 3.3 Thin-Film Approximation

We apply the thin-film approximation by expanding variables in power series of *ε*^2^,

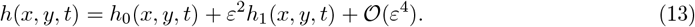

Substituting (13) into the model (10) and (11) yields a problem where the nutrient concentrations and the horizontal component of the velocity are independent of *z* to leading order in *ε*. We use the no-voids condition (1a) to eliminate *ϕ*_*m*_ from the system. It is also natural to introduce the depth-averaged cell volume fraction,

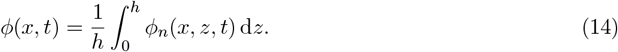

in order to integrate out the *z*-dependence in *ϕ*_*n*_, such that all of the dependent variables are functions of *x* and *t* to leading order in *ε*.

The spatially one-dimensional system to leading order in *ε* is (dropping the zero subscript on leading-order quantities)

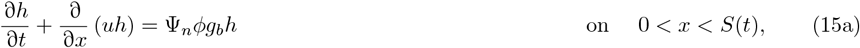

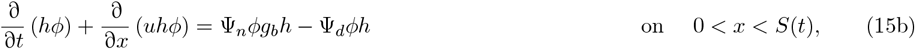

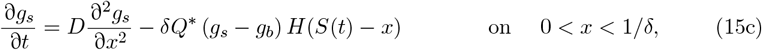

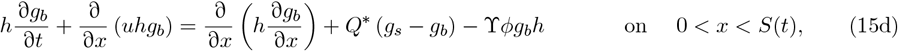

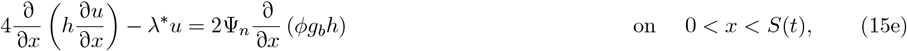

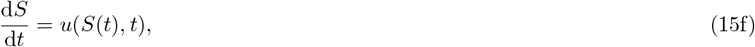

where *H*(*x*) is the Heaviside step function. We include the Heaviside step function in (15c) because nutrient uptake only occurs where the colony biofilm is present. The parameter *δ* influences nutrient depletion from the substratum through equation (15c), such that nutrients deplete more slowly on comparatively large substrates. The slip parameter *λ*^*^ provides a drag-like term in the leading-order momentum equation (15e), such that increasing *λ*^*^ represents stronger biofilm–substratum adhesion, inhibiting expansion.

To close the model, we impose the initial conditions

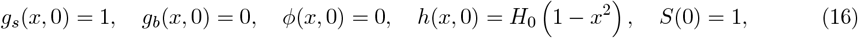

where *H*_0_ is the dimensionless initial colony-biofilm thickness. The boundary conditions are

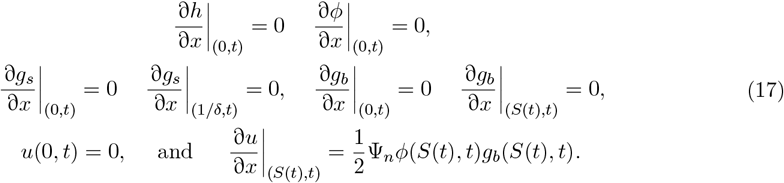

The condition for *u* at *x* = *S*(*t*) arises by enforcing zero normal stress at *x* = *S*(*t*). A complete derivation of the system (15), (16) and (17) is available in the Supporting Material. An advantage of our two-phase moving-boundary model is that it enables comparison with new experimental data for the colony biofilm composition, expansion speed, and aspect ratio. We compute numerical solutions of the model (15), (16) and (17), and compare the results with our experiments. We obtain numerical solutions to the mathematical model (15), (16) and (17) using a front-fixing transformation and the Crank–Nicolson method, similar to Tam et al. [19]. Full details of the numerical scheme are available in the Supporting Material, and open source Julia code is available on GitHub.

### 3.4 Model Parameters

Table 3 contains parameter values used throughout this work. The agar density, agar depth, amount of nutrient, duration, and Petri-dish dimensions are known from the experiments. We also obtain estimates for the nutrient diffusivities from the literature, yielding estimates for *D*_*s*_ [54, 55] and *D*_*b*_ [56]. Of the known parameters, only the nutrient diffusivity in the substratum, *D*_*s*_, depends on agar density. In practice, the nutrient diffusivity in agar will depend on physical properties of the agar, such as the porosity of its microstructure [57, 58]. Whilst we do not measure porosity directly in our experiments, the effects of porosity can be interpreted using measured relationships between agar density and porosity [59]. In our work, for simplicity we use the empirical relationship between agar density and nutrient diffusivity of Slade, Cremers, and Thomas [55]. This relationship suggests that *D*_*s*_ depends weakly on agar density, and is similar to the diffusivity of nutrients in water.

**Table 3:** Estimated values for parameters in the mathematical model.

| Parameter | Symbol | Value(s) | Units | Source(s) |
| --- | --- | --- | --- | --- |
| Agar density (weight percentage) | $a$ | $\{0.6, 0.8, 1.2, 2.0\}$ | – | Experiment |
| Experimental duration | $T$ | 30780 | min | Experiment |
| Petri dish half-width | $X_s$ | 50 | mm | Experiment |
| Agar depth | $H_s$ | 3 | mm | Experiment |
| Initial colony-biofilm half-width | $X_b$ | 2 | mm | Experiment |
| Initial colony-biofilm thickness | $H_0$ | 0.05 | – | [61–63] |
| Initial nutrient concentration | $G$ | $1.5 \times 10^{-4}$ | $\text{g mm}^{-2}$ | Experiment |
| Nutrient diffusivity (water) | $D_0$ | $4.04 \times 10^{-2}$ | $\text{mm}^2 \text{min}^{-1}$ | [54] |
| Nutrient diffusivity (substratum) | $D_s$ | $(1 - 0.023a)D_0$ | $\text{mm}^2 \text{min}^{-1}$ | [55] |
| Nutrient diffusivity (biofilm) | $D_b$ | $0.24D_0$ | $\text{mm}^2 \text{min}^{-1}$ | [56] |

In the extensional-flow regime, the colony-biofilm expansion speed depends on the initial conditions [18, 19]. If the nutrient supply and the cell volume fraction are held constant, Ward and King [18] showed that the colony-biofilm profile evolves to a travelling wave with speed proportional to 1*/H*_0_ as *t* → ∞, where *H*_0_ is the initial dimensionless colony-biofilm thickness. Consequently, *H*_0_ will impact expansion speed, and subsequently the comparison between the model and experiments. Although we have no reliable way to measure *H*_0_ in our experiments, all experiments were inoculated in the same way.

Therefore, we assume that *H*_0_ is constant across all replicates, and independent of agar density. In the experiments, cells were applied to the agar using the tip of an inoculating loop of 4.5mm diameter, such that *X*_*b*_ ≈ 2 mm. The inoculum is a thin streak of cells and fluid that is rapidly absorbed into the agar layer. The cell density of this initial streak is significantly less than the cell density once the colony biofilm develops [60]. Consequently, we assume an initial dimensional thickness of 6 µm, which is an upper estimate for the minor axis diameter of a single cell of Σ1278b *S. cerevisiae* yeast [61–63]. This assumption yields the dimensionless initial thickness *H*_0_ = 0.006*X*_*s*_*/*(*H*_*s*_*X*_*b*_) = 0.05. This small value for *H*_0_ reflects that the initial colony-biofilm thickness in experiments is much smaller than the thickness of a mature colony biofilm [8].

We still require values for the biomass-production rate Ψ_*n*_, cell-death rate Ψ_*d*_, nutrient-uptake coefficient *Q*^*^, nutrient-consumption rate ϒ, and the slip coefficient *λ*^*^. Since cellular behaviour and/or properties of the substratum determine these parameters, they are difficult to measure directly in experiments. We also hypothesise that these parameters may depend on the agar density. Consequently, we cannot adopt the parameter values used by Tam et al. [19], whose experiments were performed solely on 0.3% agar. Since we cannot estimate these five parameters (Ψ_*n*_, Ψ_*d*_, *Q*^*^, ϒ, *λ*^*^) from the experiments or the existing literature, we obtain parameter estimates by fitting the model to new experimental data. Differences in these parameters between experiments on different agar densities will indicate how the agar density impacts colony-biofilm growth.

## 4 Results and Discussion

We use numerical solutions and optimisation to fit five unknown parameters to experimental data on different density agar. We obtain good fits to expansion speed, colony-biofilm composition, and aspect-ratio data. This approach reveals a stronger impact of nutrient limitation on lower-density agar, and increased biofilm–substratum adhesion on higher-density agar. A combination of sensitivity analysis and synthetic data suggests that increased biofilm–substratum adhesion is the most robust effect of increasing agar density.

### 4.1 Parameter Estimation

We use a numerical optimisation method to obtain point estimates for the model parameters that correspond to experiments. The quantities *X*_*b*_, *G, D*_*b*_, *ε*, and *µ* are independent of the agar density, and we seek one set of optimal parameters *θ*^*^(*a*) = (Ψ_*n*_(*a*), Ψ_*d*_(*a*), *Q*^*^(*a*), ϒ(*a*), *λ*^*^(*a*)) for each of the four agar densities used in our experiments. Our parameter estimates are based on minimising the difference between the experimental data and a numerical solution to the mathematical model. To perform the comparison, we introduce three summary statistics, 𝒮_*S*_, 𝒮_Φ_, and 𝒮_*A*_. These statistics are the *L*^2^-norms of the relative differences in half-width, composition, and aspect ratio, respectively, between the model solution with parameters *θ* and the experiments on agar density *a*. Their definitions are

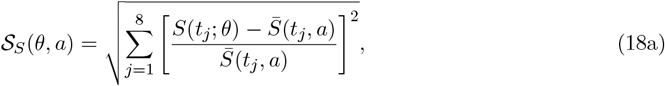

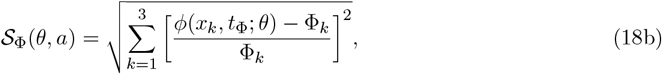

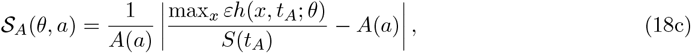

Where

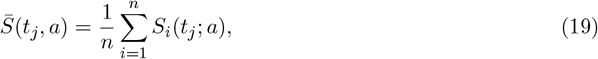

is the half-width averaged over the *n* experimental replicates on agar density *a*. Based on these summary statistics, we define the distance metric

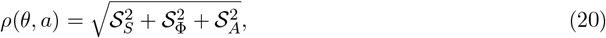

which provides a single measure of the difference between the model and experiments. Using the relative difference for each statistic (18) means that the half-width, composition, and aspect ratio will have similar weighting when calculating *ρ*. The optimal parameter values for agar density *a* are then

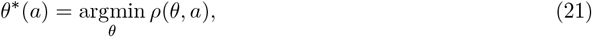

which is the set of parameters that minimises the distance between the model solution and experimental data.

To establish a procedure for computing the optimal parameter set *θ*^*^(*a*) numerically, we use optimisation packages available in Julia. For a given agar density, the optimisation procedure is as follows:

1. We obtain a coarse estimate, 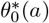, using the Differential Evolution method with adaptive weights [64, 65], as implemented in the global optimisation package `BlackBoxOptim.jl`. We terminate the black-box optimisation after 100 function evaluations, where one function evaluation refers to a computation of *ρ*(*θ, a*), which involves solving the PDE model (15), (16) and (17) for a set of parameters *θ*, and computing 𝒮_*S*_, 𝒮_Φ_, and 𝒮_*A*_. The black-box step provides a good initial guess for gradient-based optimisation. Increasing the number of function evaluations beyond 100 had a negligible effect on the results.
2. Using 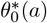 as the initial guess, we use the LBFGS [66–69] method in Optim.jl [70] to estimate *θ*^*^(*a*). Where necessary, we approximate the gradient ∇_*θ*_*ρ* numerically using first-order finite differencing, and use the Hager–Zhang line search [71]. We apply box constraints to ensure that all parameters are positive, with 3 outer iterations, and absolute tolerances of *ε* = 1 × 10^−6^ for the objective function and the *L*^∞^-norm of the minimiser.

Repeated application of the optimisation procedure to the same experimental data yielded only minor differences in the minimiser *θ*^*^(*a*). Therefore, on each run our gradient-based optimisation method converged effectively to a global minimum in *ρ*. Since each evaluation of the objective function *ρ*(*θ, a*) requires solving the model numerically once, the parameter estimation procedure is computationally expensive. In this work, we present all results using *N*_*ξ*_ = 101 grid points and *N*_*τ*_ = 401 for consistency, as the computational cost for large-scale results becomes prohibitive for finer resolutions. Although the quantitative results at this resolution are grid dependent, the qualitative conclusions reported herein remain valid as the numerical grid is refined. Further information regarding grid convergence is available in the Supporting Material. The optimal sets of dimensionless parameters are presented in Table 4. We obtained a good fit to experimental data for all agar densities considered. The accuracy of the fit is illustrated in Figure 5, where the colony-biofilm size over time predicted numerically closely matches the experiments.

**Table 4:** Numerical solutions for the optimal set of dimensionless parameters, *θ*^*^(*a*), for each agar density, *a*. The presented parameter values are the mean values from 5 repetitions of the optimisation routine, with the numerical solutions having been computed using *N*_*ξ*_ = 101 grid points and *N*_*τ*_ = 401 time steps.

| $a$ | $\Psi_n$ | $\Psi_d$ | $Q^*$ | $\Upsilon$ | $\lambda^*$ | $\rho(\theta^*, a)$ |
| --- | --- | --- | --- | --- | --- | --- |
| 0.6 | 0.301 | 0.00777 | 7.56 | 7.87 | 0.670 | 0.0437 |
| 0.8 | 0.365 | 0.00747 | 6.55 | 7.33 | 1.041 | 0.0536 |
| 1.2 | 0.409 | 0.00755 | 4.48 | 6.46 | 1.848 | 0.0636 |
| 2.0 | 0.302 | 0.00835 | 3.27 | 5.01 | 2.671 | 0.0625 |

**Figure 5:**
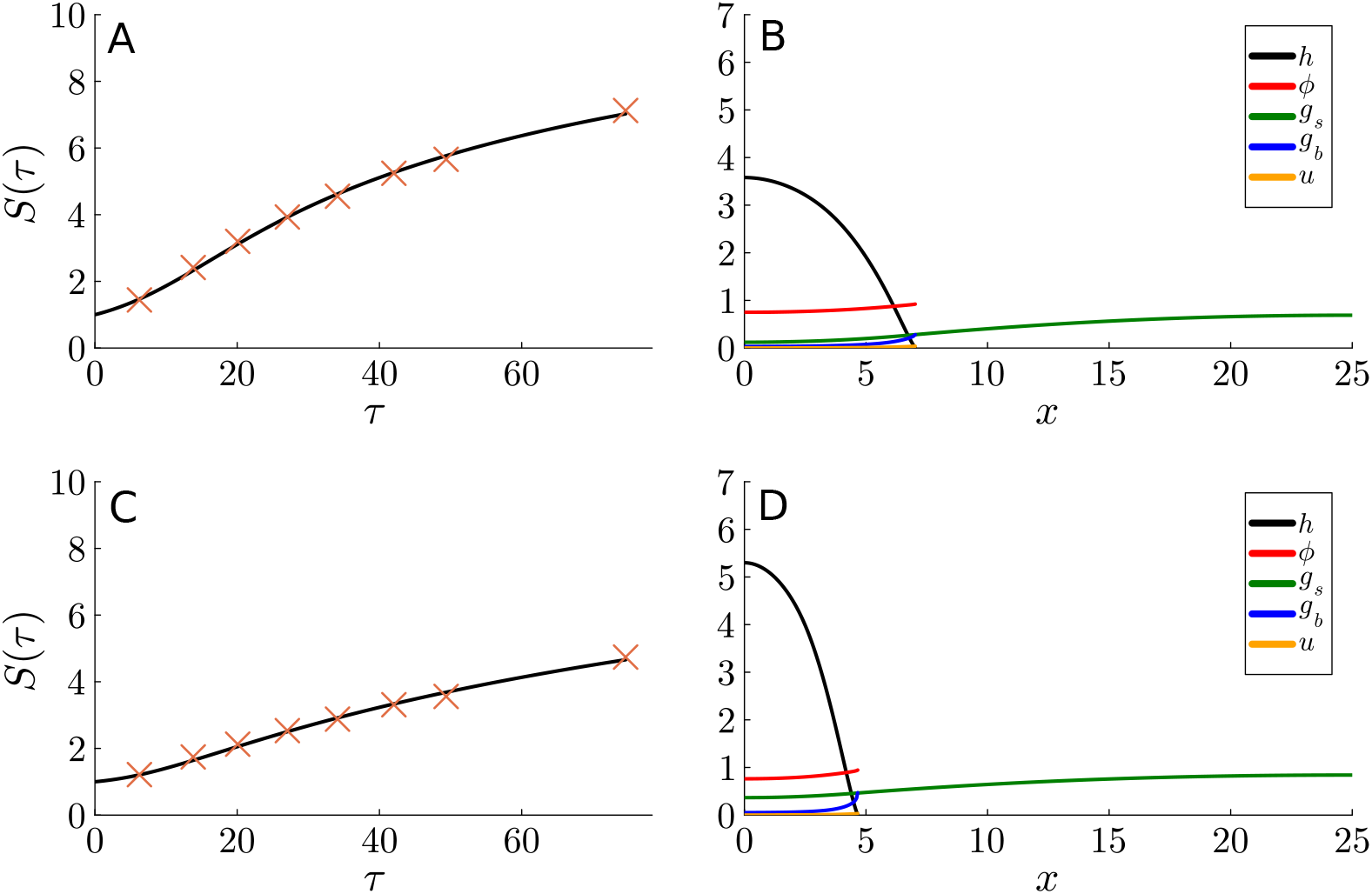
Numerical solutions to the mathematical model (15), (16) and (17) with the estimated parameters from Table 4, for both low-density (0.6%) and high-density (2.0%) agar. The numerical solutions were computed using *N*_*ξ*_ = 101 grid points and *N*_*τ*_ = 401 time steps. (A) Comparison between the model solution (black curve) and experimental data (orange crosses) for the dimensionless colony-biofilm size, *S*(*τ* ), on 0.6% agar. (B) Numerical solution for *τ* = 74.6, corresponding to Day 21 of the experiment, using the optimal parameters for 0.6% agar. (C) Comparison between the model solution (black curve) and experimental data (orange crosses) for the dimensionless colony-biofilm size, *S*(*τ* ), on 2.0% agar. (D) Numerical solution for *τ* = 74.6, corresponding to Day 21 of the experiment, using the optimal parameters for 2.0% agar.

The optimal biomass-production rate, Ψ_*n*_, and cell-death rate, Ψ_*d*_, only vary slightly with changing agar density. These results are consistent with the idea that yeast cells do not actively alter their proliferation or death rates in response to environmental cues, such as agar density. In contrast, *Q*^*^, ϒ, and *λ*^*^ vary more significantly with agar density. The nutrient-uptake rate, *Q*^*^, decreases with increasing agar density, which accords with the observations of *E. coli* bacterial colonies by He et al. [35]. Like Ψ_*n*_ and Ψ_*d*_, the nutrient-consumption rate, ϒ, is a property of the yeast cells rather than agar density. Consequently, ϒ varies less than *Q*^*^ as agar density changes. The decrease in both *Q*^*^ and ϒ with agar density suggests that nutrients deplete faster for colonies grown on lower-density agar. This depletion could also be influenced by agar dehydration or osmotic effects, which we have neglected in the model.

The most striking effect of increasing agar density is the increase in the dimensionless adhesion strength, *λ*^*^. The small value of *λ*^*^ on 0.6% agar suggests that the perfect slip approximation (*λ*^*^ = 0) may be suitable for very low-density agar. This supports the use of perfect slip by Tam et al. [19], whose experiments were on 0.3% media. The inclusion of non-zero *λ*^*^ becomes more important on higher-density agar. The increased value of *λ*^*^ observed on higher-density substrates increases drag-like friction, inhibiting horizontal expansion [72, 73] relative to vertical expansion. This explains the thicker colony-biofilm profiles on higher-density agar. Similar behaviour in *E. coli* was also reported by He et al. [35], and this prediction aligns with the recent experimental results of Pokhrel et al. [32]. Given the potential for biological variability across experiments, we perform a series of analyses to assess the robustness of the results to changes in each parameter.

### 4.2 Uncertainty and Sensitivity Analyses

Our experimental results consist of relatively few replicates that are subject to biological variation. We perform a series of analyses to determine the robustness of the parameter-estimation results presented in Section 4.1. First, we investigate how varying the model parameters influences *ρ*(*θ, a*), the global distance metric that compares the mathematical model solution and experimental data. We present these results in the form of parameter-pair heat maps for 0.6% agar in Figure 6. We vary parameters two at a time from zero to approximately three times their optimal values, holding the other parameters constant at their optimal values. The parameter-pair heat maps show the computed values of *ρ*(*θ, a*) as two parameters are varied.

**Figure 6:**
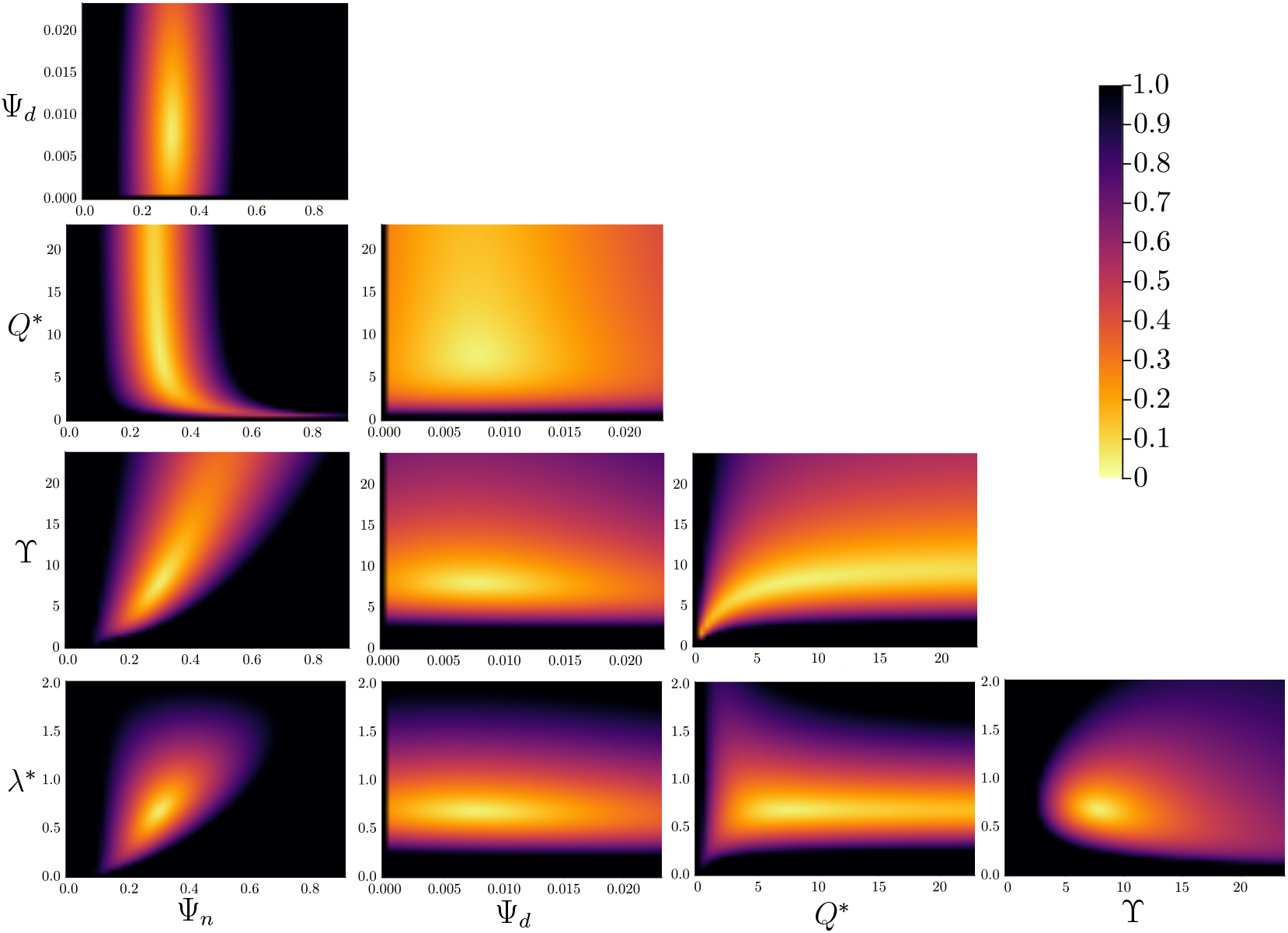
Parameter-pair heat maps for low-density (0.6%) agar. Each plot shows the value of the global distance metric, *ρ*(*θ, a*), for a given set of parameters. Unless otherwise stated, all parameters take the optimal values listed in Table 4. When varied, each parameter ranges from zero to three times the optimal value (2,500 total simulations).

The biomass-production rate, Ψ_*n*_, has a narrow band of values that yield small values of *ρ*(*θ, a*). Increasing Ψ_*n*_ increases *S*(*t*) in the numerical solutions [19]; varying Ψ_*n*_ from its optimal value causes large changes in *ρ*, suggesting that Ψ_*n*_ strongly influences the solutions and is well-identified by the experimental data. Compared to Ψ_*n*_, varying Ψ_*d*_ has a smaller impact on the size of *ρ*. This is because the cell-death rate has a less significant effect on expansion speed [19], and only significantly impacts the living-cell volume fraction, *ϕ*. Since colony-biofilm composition had minimal variation with agar density in our experiments, the optimal value for Ψ_*d*_ remains similar across all agar densities considered.

Compared to other parameters, the distance *ρ*(*θ, a*) is also less sensitive to the nutrient-uptake rate, *Q*^*^. In the model, nutrients taken up from the substratum do not occupy volume, and therefore only contribute to growth when consumed. Nutrient consumption in the colony biofilm is influenced by both the uptake rate, *Q*^*^, and consumption rate, ϒ. Consequently, there is a wide range of different values for the pair (*Q*^*^, ϒ) that yield similarly good fits to the data. Therefore, although our procedure identifies differences in *Q*^*^ and ϒ across agar densities, the parameter *Q*^*^ is less well-identified than Ψ_*n*_. A future version of the model that accounts for volumetric increase due to nutrient uptake, for example by osmotic swelling, could shed further light on the importance of nutrient uptake.

Increasing the biofilm–substratum adhesion strength *λ*^*^ increases the aspect ratio and decreases horizontal expansion [20]. Consequently, this parameter is well identified, with a comparatively narrower range of values yielding small *ρ*(*θ, a*). Since the aspect ratio increases as the agar density increases, so too does the optimal adhesion strength *λ*^*^. Taken together, this parameter-pair analysis reveals the key roles of the biomass-production rate, nutrient consumption, and cell–substratum adhesion in determining the colony-biofilm size and shape.

To investigate the effects of biological variability on our parameter estimates, in Table 5 we infer optimal parameter sets from individual experimental replicates, rather than from the mean of all experiments performed on a given agar density. Although the optimal parameters vary across replicates, the quantitative trends persist across different replicates of the same agar density. To supplement these results, we repeated the optimisation procedure using synthetic data. A synthetic experiment refers to one set of synthetic data for the half-width *S*_*i*_(*t*_*j*_; *a*), the living-cell volume fraction Φ_*k*_(*x*_*k*_), and the aspect ratio *A*(*a*). We generated synthetic data based on the variability in each quantity observed across experimental replicates. For the synthetic aspect-ratio data, we generated a synthetic cell count from a normal distribution with the mean and standard deviation from experiments, and computed the corresponding aspect ratio following the method outlined in Section 2.2. To generate synthetic half-width data *S*(*t*_*j*_; *a*), we sampled once from the standard normal distribution, *z* ∼ 𝒩 (0, 1), and set 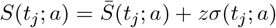 where 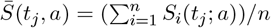 are the mean experimental measurements at each time point, and *σ*(*t*_*j*_; *a*) is the standard deviation. For the synthetic colony-biofilm composition data, we scaled the average experimental value of Φ_*k*_ by the same random number drawn from a uniform distribution *U* (0.932, 1.05), which allows for composition results across the range observed in experiments. We generated 50 synthetic experiments for each agar density, and obtained an optimal parameter set for each synthetic experiment.

**Table 5:** Optimal parameters for all individual experimental replicates.

| $a$ | Rep. | $\Psi_n$ | $\Psi_d$ | $Q^*$ | $\Upsilon$ | $\lambda^*$ | $\rho(\theta^*, a)$ |
| --- | --- | --- | --- | --- | --- | --- | --- |
| 0.6 | 1 | 0.299 | 0.00778 | 7.68 | 7.82 | 0.666 | 0.0446 |
|  | 2 | 0.306 | 0.00778 | 6.84 | 8.01 | 0.691 | 0.0460 |
|  | 3 | 0.312 | 0.00771 | 7.58 | 7.63 | 0.653 | 0.0448 |
|  | 4 | 0.290 | 0.00780 | 8.10 | 8.02 | 0.674 | 0.0405 |
| 0.8 | 1 | 0.363 | 0.00762 | 5.94 | 6.35 | 0.973 | 0.0537 |
|  | 2 | 0.312 | 0.00787 | 6.99 | 6.51 | 0.982 | 0.0451 |
|  | 3 | 0.378 | 0.00731 | 6.84 | 8.27 | 1.107 | 0.0555 |
|  | 4 | 0.416 | 0.00709 | 6.89 | 8.64 | 1.119 | 0.0590 |
| 1.2 | 1 | 0.380 | 0.00783 | 3.89 | 5.67 | 1.806 | 0.0629 |
|  | 2 | 0.392 | 0.00766 | 4.50 | 6.09 | 1.785 | 0.0619 |
|  | 3 | 0.429 | 0.00743 | 4.54 | 6.89 | 1.904 | 0.0652 |
|  | 4 | 0.434 | 0.00731 | 5.13 | 7.21 | 1.883 | 0.0640 |
| 2.0 | 1 | 0.301 | 0.00835 | 3.26 | 5.01 | 2.681 | 0.0620 |
|  | 2 | 0.236 | 0.00919 | 3.27 | 5.62 | 2.357 | 0.0510 |
|  | 3 | 0.404 | 0.00757 | 3.59 | 7.37 | 3.088 | 0.0723 |

The variability in the optimal parameters for the synthetic experiments is shown in the box plots in Figure 7. For a fixed agar density, the optimal biomass-production rates Ψ_*n*_ and cell-death rates Ψ_*d*_ vary across synthetic replicates. However, as expected both parameters do not depend strongly on agar density. Some studies have predicted increased cell death with increasing adhesion strength [74, 75]. Although we predict slightly higher values of Ψ_*d*_ on higher-density agar, we cannot reliably conclude that agar density affects Ψ_*d*_ in yeast from our available data. Our parameter estimates in Table 4 indicate that the nutrient-uptake rate *Q*^*^ and nutrient-consumption rate ϒ both decrease with increased agar density. The trend in *Q*^*^ is consistent with the study of He et al. [35], which found that high-density agar inhibits nutrient uptake in *E. coli* bacteria. However, there is wide variability in *Q*^*^ across the synthetic experiments. This finding is consistent with the parameter-pair heat maps in Figure 6, where *Q*^*^ was a difficult parameter to identify. The decreasing trend in ϒ with agar density is more robust than the trend in *Q*^*^, but is still subject to significant variability. The biofilm–substratum adhesion strength *λ*^*^ has the strongest variation with agar density in synthetic experiments, and the least variability across synthetic replicates. This analysis provides provides further evidence that the strongest effect of agar density is to mediate friction between the colony biofilm and substratum in *S. cerevisiae* colony biofilms.

**Figure 7:**
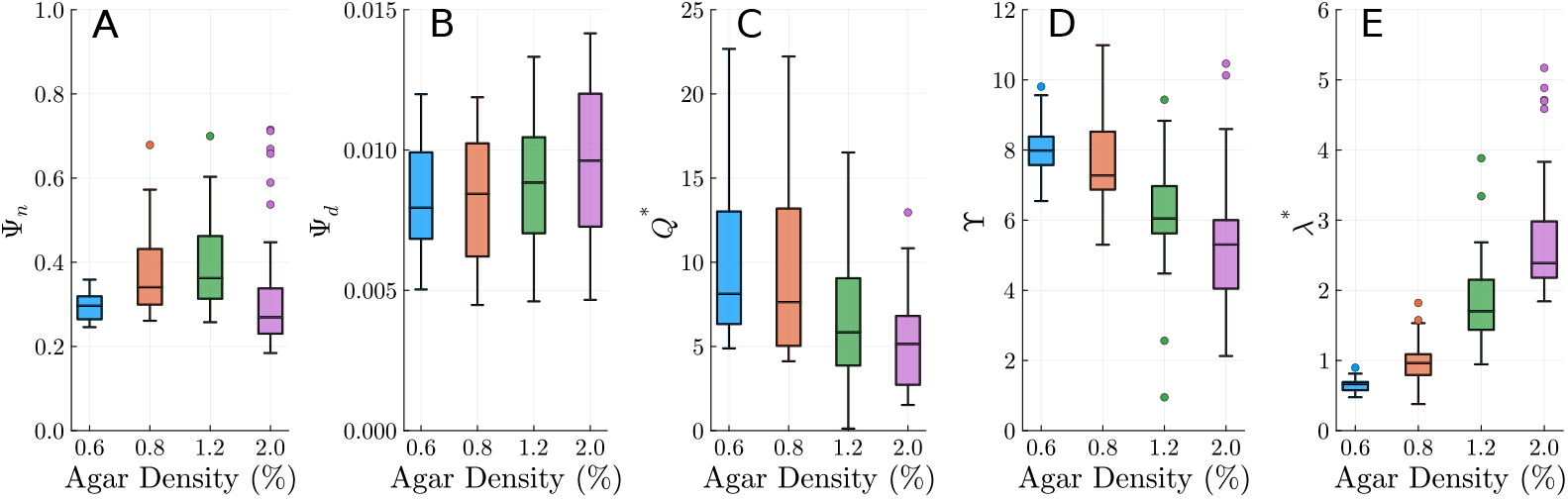
Box plots representing the optimal parameter values found using our parameter estimation procedure for 50 synthetic experiments on each agar density. Synthetic data for each of the 50 synthetic experiments was generated based on the experimental variability in cell count, half-width, and composition, using the method described in the text. Circle markers indicate outliers. (A) Ψ_*n*_. (B) Ψ_*d*_. (C) *Q*^*^. (D) ϒ. (E) *λ*^*^.

A characteristic of extensional flows is that the expansion speed depends on the initial condition [18, 19]. Since the initial colony-biofilm thickness, *H*_0_, is unknown, we also investigate the effect of *H*_0_ on the results. Varying *H*_0_ between zero and three *S. cerevisiae* cells thick reveals that the distance *ρ*(*θ, a*) is relatively insensitive to *H*_0_, compared to other parameters. This is especially true for values exceeding *H*_0_ = 0.05, which is the dimensionless thickness of a single yeast cell. Therefore, the differences between replicates are unlikely to be due to experimental variations in *H*_0_. These results are presented in Figure 8.

**Figure 8:**
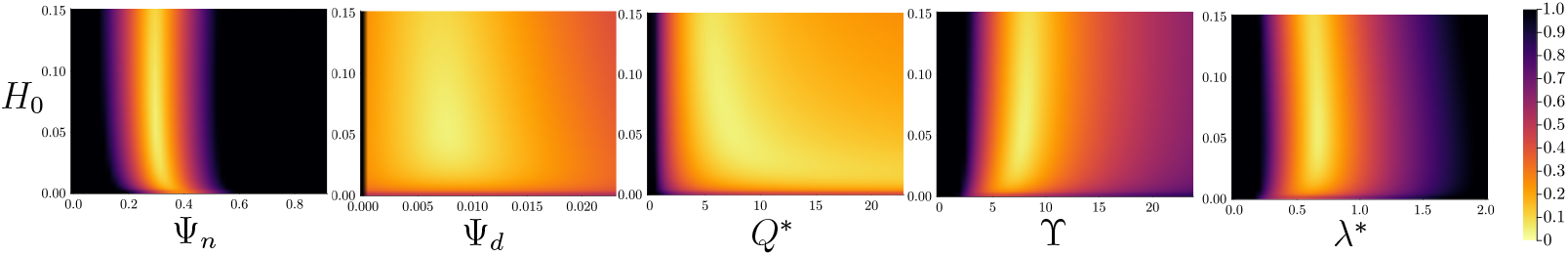
Parameter-pair heat maps for the initial colony-biofilm thickness, *H*_0_, on 0.6% agar. Plots represent the distance between the numerical solution and experimental data, *ρ*(*θ, a*), for given parameter combination. Unless otherwise stated, parameters take the optimal values. When varied, each parameter ranges from zero to three times the optimal value (2,500 total simulations).

## 5 Conclusion

In this manuscript, we have combined *S. cerevisiae* experiments and mathematical modelling to quantify how agar density and cell death impact yeast-colony-biofilm growth. Our experimental method extended previous mat formation experiments [7–9] to incorporate rectangular geometry, colony-biofilm composition measurements, and aspect-ratio measurements. We adapted the thin-film extensional-flow model of Tam et al. [19] to include cell death and resistance to slip caused by biofilm–substratum adhesion. Using new experimental data across multiple experiments, we estimated model parameters across four agar densities. The strongest impact of increasing agar density was to increase the biofilm–substratum adhesion strength, measured through the parameter *λ*^*^. This finding is consistent with previous studies in bacteria. There were also decreases in the nutrient-uptake rate and the nutrient-consumption rate on higher-density agar. These effects have also been hypothesised or observed in bacteria. However, parameter-sensitivity analysis and synthetic data yielded wider variability in the nutrient-uptake rate and consumption parameters compared to biofilm–substratum adhesion. The biomass-production rate and cell-death rate were consistent across agar densities.

Our mathematical model is based on the assumption that the colony-biofilm constituents (phases) can be modelled as Newtonian fluids [76]. Provided that this assumption is valid, the model describes the mechanics of colony-biofilm growth due to biomass proliferation, mediated by a depleting nutrient supply. Although the slip parameter *λ*^*^ quantifies the drag due to biofilm–substratum adhesion, it does not provide a full description of the substratum mechanics. The agar may dehydrate over time [35], and colony-biofilm attachment to the substratum may exert forces that deform the agar [31]. Our micrographs of the colony-biofilm cross sections indicate that these deformations are small, leading to the neglect of such deformations in the present model. To confirm that agar deformation does not alter our main conclusions, a possible extension to the model could be to incorporate the agar mechanics explicitly. This would complicate the analysis significantly, as one would need to incorporate constitutive relations for the material properties of the agar, and continuity of stress on the biofilm–substratum interface, which may also change shape over time. This extension is left for future work.

A further extension to this work is to incorporate osmotic swelling, which is known to influence bacterial-biofilm expansion [24, 77, 78]. Despite evidence that osmotic stresses indeed affect individual *S. cerevisiae* cells [79], the phenomenon of osmotic swelling in yeast colony biofilms is not widely reported. Osmotic influx from the substratum would contribute to volumetric growth of the colony biofilm, and nutrient supply. If significant, it would introduce new parameters to the model, and alter the estimates of existing parameters. Another avenue for future investigation is to better understand the non-uniform front pattern that often develops at later times, as shown in Figure 1. The amplitude of this pattern decreases on higher-density agar, and future work could focus on understanding how nutrient availability and agar density affect these interfacial instabilities.

Fungal biofilms are notorious for their ability to colonise indwelling medical devices [80, 81], and are a leading cause of hospital-acquired infections [82–85]. Our cross-sectional images reveal that a small number of yeast cells invade the agar medium in our experiments. Although the continuum model used in this paper does not permit invasion, invasive growth of yeast is important in clinical settings, because invasion of healthy tissue leads to the spread of infections [86]. Modelling the biophysics of yeast-cell invasion is a problem for future work. Other colony-biofilm experiments have featured a region of dead cells close to the leading edge. This region is thought to appear due to cells undergoing regulated cell death [30]. We neglected regulated cell death in this work, because our colony biofilms instead have regions of cell death near their inoculation site, reminiscent of a necrotic core [87, 88] expected to be formed by accidental cell death [30]. Since regulated cell death is known to occur in yeast colony biofilms, fully characterising the roles of accidental and regulated cell death [27], and their relationship to mechanics, remains an open question.

Additional future work could involve applying the multiphase model to multispecies colony biofilms, which more closely resemble the biofilms found in healthcare environments. Natural and artificial surfactants can disrupt biofilm growth in wounds and hasten their removal [89]. Whilst this effect is irrelevant to sliding motility where surface tension is negligible, it provides another promising avenue for future modelling work, as researchers continue to explore the wide range of yeast-growth modes and strategies for eliminating biofilms.

## Acknowledgements

We acknowledge funding from the Australian Research Council (Grant numbers DP230100406, and DE240100097).

## Authorship Contribution Statement

- **AKYT:** Conceptualisation, methodology, software, validation, formal analysis, investigation, data curation, writing – original draft, writing – review & editing, visualisation, funding acquisition.
- **DJN:** Conceptualisation, formal analysis, writing – review & editing.
- **JMG:** Methodology, investigation, data curation, writing – original draft.
- **JZ:** Methodology, investigation, data curation.
- **CWG:** Conceptualisation, funding acquisition.
- **VJ:** Resources, writing – review & editing, supervision, funding acquisition.
- **BJB:** Conceptualisation, writing – review & editing, project administration, funding acquisition.
- **JEFG:** Conceptualisation, writing – review & editing, supervision, funding acquisition.

## Data Availability

Open source Julia code with a README and the experimental photographs used to generate the results are available on [GitHub](https://github.com/alex-tam/YeastBiofilm_AgarSlip).

## References

[1] J. W. Costerton, P. S. Stewart, and E. P. Greenberg, “Bacterial Biofilms: A Common Cause of Persistent Infections”, Science 284 (1999), pp. 1318–1322, DOI: 10.1126/science.284.5418.1318.

[2] J. R. Blankenship and A. P. Mitchell, “How to Build a Biofilm: A Fungal Perspective”, Curr. Opin. Microbiol. 9 (2006), pp. 588–594, DOI: 10.1016/j.mib.2006.10.003.

[3] P. Stoodley, K. Sauer, D. G. Davies, and J. W. Costerton, “Biofilms as Complex Differ-entiated Communities”, Annu. Rev. Microbiol. 56 (2002), pp. 187–209, DOI: 10.1146/annurev.micro.56.012302.160705.

[4] D. Botstein, S. A. Chervitz, and M. Cherry, “Yeast as a Model Organism”, Science 277 (1997), pp. 1259–1260, DOI: 10.1126/science.277.5330.1259.

[5] R. K. Bojsen, K. S. Andersen, and B. Regenberg, “Saccharomyces cerevisiae — a Model to Uncover Molecular Mechanisms for Yeast Biofilm Biology”, FEMS Immunol. Med. Microbiol. 65 (2012), pp. 169–182, DOI: 10.1111/j.1574-695X.2012.00943.x.

[6] A. Goffeau, B. G. Barrell, H. Bussey, R. W. Davis, B. Dujon, H. Feldmann, F. Galibert, J. D. Hoheisel, C. Jacq, M. Johnston, E. J. Louis, H. W. Mewes, Y. Murakami, P. Philippsen, H. Tettelin, and S. G. Oliver, “Life with 6000 Genes”, Science 274 (1996), pp. 546–567, DOI: 10.1126/science.274.5287.546.

[7] T. B. Reynolds and G. R. Fink, “Bakers’ Yeast, a Model for Fungal Biofilm Formation”, Science 291 (2001), pp. 878–881, DOI: 10.1126/science.291.5505.878.

[8] A. Tam, J. E. F. Green, S. Balasuriya, E. L. Tek, J. M. Gardner, J. F. Sundstrom, V. Jiranek, and B. J. Binder, “Nutrient-Limited Growth with Non-Linear Cell Diffusion as a Mechanism for Floral Pattern Formation in Yeast Biofilms”, J. Theor. Biol. 448 (2018), pp. 122–141, DOI: 10.1016/j.jtbi.2018.04.004.

[9] E. L. Tek, J. F. Sundstrom, J. M. Gardner, S. G. Oliver, and V. Jiranek, “Evaluation of the Ability of Commercial Wine Yeasts to Form Biofilms (Mats) and Adhere to Plastic: Implications for the Microbiota of the Winery Environment”, FEMS Microbiol. Ecol. 94 (2018), fix188, DOI: 10.1093/femsec/fix188.

[10] T. B. Reynolds, A. Jansen, X. Peng, and G. R. Fink, “Mat Formation in Saccharomyces cerevisiae Requires Nutrient and pH Gradients”, Eukaryotic Cell 7 (2008), pp. 122–130, DOI: 10.1128/ec.00310-06.

[11] B. J. Binder, J. F. Sundstrom, J. M. Gardner, V. Jiranek, and S. G. Oliver, “Quantifying Two-Dimensional Filamentous and Invasive Growth Spatial Patterns in Yeast Colonies”, PLOS Comput. Biol. 11 (2015), e1004070, DOI: 10.1371/journal.pcbi.1004070.

[12] J. Recht, A. Martínez, S. Torello, and R. Kolter, “Genetic Analysis of Sliding Motility in Mycobacterium smegmatis”, J. Bacteriol. 182 (2000), pp. 4348–4351, DOI: 10.1128/JB.182.15.4348-4351.2000.

[13] R. M. Harshey, “Bacterial Motility on a Surface: Many Ways to a Common Goal”, Annu. Rev. Microbiol. 57 (2003), pp. 249–273, DOI: 10.1146/annurev.micro.57.030502.091014.

[14] M. R. Mattei, L. Frunzo, B. D’Acunto, Y. Pechaud, F. Pirozzi, and G. Esposito, “Continuum and Discrete Approach in Modeling Biofilm Development and Structure: A Review”, J. Math. Biol. 76 (2018), pp. 945–1003, DOI: 10.1007/s00285-017-1165-y.

[15] I. Klapper and J. Dockery, “Mathematical Description of Microbial Biofilms”, SIAM Rev. 52 (2010), pp. 221–265, DOI: 10.1137/080739720.

[16] O. Wanner and W. Gujer, “A Multispecies Biofilm Model”, Biotechnol. Bioeng. 28 (1986), pp. 314–328, DOI: 10.1002/bit.260280304.

[17] H. J. Eberl, D. F. Parker, and M. C. M. van Loosdrecht, “A New Deterministic Spatio-Temporal Continuum Model for Biofilm Development”, J. Theor. Med. 3 (2001), pp. 161–175, DOI: 10.1080/10273660108833072.

[18] J. P. Ward and J. R. King, “Thin-Film Modelling of Biofilm Growth and Quorum Sensing”, J. Eng. Math. 73 (2012), pp. 71–92, DOI: 10.1007/s10665-011-9490-4.

[19] A. Tam, J. E. F. Green, S. Balasuriya, E. L. Tek, J. M. Gardner, J. F. Sundstrom, V. Jiranek, and B. J. Binder, “A Thin-Film Extensional Flow Model for Biofilm Expansion by Sliding Motility”, Proc. Royal Soc. A 475 (2019), 20190175, DOI: 10.1098/rspa.2019.0175.

[20] A. K. Y. Tam, B. Harding, J. E. F. Green, S. Balasuriya, and B. J. Binder, “Thin-Film Lubrication Model for Biofilm Expansion under Strong Adhesion”, Phys. Rev. E 105 (2022), 014408, DOI: 10.1103/PhysRevE.105.014408.

[21] S. Trinschek, K. John, and U. Thiele, “From a Thin Film Model for Passive Suspensions towards the Description of Osmotic Biofilm Spreading”, AIMS Mater. Sci. 3 (2016), pp. 1138–1159, DOI: 10.3934/matersci.2016.3.1138.

[22] S. Trinschek, K. John, S. Lecuyer, and U. Thiele, “Continuous versus Arrested Spreading of Biofilms at Solid–Gas Interfaces: The Role of Surface Forces”, Phys. Rev. Lett. 119 (2017), 078003, DOI: 10.1103/PhysRevLett.119.078003.

[23] S. Trinschek, K. John, and U. Thiele, “Modelling of Surfactant-Driven Front Instabilities in Spreading Bacterial Colonies”, Soft Matter 14 (2018), pp. 4464–4476, DOI: 10.1039/c8sm00422f.

[24] S. Srinivasan, C. N. Kaplan, and L. Mahadevan, “A Multiphase Theory for Spreading Microbial Swarms and Films”, eLife 8 (2019), e42697, DOI: 10.7554/eLife.42697.

[25] G. T. Fortune, N. M. Oliveira, and R. E. Goldstein, “Biofilm Growth under Elastic Confinement”, Phys. Rev. Lett. 128 (2022), 178102, DOI: 10.1103/PhysRevLett.128.178102.

[26] P. Bravo, S. L. Ng, K. A. MacGillivray, B. K. Hammer, and P. J. Yunker, “Vertical Growth Dynamics of Biofilms”, Proc. Natl. Acad. Sci. U.S.A. 120 (2023), e2214211120, DOI: 10.1073/pnas.2214211120.

[27] D. Carmona-Gutierrez, M. A. Bauer, A. Zimmermann, A. Aguilera, N. Austriaco, K. Ayscough, R. Balzan, S. Bar-Nun, A. Barrientos, P. Belenky, M. Blondel, R. J. Braun, M. Breitenbach, W. C. Burhans, S. Büttner, D. Cavalieri, M. Chang, K. F. Cooper, M. Côrte-Real, V. Costa, C. Cullin, I. Dawes, J. Dengjel, M. B. Dickman, T. Eisenberg, B. Fahrenkrog, N. Fasel, K. U. Fröhlich, A. Gargouri, S. Giannattasio, P. Goffrini, C. W. Gourlay, C. M. Grant, M. T. Greenwood, N. Guaragnella, T. Heger, J. Heinisch, E. Herker, J. M. Herrmann, S. Hofer, A. Jiménez-Ruiz, H. Jungwirth, K. Kainz, D. P. Kontoyiannis, P. Ludovico, S. Manon, E. Martegani, C. Mazzoni, L. A. Megeney, C. Meisinger, J. Nielsen, T. Nyström, H. D. Osiewacz, T. F. Outeiro, H. Park, T. Pendl, D. Petranovic, S. Picot, P. Polčic, T. Powers, M. Ramsdale, M. Rinnerthaler, P. Rockenfeller, C. Ruckenstuhl, R. Schaffrath, M. Segovia, F. F. Severin, A. Sharon, S. J. Sigrist, C. Sommer-Ruck, M. J. Sousa, J. M. Thevelein, K. Thevissen, V. Titorenko, M. B. Toledano, M. Tuite, F. N. Vögtle, B. Westermann, J. Winderickx, S. Wissing, S. Wölfl, Z. J. Zhang, R. Y. Zhao, B. Zhou, L. Galluzzi, G. Kroemer, and F. Madeo, “Guidelines and Recommendations on Yeast Cell Death Nomenclature”, Microb. Cell 5 (2018), pp. 4–31, DOI: 10.15698/mic2018.01.607, PMID: 29354647.

[28] Z. Palková and L. Váchová, “Cell Differentiation, Aging, and Death in Spatially Organized Yeast Communities: Mechanisms and Consequences”, Cell Death Differ. (2025), pp. 1–13, DOI: 10.1038/s41418-025-01485-9.

[29] M. Čáp, L. Štěpánek, K. Harant, L. Váchová, and Z. Palková, “Cell Differentiation within a Yeast Colony: Metabolic and Regulatory Parallels with a Tumor-Affected Organism”, Mol. Cell 46 (2012), pp. 436–448, DOI: 10.1016/j.molcel.2012.04.001, PMID: 22560924.

[30] D. J. Netherwood, A. K. Y. Tam, C. W. Gourlay, T. Knežević, J. M. Gardner, V. Jiranek, B. J. Binder, and J. E. F. Green, “Accidental and Regulated Cell Death in Yeast Colony Biofilms”, Bull. Math. Biol. 87 (2025), 110, DOI: 10.1101/2025.02.02.636168.

[31] A. Pietz, K. John, and U. Thiele, “The Role of Substrate Mechanics in Osmotic Biofilm Spreading”, Soft Matter 21 (2025), pp. 2935–2945, DOI: 10.1039/D4SM01463D.

[32] A. R. Pokhrel, R. Copeland, M. Hejri, T. E. R. Belpaire, G. Steinbach, S. L. Ng, B. K. Hammer, and P. J. Yunker, Cell-Substrate Friction Controls Biofilm Development, 2025, DOI: 10.1101/2025.07.11.664457, URL: https://www.biorxiv.org/content/10.1101/2025.07.11.664457v1 (visited on 07/21/2025), pre-published.

[33] A. K. Verma, A. Mookherjee, C. Tropini, D. Fusco, and L. Ruiz Pestana, “The Interplay of Adhesion, Friction, and Nutrient Availability in Modulating Biofilm Wrinkling Behavior”, npj Soft Matter 1 (2025), p. 8, DOI: 10.1038/s44431-025-00007-4.

[34] C. Fei, S. Mao, J. Yan, R. Alert, H. A. Stone, B. L. Bassler, N. S. Wingreen, and A. Košmrlj, “Nonuniform Growth and Surface Friction Determine Bacterial Biofilm Morphology on Soft Substrates”, Proc. Natl. Acad. Sci. U.S.A. 117 (2020), pp. 7622–7632, DOI: 10.1073/pnas.1919607117.

[35] C. He, L. Han, D. C. Harris, S. Bayakhmetov, X. Wang, and Y. Kuang, “Reaction– Diffusion Modeling of E. coli Colony Growth Based on Nutrient Distribution and Agar Dehydration”, Bull. Math. Biol. 85 (2023), p. 61, DOI: 10.1007/s11538-023-01163-2.

[36] W. J. Middelhoven, B. Broekhuizen, and J. van Eijk, “Detection, with the Dye Phloxine B, of Yeast Mutants Unable to Utilize Nitrogenous Substances as the Sole Nitrogen Source”, J. Bacteriol. 128 (1976), pp. 851–852, DOI: 10.1128/jb.128.3.851-852.1976.

[37] J. F. Cannon, J. B. Gibbs, and K. Tatchell, “Supperssors of the Ras2 Mutation of Saccharomyces cerevisiae”, Genetics 113 (1986), pp. 247–264, DOI: 10.1093/genetics/113.2.247.

[38] I. Klapper, C. J. Rupp, R. Cargo, B. Purvedorj, and P. Stoodley, “Viscoelastic Fluid Description of Bacterial Biofilm Material Properties”, Biotechnol. Bioeng. 80 (2002), pp. 289–296, DOI: 10.1002/bit.10376.

[39] D. A. Drew, “Mathematical Modeling of Two-Phase Flow”, Annu. Rev. Fluid Mech. 15 (1983), pp. 261–291, DOI: 10.1146/annurev.fl.15.010183.001401.

[40] N. Kandemir, W. Vollmer, N. S. Jakubovics, and J. Chen, “Mechanical Interactions between Bacteria and Hydrogels”, Sci. Rep. 8 (2018), 10893, DOI: 10.1038/s41598-018-29269-x.

[41] S. Gomez, L. Bureau, K. John, E.-N. Chêne, D. Débarre, and S. Lecuyer, “Substrate Stiffness Impacts Early Biofilm Formation by Modulating Pseudomonas aeruginosa Twitching Motility”, eLife 12 (2023), e81112, DOI: 10.7554/eLife.81112.

[42] J. Maršíková, D. Wilkinson, O. Hlaváček, G. D. Gilfillan, A. Mizeranschi, T. R. Hughes, M. Begany, S. Rešetárová, L. Váchová, and Z. Palková, “Metabolic Differentiation of Surface and Invasive Cells of Yeast Colony Biofilms Revealed by Gene Expression Profiling”, BMC Genom. 18 (2017), 814, DOI: 10.1186/s12864-017-4214-4.

[43] S. J. Franks and J. R. King, “Interactions between a Uniformly Proliferating Tumour and Its Surroundings: Uniform Material Properties”, Math. Med. Biol. 20 (2003), pp. 47–89, DOI: 10.1093/imammb/20.1.47.

[44] R. D. O’Dea, S. L. Waters, and H. M. Byrne, “A Multiphase Model for Tissue Construct Growth in a Perfusion Bioreactor”, Math. Med. Biol. 27 (2010), pp. 95–127, DOI: 10.1093/imammb/dqp003.

[45] D. A. Drew and L. A. Segel, “Averaged Equations for Two-Phase Flows”, Stud. Appl. Math. 50 (1971), pp. 205–231, DOI: 10.1002/sapm1971503205.

[46] G. Lemon, J. R. King, H. M. Byrne, O. E. Jensen, and K. M. Shakesheff, “Mathematical Modelling of Engineered Tissue Growth Using a Multiphase Porous Flow Mixture Theory”, J. Math. Biol. 52 (2006), pp. 571–594, DOI: 10.1007/s00285-005-0363-1.

[47] J. E. F. Green, J. P. Whiteley, J. M. Oliver, H. M. Byrne, and S. L. Waters, “Pattern Formation in Multiphase Models of Chemotactic Cell Aggregation”, Math. Med. Biol. 35 (2018), pp. 319–346, DOI: 10.1093/imammb/dqx005.

[48] R. D. O’Dea, S. L. Waters, and H. M. Byrne, “A Two-Fluid Model for Tissue Growth within a Dynamic Flow Environment”, Eur. J. Appl. Math. 19 (2008), pp. 607–634, DOI: 10.1017/S0956792508007687.

[49] S. J. Franks, H. M. Byrne, J. R. King, J. C. E. Underwood, and C. E. Lewis, “Modelling the Early Growth of Ductal Carcinoma in Situ of the Breast”, J. Math. Biol. 47 (2003), pp. 424–452, DOI: 10.1007/s00285-003-0214-x.

[50] S. R. Lubkin and T. L. Jackson, “Multiphase Mechanics of Capsule Formation in Tumors”, J. Biomech. Eng. 124 (2002), pp. 237–243, DOI: 10.1115/1.1427925.

[51] G. Forgacs, R. A. Foty, Y. Shafrir, and M. S. Steinberg, “Viscoelastic Properties of Living Embryonic Tissues: A Quantitative Study”, Biophys. J. 74 (1998), pp. 2227–2234, DOI: 10.1016/S0006-3495(98)77932-9.

[52] J. R. King and J. M. Oliver, “Thin-Film Modelling of Poroviscous Free Surface Flows”, Eur. J. Appl. Math. 16 (2005), pp. 519–553, DOI: 10.1017/s095679250500584x.

[53] P. D. Howell, B. Scheid, and H. A. Stone, “Newtonian Pizza: Spinning a Viscous Sheet”, J. Fluid Mech. 659 (2010), pp. 1–23, DOI: 10.1017/S0022112010001564.

[54] L. G. Longsworth, “Diffusion in Liquids and the Stokes–Einstein Relation”, Electrochemistry in Biology and Medicine, ed. by T. Shedlovsky, First edition, New York: John Wiley & Sons, Ltd, 1955, pp. 225–247.

[55] A. L. Slade, A. E. Cremers, and H. C. Thomas, “The Obstruction Effect in the Self-Diffusion Coefficients of Sodium and Cesium in Agar Gels”, J. Phys. Chem. 70 (1966), pp. 2840–2844, DOI: 10.1021/j100881a020.

[56] P. S. Stewart, “A Review of Experimental Measurements of Effective Diffusive Permeabilities and Effective Diffusion Coefficients in Biofilms”, Biotechnol. Bioeng. 59 (1998), pp. 261–272, DOI: 10.1002/(SICI)1097-0290(19980805)59:3<261::AID-BIT1>3.0.CO;2-9.

[57] M. E. Asp, M.-T. Ho Thanh, D. A. Germann, R. J. Carroll, A. Franceski, R. D. Welch, A. Gopinath, and A. E. Patteson, “Spreading Rates of Bacterial Colonies Depend on Substrate Stiffness and Permeability”, PNAS Nexus 1 (2022), pgac025, DOI: 10.1093/pnasnexus/pgac025.

[58] F. D. Martinez-Garcia, T. Fischer, A. Hayn, C. T. Mierke, J. K. Burgess, and M. C. Harmsen, “A Beginner’s Guide to the Characterization of Hydrogel Microarchitecture for Cellular Applications”, Gels 8 (2022), 535, DOI: 10.3390/gels8090535.

[59] I. Jayawardena, P. Turunen, B. C. Garms, A. Rowan, S. Corrie, and L. Grøndahl, “Evaluation of Techniques Used for Visualisation of Hydrogel Morphology and Determination of Pore Size Distributions”, Mater. Adv. 4 (2023), pp. 669–682, DOI: 10.1039/D2MA00932C.

[60] A. Tam, “Mathematical Modelling of Pattern Formation in Yeast Biofilms”, PhD thesis, The University of Adelaide, 2019, HDL: 2440/122613.

[61] C. J. Gimeno, P. O. Ljungdahl, C. A. Styles, and G. R. Fink, “Unipolar Cell Divisions in the Yeast S. cerevisiae Lead to Filamentous Growth: Regulation by Starvation and RAS”, Cell 68 (1992), pp. 1077–1090, DOI: 10.1016/0092-8674(92)90079-R.

[62] M. Zakhartsev and M. Reuss, “Cell Size and Morphological Properties of Yeast Saccharomyces cerevisiae in Relation to Growth Temperature”, FEMS Yeast Res. 18 (2018), foy052, DOI: 10.1093/femsyr/foy052.

[63] K. Li, J. E. F. Green, H. Tronnolone, A. K. Y. Tam, A. J. Black, J. M. Gardner, J. F. Sundstrom, V. Jiranek, and B. J. Binder, “An Off-Lattice Discrete Model to Characterise Filamentous Yeast Colony Morphology”, PLOS Comput. Biol. 20 (2024), e1012605, DOI: 10.1371/journal.pcbi.1012605.

[64] R. M. Storn and K. V. Price, “Differential Evolution – a Simple and Efficient Heuristic for Global Optimization over Continuous Spaces”, J. Glob. Optim. 11 (1997), pp. 341–359, DOI: 10.1023/A:1008202821328.

[65] K. V. Price, R. M. Storn, and J. A. Lampinen, Differential Evolution: A Practical Approach to Global Optimization, Natural Computing Series, Berlin/Heidelberg: Springer-Verlag, 2005, ISBN: 978-3-540-20950-8, DOI: 10.1007/3-540-31306-0.

[66] C. G. Broyden, “The Convergence of a Class of Double-Rank Minimization Algorithms 1. General Considerations”, IMA J. Appl. Math. 6 (1970), pp. 76–90, DOI: 10.1093/imamat/6.1.76.

[67] R. Fletcher, “A New Approach to Variable Metric Algorithms”, Comput. J. 13 (1970), pp. 317–322, DOI: 10.1093/comjnl/13.3.317.

[68] D. Goldfarb, “A Family of Variable-Metric Methods Derived by Variational Means”, Math. Comp. 24 (1970), pp. 23–26, DOI: 10.1090/S0025-5718-1970-0258249-6.

[69] D. F. Shanno, “Conditioning of Quasi-Newton Methods for Function Minimization”, Math. Comp. 24 (1970), pp. 647–656, DOI: 10.1090/S0025-5718-1970-0274029-X.

[70] P. K. Mogensen and A. N. Risbeth, “Optim: A Mathematical Optimization Package for Julia”, J. Open Source Softw. 3 (2018), 615, DOI: 10.21105/joss.00615.

[71] W. W. Hager and H. Zhang, “Algorithm 851: CG_DESCENT, a Conjugate Gradient Method with Guaranteed Descent”, ACM Trans. Math. Softw. 32 (2006), pp. 113–137, DOI: 10.1145/1132973.1132979.

[72] M. L. Stecchini, M. Del Torre, S. Donda, E. Maltini, and S. Pacor, “Influence of Agar Content on the Growth Parameters of Bacillus cereus”, Int. J. Food Microbiol. 64 (2001), pp. 81–88, DOI: 10.1016/S0168-1605(00)00436-0.

[73] R. Ziege, A. Tsirigoni, B. Large, D. O. Serra, K. G. Blank, R. Hengge, P. Fratzl, and C. M. Bidan, “Adaptation of Escherichia coli Biofilm Growth, Morphology, and Mechanical Properties to Substrate Water Content”, ACS Biomater. Sci. Eng. 7 (2021), pp. 5315–5325, DOI: 10.1021/acsbiomaterials.1c00927.

[74] L. Z. Y. Huang, Z. L. Shaw, R. Penman, S. Cheeseman, V. K. Truong, M. J. Higgins, R. A. Caruso, and A. Elbourne, “Cell Adhesion, Elasticity, and Rupture Forces Guide Microbial Cell Death on Nanostructured Antimicrobial Titanium Surfaces”, ACS Appl. Bio Mater. 7 (2024), pp. 344–361, DOI: 10.1021/acsabm.3c00943.

[75] S. Desai, K. Sanghrajka, and D. Gajjar, “High Adhesion and Increased Cell Death Contribute to Strong Biofilm Formation in Klebsiella pneumoniae”, Pathogens 8 (2019), 277, DOI: 10.3390/pathogens8040277.

[76] M. S. Steinberg, “Reconstruction of Tissues by Dissociated Cells”, Science 141 (1963), pp. 401–408, DOI: 10.1126/science.141.3579.401.

[77] J. Yan, C. D. Nadell, H. A. Stone, N. S. Wingreen, and B. L. Bassler, “Extracellular-Matrix-Mediated Osmotic Pressure Drives Vibrio cholerae Biofilm Expansion and Cheater Exclusion”, Nat. Commun. 8 (2017), 327, DOI: 10.1038/s41467-017-00401-1.

[78] A. Seminara, T. E. Angelini, J. N. Wilking, H. Vlamakis, S. Ebrahim, R. Kolter, D. A. Weitz, and M. P. Brenner, “Osmotic Spreading of Bacillus subtilis Biofilms Driven by an Extracellular Matrix”, Proc. Natl. Acad. Sci. U.S.A. 109 (2012), pp. 1116–1121, DOI: 10.1073/pnas.1109261108.

[79] G. J. Morris, L. Winters, G. E. Coulson, and K. J. Clarke, “Effect of Osmotic Stress on the Ultrastructure and Viability of the Yeast Saccharomyces cerevisiae”, Microbiology 132 (1986), pp. 2023–2034, DOI: 10.1099/00221287-132-7-2023.

[80] E. M. Kojic and R. O. Darouiche, “Candida Infections of Medical Devices”, Clin. Microbiol. Rev. 17 (2004), pp. 255–267, DOI: 10.1128/cmr.17.2.255-267.2004.

[81] L. M. Martinez and B. C. Fries, “Fungal Biofilms: Relevance in the Setting of Human Disease”, Curr. Fungal Infect. Rep. 4 (2010), pp. 266–275, DOI: 10.1007/s12281-010-0035-5.

[82] C. J. Nobile and A. D. Johnson, “Candida albicans Biofilms and Human Disease”, Annu. Rev. Microbiol. 69 (2015), pp. 71–92, DOI: 10.1146/annurev-micro-091014-104330.

[83] G. Ramage, R. Rajendran, L. Sherry, and C. Williams, “Fungal Biofilm Resistance”, Int. J. Microbiol. 2012 (2012), 528521, DOI: 10.1155/2012/528521.

[84] P. G. Pappas, M. S. Lionakis, M. C. Arendrup, L. Ostrosky-Zeichner, and B. J. Kullberg, “Invasive Candidiasis”, Nat. Rev. Dis. Primers 4 (2018), 18026, DOI: 10.1038/nrdp.2018.26.

[85] M. S. Lionakis, “New Insights into Innate Immune Control of Systemic Candidiasis”, Med. Mycol. 52 (2014), pp. 555–564, DOI: 10.1093/mmy/myu029.

[86] P. J. Cullen and G. F. Sprague, “Glucose Depletion Causes Haploid Invasive Growth in Yeast”, Proc. Natl. Acad. Sci. U.S.A. 97 (2000), pp. 13619–13624, DOI: 10.1073/pnas.240345197.

[87] H. P. Greenspan, “Models for the Growth of a Solid Tumor by Diffusion”, Stud. Appl. Math. 51 (1972), pp. 317–340, DOI: 10.1002/sapm1972514317.

[88] H. M. Byrne and M. A. J. Chaplain, “Modelling the Role of Cell-Cell Adhesion in the Growth and Development of Carcinomas”, Math. Comput. Model. 24 (1996), pp. 1–17, DOI: 10.1016/s0895-7177(96)00174-4.

[89] S. L. Percival, D. Mayer, R. S. Kirsner, G. Schultz, D. Weir, S. Roy, A. Alavi, and M. Romanelli, “Surfactants: Role in Biofilm Management and Cellular Behaviour”, Int. Wound J. 16 (2019), pp. 753–760, DOI: 10.1111/iwj.13093.

