## Supporting Material for "Quantifying the effects of cell death and agar density on yeast colony biofilms using an extensional-flow mathematical model"

---

**D Code and Data Availability** **26**

**References** **26**

### A Experimental Results

#### A.1 Experimental Photographs and Image Processing

Figure A.1 illustrates a time series of photographs for one replicate of colony-biofilm growth on each agar density. We used photographs to obtain experimental measurements of the colony-biofilm half-width. To achieve this, we imported the photograph using Julia's `FileIO` and `Images` packages. We then converted the photograph to a binary image using Otsu's method. An example binary image is shown in Figure A.2.

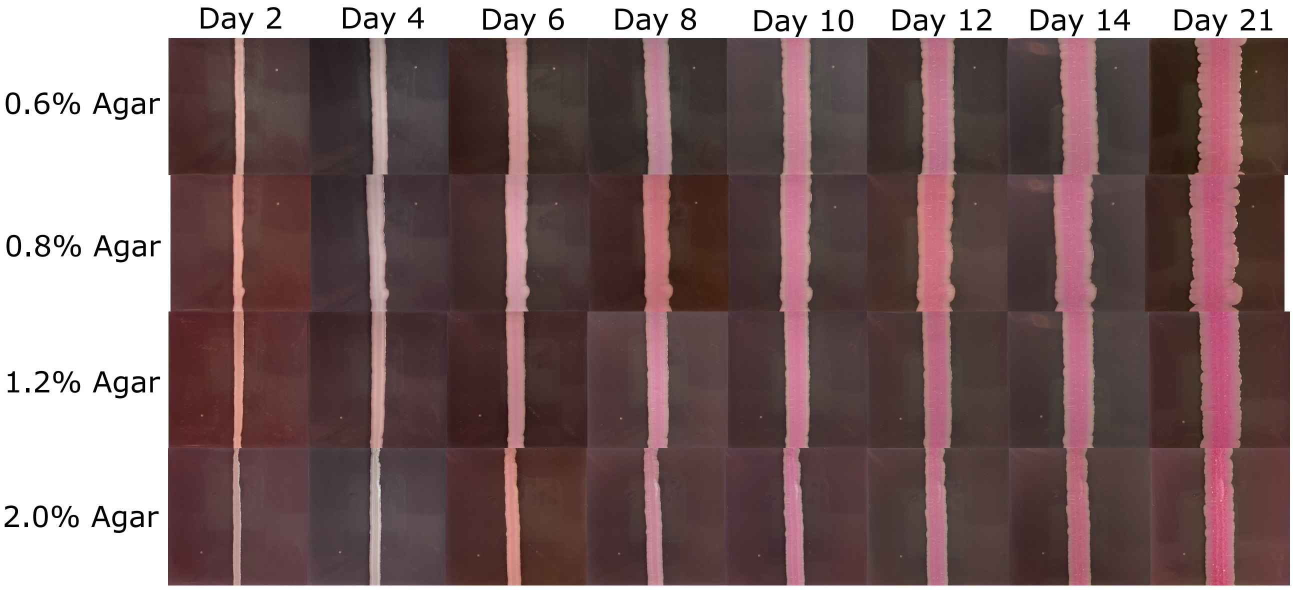

Figure A.1: Time series of cropped photographs for one replicate of colony-biofilm growth on each agar density.

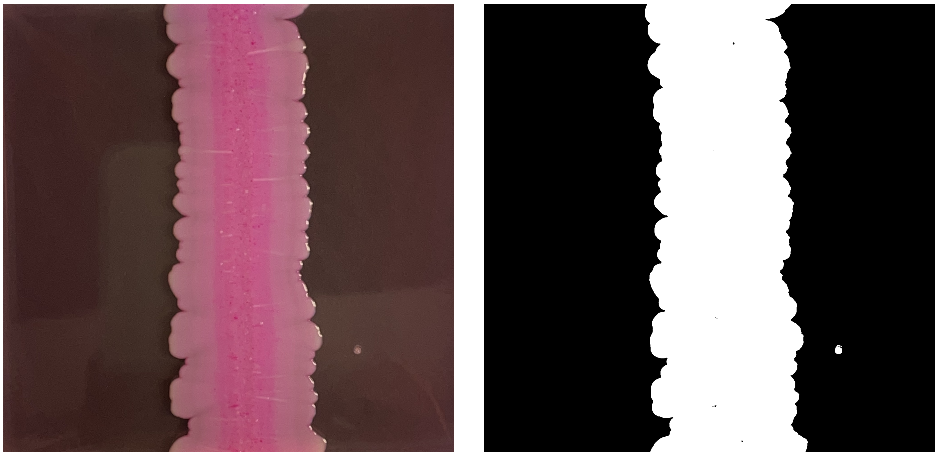

(a) Experiment.

(b) Binary image.

Figure A.2: (a) Experimental photograph for replicate 2 on 0.8% agar, after 21 days of growth. (b) Corresponding binary image obtained using Otsu's method.

After binarisation, we computed the total number of occupied (white) pixels in the largest connected component in the binary image, which will be the colony biofilm. Then, we divided by the height (in pixels) of the image to obtain the mean colony-biofilm width, in pixels. Using a ruler placed next to the Petri dish (not shown in Figure A.2), we then determined the physical width of a pixel in each photograph. This allowed us to convert the mean colony-biofilm width in pixels to a half-width in physical units. Repeating this process on all experimental photographs yielded $S(t)$ for each experimental replicate. We used the standard deviation in $S(t)$ across all replicates of the same agar density when generating synthetic data, as described in the main manuscript.

#### A.2 Cell Counts

The cell-count data were collected by sampling from a $4\,\text{cm}^2$ region of the colony biofilms on Day 21. The collection regions were rectangles with dimensions $4\,\text{cm} \times 1\,\text{cm}$, the blue boxes in Figure A.3. After sampling, cells were counted using a hemocytometer and a Nikon Eclipse 50i microscope. The total cell counts and the standard deviations are shown in Table A.1.

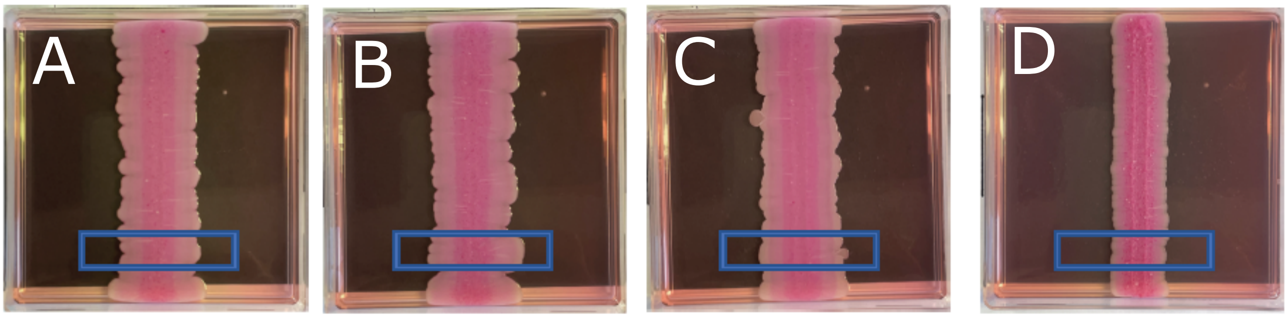

Figure A.3: Colony-biofilm photographs on Day 21 indicating the regions (blue box) from which the cell-count samples were collected. (A) 0.6% agar. (B) 0.8% agar. (C) 1.2% agar. (D) 2.0% agar.

Table A.1: Experimental cell-count data from Day 21 of yeast growth on agar media of different density. The final column is the ratio of the standard deviation to the mean cell count for that agar density, which was used in the synthetic data generation.

| **Agar Density (%)** | $n$ | **Mean Cell Count** [cells] | **Standard Deviation** [cells] | **Relative Std. Dev.** |
|---|---|---|---|---|
| 0.6 | 4 | $1.06 \times 10^9$ | $9.25 \times 10^7$ | 0.0873 |
| 0.8 | 4 | $1.40 \times 10^9$ | $1.82 \times 10^8$ | 0.130 |
| 1.2 | 4 | $1.60 \times 10^9$ | $1.64 \times 10^8$ | 0.103 |
| 2.0 | 3 | $1.12 \times 10^9$ | $1.38 \times 10^8$ | 0.123 |

Although the cell counts were taken from a $4\,\text{cm}^2$ region for each colony biofilm, the sampled area of the colony biofilm varies with agar density, because the colony-biofilm half-width $S(t)$ varies with agar density. The cell counts presented in the main manuscript represent the average number of cells across the region of width $S(t)$ in the $x$-direction, per millimetre in the $y$-direction. This cell number then provided a way to estimate the colony-biofilm height, which we converted to an aspect ratio because $S(t)$ was known. For the synthetic data, we generated a random cell count by sampling from a normal distribution with mean and standard deviation as per Table A.1. We then converted the cell count to the aspect ratio using the procedure described in the main manuscript, assuming constant $S(t)$.

#### A.3 Cell Viability

We measured the proportion of living and dead cells in the colony biofilm on Day 14 of growth. Flow cytometry was used to detect Phloxine B stained cells, which we took to indicate dead cells [1, 2]. Collecting these data is invasive, so these experiments were terminated on Day 14 and not included in the data set for the colony-biofilm half-width. To avoid terminating more experiments than necessary, we used two experimental replicates to characterise cell viability, one on each of 0.6% and 2.0% agar. A summary of the cell-viability experimental data is shown in Figure A.4.

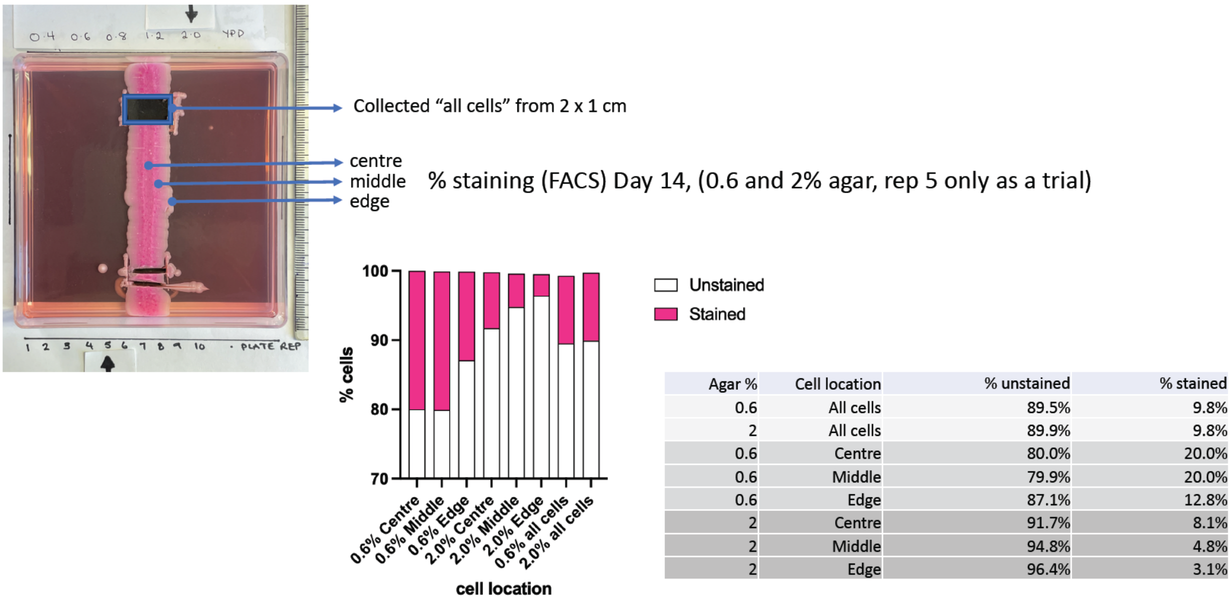

| Agar % | Cell location | % unstained | % stained |
|---|---|---|---|
| 0.6 | All cells | 89.5% | 9.8% |
| 2 | All cells | 89.9% | 9.8% |
| 0.6 | Centre | 80.0% | 20.0% |
| 0.6 | Middle | 79.9% | 20.0% |
| 0.6 | Edge | 87.1% | 12.8% |
| 2 | Centre | 91.7% | 8.1% |
| 2 | Middle | 94.8% | 4.8% |
| 2 | Edge | 96.4% | 3.1% |

Figure A.4: Experimental procedure and results for the cell viability measurements.

As the annotated experimental photograph indicates, the total cell sampling from the blue box provided a larger and more reliable source of cell viability data. The proportions of viable cells were very similar for colonies grown on 0.6% and 2.0% agar. The viability for three spatial locations across the colony biofilm are also provided in Figure A.4. Since these were collected from small samples, there is variability in the results for 0.6% and 2.0% agar. However, on both agar densities the cell viability was lower in the centre than at the edge. We took the mean of the 0.6% and 2.0% data to yield the cell-viability data used in the parameter estimation. When generating synthetic data, we scaled the mean cell-viability data at each location by the same factor drawn from the uniform distribution $U(0.932, 1.05)$. This ensured that the minimum possible viability in the synthetic data at $x = 0$ is 80%, and that the maximum possible viability at $x = S(t)$ is 96.4%, as per the experimental results in Figure A.4.

### B Mathematical Model Derivation

Our mathematical model closely follows Tam [3] and Tam et al. [4], but we present full details here for completeness. We model colony-biofilm expansion over an agar substratum, from which the colony biofilm obtains nutrients [3–6]. We formulate the model in 1D Cartesian geometry, as illustrated in Figure B.1.

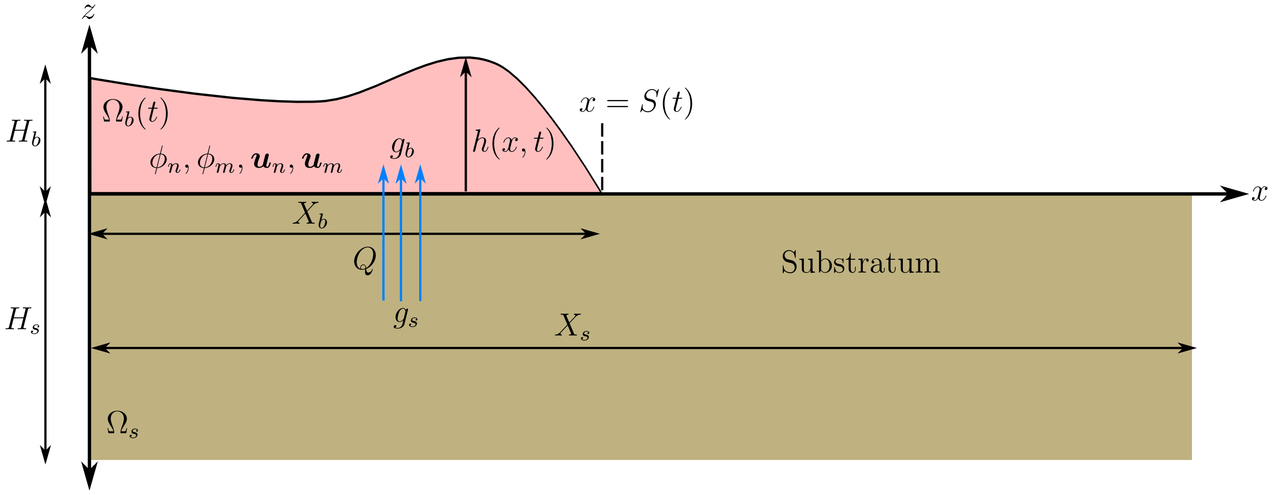

Figure B.1: Schematic of the mathematical model and colony-biofilm geometry. See the text for the definition of each symbol.

The colony biofilm exists in the region $\Omega_b(t)$ bounded by $0 < x < S(t)$ and $0 < z < h(x,t)$. The domain $\Omega_s$ represents the agar substratum, and is the fixed region bounded by $-H_s < z < 0$, and $0 < x < X_s$ where $H_s$ and $X_s$ are the Petri-dish depth and width, respectively. We assume that the colony biofilm consists of two phases, both of which are Newtonian viscous fluids. The phases are an active biomass phase containing proliferating cells and EPS, and an inactive biomass phase consisting of dead cells and extracellular material. We formulate the model using mixture theory, where the volume fractions $\phi_n$ and $\phi_m$ satisfy the no-voids condition, $\phi_n + \phi_m = 1$. Since the two phases cannot physically occupy the same space, when introducing the volume fractions we assume that appropriate ensemble averaging has occurred, according to the process outlined by Drew [7]. We assume that the agar substratum cannot deform. The substratum contains nutrients, which we assume occupy no volume. The colony biofilm can take up nutrients through the biofilm–substratum interface. After uptake, nutrients become available for consumption, driving cell proliferation and colony-biofilm expansion. Under this two-phase framework, the model variables are

- $h(x,t)$: Colony-biofilm height.
- $\phi_n(x,z,t)$: Volume fraction of active biomass (living cells, EPS).
- $\phi_m(x,z,t)$: Volume fraction of inactive biomass (dead cells, water).
- $g_s(x,z,t)$: Nutrient concentration in the substratum, defined in $\Omega_s$.
- $g_b(x,z,t)$: Nutrient concentration in the colony biofilm, defined in $\Omega_b(t)$.

- $\boldsymbol{u}_\alpha(x, y, t) = (u_\alpha(x, y, t), w_\alpha(x, y, t))$: Fluid velocity of phase $\alpha = n, m$.
- $p_\alpha(x, z, t)$: Fluid pressure of phase $\alpha = n, m$.

We also use the symbols $X_b$ and $H_b$ to denote the characteristic width and height of the colony biofilm, respectively.

#### B.1 Mass Conservation

We obtain governing equations using the principles of mass conservation and momentum conservation. The volume fractions for both phases obey the continuity equation

$$\frac{\partial \rho_\alpha}{\partial t} + \nabla \cdot (\rho_\alpha \boldsymbol{u}_\alpha) = J_\alpha, \tag{B.1}$$

where $\rho_\alpha$ is the density of phase $\alpha$, and $J_\alpha$ is a source term describing net production of phase $\alpha$. We assume that both fluid phases are incompressible, such that their densities $\rho_\alpha$ are constant. For the source terms $J_\alpha$, we assume first-order kinetics, whereby cell proliferation is proportional to the local active volume fraction and nutrient concentration, and cell death is proportional to the local active volume fraction. Upon death, cells immediately become part of the inactive phase, with no corresponding change in volume. Under these assumptions, the no-voids and mass-conservation equations for the two phases read

$$\phi_n + \phi_m = 1, \tag{B.2a}$$

$$\frac{\partial \phi_n}{\partial t} + \frac{\partial}{\partial x}(\phi_n u_n) + \frac{\partial}{\partial z}(\phi_n w_n) = \psi_n \phi_n g_b - \psi_d \phi_n, \tag{B.2b}$$

$$\frac{\partial \phi_m}{\partial t} + \frac{\partial}{\partial x}(\phi_m u_m) + \frac{\partial}{\partial z}(\phi_m w_m) = \psi_d \phi_n, \tag{B.2c}$$

where $\psi_n$ is the cell-proliferation rate and $\psi_d$ is the cell-death rate, both of which we assume constant. Adding (B.2b) and (B.2c) and applying the no-voids condition allows us to eliminate $\phi_m$ to write the system (B.2) in the form

$$\frac{\partial u}{\partial x} + \frac{\partial w}{\partial z} = \psi_n \phi_n g_b, \tag{B.3a}$$

$$\frac{\partial \phi_n}{\partial t} + \frac{\partial}{\partial x}(\phi_n u_n) + \frac{\partial}{\partial z}(\phi_n w_n) = \psi_n \phi_n g_b - \psi_d \phi_n, \tag{B.3b}$$

where $\boldsymbol{u} = (u, w) = (\phi_n u_n + \phi_m u_m, \phi_n w_n + \phi_m w_m)$, is the mixture-averaged fluid velocity.

Nutrient disperse by diffusion in the substratum, and by diffusion and advection in the colony biofilm, where they also become available for consumption. Thus, the nutrient concentrations satisfy the equations

$$\frac{\partial g_s}{\partial t} = D_s\left(\frac{\partial^2 g_s}{\partial x^2} + \frac{\partial^2 g_s}{\partial z^2}\right), \tag{B.4a}$$

$$\frac{\partial g_b}{\partial t} + \frac{\partial}{\partial x}(g_b u) + \frac{\partial}{\partial z}(g_b w) = D_b\left(\frac{\partial^2 g_b}{\partial x^2} + \frac{\partial^2 g_b}{\partial z^2}\right) - \eta\phi_n g_b, \tag{B.4b}$$

where $D_s$ and $D_b$ are nutrient diffusion coefficients, and $\eta$ is the maximum nutrient-consumption rate.

#### B.2 Momentum Conservation

We obtain the remaining governing equations using conservation of momentum. Since the fluid flows in colony biofilms are very viscous, with Re $\approx 0.001$ [8], we neglect inertia such that Cauchy's momentum equation applied to one phase becomes

$$\nabla\cdot(\phi_\alpha\boldsymbol{\sigma}_\alpha) + \boldsymbol{F}_\alpha = \mathbf{0}, \tag{B.5}$$

where $\boldsymbol{\sigma}_\alpha$ is the stress tensor, and $\boldsymbol{F}_\alpha$ represents external sources of momentum. We assume that the entire colony biofilm is a Newtonian viscous material, such that the stress tensor for a given phase is

$$\boldsymbol{\sigma}_\alpha = -\left(p_\alpha + \frac{2\mu_\alpha}{3}\nabla\cdot\boldsymbol{u}_\alpha\right)\boldsymbol{I} + \mu_\alpha\left[\nabla\boldsymbol{u}_\alpha + (\nabla\boldsymbol{u}_\alpha)^T\right], \tag{B.6}$$

where $\boldsymbol{I}$ is the identity tensor, and $\mu_\alpha$ is the dynamic viscosity. Owing to cell proliferation, the velocity field is not necessarily divergence free, so we invoke Stokes' hypothesis [4, 9–11], such that the coefficient of the bulk-viscosity (divergence of velocity) term in (B.6) is $-2\mu_\alpha/3$. Due to Newton's third law, the momentum sources are equal and opposite, $\boldsymbol{F}_n = -\boldsymbol{F}_m$. We assume that these momentum sources consist of interphase drag and interfacial forces. Since both phases consist of biological material dispersed within a continuous fluid, we propose interfacial forces of the same form as Lemon et al. [12], as originally derived by Drew and Segel [13]. This yields $\boldsymbol{F}_n = -k(\boldsymbol{u}_n - \boldsymbol{u}_m) + p_n\nabla\phi_n$ and $\boldsymbol{F}_m = -k(\boldsymbol{u}_m - \boldsymbol{u}_n) + p_m\nabla\phi_m$, where $k(\phi_n, \phi_m)$ is the interphase drag coefficient. Componentwise, the momentum balances then contribute two equations per phase (four equations total) to the model,

$$-\frac{\partial p_\alpha}{\partial x} + \mu_\alpha\left(\frac{\partial^2 u_\alpha}{\partial x^2} + \frac{\partial^2 u_\alpha}{\partial z^2}\right) + \frac{\mu_\alpha}{3}\frac{\partial}{\partial x}\left(\frac{\partial u_\alpha}{\partial x} + \frac{\partial w_\alpha}{\partial z}\right) = k(u_\alpha - u_\beta), \tag{B.7a}$$

$$-\frac{\partial p_\alpha}{\partial z} + \mu_\alpha\left(\frac{\partial^2 w_\alpha}{\partial x^2} + \frac{\partial^2 w_\alpha}{\partial z^2}\right) + \frac{\mu_\alpha}{3}\frac{\partial}{\partial z}\left(\frac{\partial u_\alpha}{\partial x} + \frac{\partial w_\alpha}{\partial z}\right) = k(w_\alpha - w_\beta), \tag{B.7b}$$

where the subscript $\beta$ denotes the opposite phase to $\alpha$, $\mu_\alpha$ is the dynamic viscosity for phase $\alpha$.

#### B.3 Initial and Boundary Conditions

When deriving the model we leave the initial conditions general, and write

$$g_s(x, z, 0) = \mathcal{G}(x, z), \tag{B.8a}$$
$$g_b(x, z, 0) = 0, \tag{B.8b}$$
$$\phi_n(x, z, 0) = \Phi(x, z), \tag{B.8c}$$
$$h(x, 0) = \mathcal{H}(x), \tag{B.8d}$$

Precise forms for (B.8) will be determined later, informed by experiments. Several boundary conditions are required to close the model, and we initially consider conditions on horizontal interfaces. For the nutrients, we include conditions on the substratum base, biofilm–substratum interface, and the colony-biofilm free surface. Nutrients cannot pass through the base of the substratum, yielding a no-flux condition. In general, the no-flux condition is $\boldsymbol{q} \cdot \hat{\boldsymbol{n}} = 0$, where $\boldsymbol{q}$ is the flux and $\hat{\boldsymbol{n}}$ is the unit outward normal vector to the relevant surface. At the substratum base, the no-flux condition is

$$\frac{\partial g_s}{\partial z} = 0 \quad \text{on} \quad z = -H_s. \tag{B.9}$$

Colony biofilms take up nutrients from the substratum. We assume that nutrient uptake occurs at a rate proportional to the difference in nutrient concentrations across the biofilm–substratum interface. This yields the two conditions

$$D_s \frac{\partial g_s}{\partial z} = -Q\left(g_s - g_b\right) \quad \text{on} \quad z = 0, \tag{B.10a}$$
$$D_b \frac{\partial g_b}{\partial z} = -Q\left(g_s - g_b\right) \quad \text{on} \quad z = 0, \tag{B.10b}$$

where $Q$ is a mass-transfer coefficient describing the rate of nutrient uptake. Finally, nutrients in the colony biofilm cannot pass through the free surface. Since the unit outward normal vector to the free surface is $\hat{\boldsymbol{n}} = (-h_x, 1)$, we obtain the no-flux condition

$$g_b\left(u\frac{\partial h}{\partial x} - w\right) = D_b\left(\frac{\partial g_b}{\partial x}\frac{\partial h}{\partial x} - \frac{\partial g_b}{\partial z}\right) \quad \text{on} \quad z = h. \tag{B.11}$$

We obtain the remaining boundary conditions for the colony biofilm from standard conditions from fluid mechanics. On the biofilm–substratum interface, the no-penetration condition yields

$$w_\alpha = 0 \quad \text{on} \quad z = 0. \tag{B.12}$$

The tangential-stress condition on the biofilm–substratum interface reads $\hat{\boldsymbol{t}} \cdot (\phi_\alpha \boldsymbol{\sigma}_\alpha \cdot \hat{\boldsymbol{n}}) = \lambda_\alpha(\phi_\alpha \boldsymbol{u}_\alpha \cdot \hat{\boldsymbol{t}})$, where $\hat{\boldsymbol{t}}$ is any unit tangent vector, and the new constant $\lambda_\alpha$ is a slip coefficient representing the adhesion strength between the fluid phase and the agar. This yields the slip boundary condition

$$\mu_\alpha \left( \frac{\partial u_\alpha}{\partial z} + \frac{\partial w_\alpha}{\partial x} \right) = \mu_\alpha \frac{\partial u_\alpha}{\partial z} = \lambda_\alpha u_\alpha \quad \text{on} \quad z = 0, \tag{B.13}$$

Perfect slip corresponds to $\lambda_\alpha = 0$, whereas no slip requires $\lambda_\alpha \to \infty$. On the colony-biofilm free surface, the kinematic condition

$$\frac{\partial h}{\partial t} + u_\alpha \frac{\partial h}{\partial x} = w_\alpha \quad \text{on} \quad z = h, \tag{B.14}$$

states that fluid particles initially on the free surface must remain there. We also have one condition per phase stating that the tangential stress is zero on the free surface. That is, $\hat{\boldsymbol{t}} \cdot (\phi_\alpha \boldsymbol{\sigma}_\alpha \cdot \hat{\boldsymbol{n}}) = 0$. In expanded form this condition reads

$$-2\frac{\partial h}{\partial x}\left( \frac{\partial u_\alpha}{\partial x} - \frac{\partial w_\alpha}{\partial z} \right) + \frac{\partial u_\alpha}{\partial z} + \frac{\partial w_\alpha}{\partial x} - \left( \frac{\partial h}{\partial x} \right)^2 \left( \frac{\partial u_\alpha}{\partial z} + \frac{\partial w_\alpha}{\partial x} \right) = 0 \quad \text{on} \quad z = h. \tag{B.15}$$

Finally, we impose zero normal stress on the free surface, $\hat{\boldsymbol{n}} \cdot (\phi_\alpha \boldsymbol{\sigma}_\alpha \cdot \hat{\boldsymbol{n}}) = 0$. This boundary condition assumes zero surface tension. Surface tension might represent the typical fluid surface tension, or the strength of cell–cell adhesion at the biofilm–air interface [14]. For colony biofilms expanding by sliding motility [15, 16], cell–cell adhesion forces are weak. Previously, we showed that surface tension has minimal impact on the colony-biofilm expansion speed [4]. Surface tension primarily impacted colony-biofilm shape by mitigating ridge formation, an effect that is unimportant to the experimental results in this manuscript. Therefore, we neglect surface tension in the model. The normal-stress condition on the free surface is then

$$\begin{aligned} & p_\alpha + \frac{2\mu_\alpha}{3}\left( \frac{\partial u_\alpha}{\partial x} + \frac{\partial w_\alpha}{\partial z} \right) \\ &= \mu_\alpha \left[ 2\left( \frac{\partial h}{\partial x} \right)^2 \frac{\partial u_\alpha}{\partial x} + 2\frac{\partial w_\alpha}{\partial z} - \frac{\partial h}{\partial x}\left( \frac{\partial u_\alpha}{\partial z} + \frac{\partial w_\alpha}{\partial x} \right) \right] \left[ 1 + \left( \frac{\partial h}{\partial x} \right)^2 \right]^{-3/2} \quad \text{on} \quad z = h. \end{aligned} \tag{B.16}$$

Equations (B.2), (B.4) and (B.7), combined with initial conditions (B.8) and boundary conditions (B.9)–(B.16), then form the closed model.

#### B.4 Model Simplification

We apply several simplifying assumptions prior to nondimensionalisation. O'Dea, Waters, and Byrne [17] built on several studies [18, 19] to show that the interphase drag coefficient is large for flow in a bioreactor. Since the length scales of colony-biofilm growth are similar to a bioreactor [17], the same large-drag assumption applies. Taking $k \to \infty$ requires $\boldsymbol{u}_n = \boldsymbol{u}_m$ for the momentum source terms to remain bounded. Therefore, one simplification is that both phases move at the common mixture velocity $\boldsymbol{u} = \boldsymbol{u}_n = \boldsymbol{u}_m$. Furthermore, although the precise composition of the colony biofilm is complex, the active and passive biomass phases consist of water, extracellular polymeric substances (EPS), and cellular material. Therefore, we can assume that both phases have common density and viscosity, $\rho_n = \rho_m$ and $\mu_n = \mu_m$. The strength of biofilm–substratum adhesion may depend on the agar density, but can also be assumed the same across the two phases, $\lambda = \lambda_n = \lambda_m$. Owing to the similarity of other physical properties, the pressure, $p = p_n = p_m$, is also assumed the same for both phases. Under these assumptions, the model reduces to (where all PDEs apply in $\Omega_b(t)$, except the equation for $g_s$ which applies in $\Omega_s$)

$$\frac{\partial u}{\partial x} + \frac{\partial w}{\partial z} = \psi_n \phi_n g_b, \tag{B.17a}$$

$$\frac{\partial \phi_n}{\partial t} + \frac{\partial}{\partial x}(\phi_n u) + \frac{\partial}{\partial z}(\phi_n w) = \psi_n \phi_n g_b - \psi_d \phi_n, \tag{B.17b}$$

$$\frac{\partial g_s}{\partial t} = D_s\left(\frac{\partial^2 g_s}{\partial x^2} + \frac{\partial^2 g_s}{\partial z^2}\right), \tag{B.17c}$$

$$\frac{\partial g_b}{\partial t} + \frac{\partial}{\partial x}(g_b u) + \frac{\partial}{\partial z}(g_b w) = D_b\left(\frac{\partial^2 g_b}{\partial x^2} + \frac{\partial^2 g_b}{\partial z^2}\right) - \eta \phi_n g_b, \tag{B.17d}$$

$$-\frac{\partial p}{\partial x} + \mu\left(\frac{\partial^2 u}{\partial x^2} + \frac{\partial^2 u}{\partial z^2}\right) + \frac{\mu}{3}\frac{\partial}{\partial x}\left(\frac{\partial u}{\partial x} + \frac{\partial w}{\partial z}\right) = 0, \tag{B.17e}$$

$$-\frac{\partial p}{\partial z} + \mu\left(\frac{\partial^2 w}{\partial x^2} + \frac{\partial^2 w}{\partial z^2}\right) + \frac{\mu}{3}\frac{\partial}{\partial z}\left(\frac{\partial u}{\partial x} + \frac{\partial w}{\partial z}\right) = 0, \tag{B.17f}$$

and is subject to the initial conditions (B.8) and boundary conditions

$$\frac{\partial g_s}{\partial z} = 0 \quad \text{on} \quad z = -H_s, \tag{B.18a}$$

$$D_s \frac{\partial g_s}{\partial z} = -Q\left(g_s - g_b\right) \quad \text{on} \quad z = 0, \tag{B.18b}$$

$$D_b \frac{\partial g_b}{\partial z} = -Q\left(g_s - g_b\right) \quad \text{on} \quad z = 0, \tag{B.18c}$$

$$w = 0 \quad \text{on} \quad z = 0, \tag{B.18d}$$

$$\mu \frac{\partial u}{\partial z} = \lambda u \quad \text{on} \quad z = 0, \tag{B.18e}$$

$$g_b\left(u\frac{\partial h}{\partial x} - w\right) = D_b\left(\frac{\partial g_b}{\partial x}\frac{\partial h}{\partial x} - \frac{\partial g_b}{\partial z}\right) \quad \text{on} \quad z = h, \tag{B.18f}$$

$$\frac{\partial h}{\partial t} + u\frac{\partial h}{\partial x} = w \quad \text{on} \quad z = h, \tag{B.18g}$$

$$-2\frac{\partial h}{\partial x}\left(\frac{\partial u}{\partial x} - \frac{\partial w}{\partial z}\right) + \frac{\partial u}{\partial z} + \frac{\partial w}{\partial x} - \left(\frac{\partial h}{\partial x}\right)^2\left(\frac{\partial u}{\partial z} + \frac{\partial w}{\partial x}\right) = 0 \quad \text{on} \quad z = h, \tag{B.18h}$$

$$\begin{aligned} &p + \frac{2\mu}{3}\left(\frac{\partial u}{\partial x} + \frac{\partial w}{\partial z}\right) \\ &= \mu\left[2\left(\frac{\partial h}{\partial x}\right)^2\frac{\partial u}{\partial x} + 2\frac{\partial w}{\partial z} - \frac{\partial h}{\partial x}\left(\frac{\partial u}{\partial z} + \frac{\partial w}{\partial x}\right)\right]\left[1 + \left(\frac{\partial h}{\partial x}\right)^2\right]^{-3/2} \end{aligned} \quad \text{on} \quad z = h. \tag{B.18i}$$

#### B.5 Nondimensionalisation

We nondimensionalise by introducing dimensionless variables denoted with hats, and the scalings

$$\begin{gathered} t = \frac{X_b^2}{D_b}\hat{t}, \quad (x, z) = (X_b\hat{x}, \varepsilon X_b\hat{z}), \quad (u, w) = \left(\frac{D_b}{X_b}\hat{u}, \frac{\varepsilon D_b}{X_b}\hat{w}\right), \\ g_s = G\hat{g}_s, \quad g_b = G\hat{g}_b, \quad \text{and} \quad p = \frac{\mu D_b}{X_b^2}\hat{p}, \end{gathered} \tag{B.19}$$

where $X_b = S(0)$ is the initial colony-biofilm size, and $\varepsilon = H_b/X_b = H_s/X_s \ll 1$ is a small thin-film parameter. The dimensionless governing equations are then (where all PDEs apply in $\Omega_b(t)$, except the equation for $g_s$ which applies in $\Omega_s$)

$$\frac{\partial u}{\partial x} + \frac{\partial w}{\partial z} = \Psi_n \phi_n g_b, \tag{B.20a}$$

$$\frac{\partial \phi_n}{\partial t} + \frac{\partial}{\partial x}(\phi_n u) + \frac{\partial}{\partial z}(\phi_n w) = \Psi_n \phi_n g_b - \Psi_d \phi_n, \tag{B.20b}$$

$$\frac{\partial g_s}{\partial t} = D\left(\frac{\partial^2 g_s}{\partial x^2} + \frac{1}{\varepsilon^2}\frac{\partial^2 g_s}{\partial z^2}\right), \tag{B.20c}$$

$$\frac{\partial g_b}{\partial t} + \frac{\partial}{\partial x}(g_b u) + \frac{\partial}{\partial z}(g_b w) = \frac{\partial^2 g_b}{\partial x^2} + \frac{1}{\varepsilon^2}\frac{\partial^2 g_b}{\partial z^2} - \Upsilon \phi_n g_b, \tag{B.20d}$$

$$-\frac{\partial p}{\partial x} + \frac{\partial^2 u}{\partial x^2} + \frac{1}{\varepsilon^2}\frac{\partial^2 u}{\partial z^2} + \frac{1}{3}\frac{\partial}{\partial x}\left(\frac{\partial u}{\partial x} + \frac{\partial w}{\partial z}\right) = 0, \tag{B.20e}$$

$$-\frac{\partial p}{\partial z} + \varepsilon^2\frac{\partial^2 w}{\partial x^2} + \frac{\partial^2 w}{\partial z^2} + \frac{1}{3}\frac{\partial}{\partial z}\left(\frac{\partial u}{\partial x} + \frac{\partial w}{\partial z}\right) = 0, \tag{B.20f}$$

subject to the boundary conditions,

$$\frac{\partial g_s}{\partial z} = 0 \quad \text{on} \quad z = -1/\delta, \tag{B.21a}$$

$$\frac{\partial g_s}{\partial z} = -\frac{Q^\dagger}{D}\left(g_s - g_b\right) \quad \text{on} \quad z = 0, \tag{B.21b}$$

$$\frac{\partial g_b}{\partial z} = -Q^\dagger\left(g_s - g_b\right) \quad \text{on} \quad z = 0, \tag{B.21c}$$

$$w = 0 \quad \text{on} \quad z = 0, \tag{B.21d}$$

$$\frac{\partial u}{\partial z} = \lambda^\dagger u \quad \text{on} \quad z = 0, \tag{B.21e}$$

$$g_b\left(u\frac{\partial h}{\partial x} - w\right) = \frac{\partial g_b}{\partial x}\frac{\partial h}{\partial x} - \frac{1}{\varepsilon^2}\frac{\partial g_b}{\partial z} \quad \text{on} \quad z = h, \tag{B.21f}$$

$$\frac{\partial h}{\partial t} + u\frac{\partial h}{\partial x} = w \quad \text{on} \quad z = h, \tag{B.21g}$$

$$-2\frac{\partial h}{\partial x}\left(\frac{\partial u}{\partial x} - \frac{\partial w}{\partial z}\right) + \frac{1}{\varepsilon^2}\frac{\partial u}{\partial z} + \frac{\partial w}{\partial x} - \left(\frac{\partial h}{\partial x}\right)^2\left(\frac{\partial u}{\partial z} + \varepsilon^2\frac{\partial w}{\partial x}\right) = 0 \quad \text{on} \quad z = h, \tag{B.21h}$$

$$\begin{aligned} p + \frac{2}{3}\left(\frac{\partial u}{\partial x} + \frac{\partial w}{\partial z}\right) = \Bigg[2\varepsilon^2\left(\frac{\partial h}{\partial x}\right)^2\frac{\partial u}{\partial x} + 2\frac{\partial w}{\partial z} \\ -\frac{\partial h}{\partial x}\left(\frac{\partial u}{\partial z} + \varepsilon^2\frac{\partial w}{\partial x}\right)\Bigg]\left[1 + \varepsilon^2\left(\frac{\partial h}{\partial x}\right)^2\right]^{-3/2} \end{aligned} \quad \text{on} \quad z = h. \tag{B.21i}$$

In writing (B.20) and (B.21), we have introduced the dimensionless parameters

$$\begin{aligned} &\delta = \frac{X_b}{X_s}, \quad \Psi_n = \frac{GX_b^2\psi_n}{D_b}, \quad \Psi_d = \frac{X_b^2\psi_d}{D_b}, \quad D = \frac{D_s}{D_b}, \\ &\Upsilon = \frac{X_b^2\eta}{D_b}, \quad Q^\dagger = \frac{H_b Q}{D_b}, \quad \text{and} \quad \lambda^\dagger = \frac{H_b\lambda}{\mu}. \end{aligned} \tag{B.22}$$

Previously, we assumed that the substratum and colony biofilm were of similar size in Tam et al. [4]. Here, we account for the difference in length scales in the colony biofilm and substratum by defining the parameter $\delta$. After introducing the dimensionless parameters (B.22), we subsequently consider the distinguished limit in which $Q^\dagger$ and $\lambda^\dagger$ are of $\mathcal{O}(\varepsilon^2)$ size as $\varepsilon \to 0$. This small $\lambda^\dagger$ limit represents weak adhesion suitable for biofilm expansion by sliding motility, and small $Q^\dagger$ represents slow nutrient transfer across the biofilm–substratum interface. Accordingly, we rescale $Q^\dagger$ and $\lambda^\dagger$ by setting $Q^\dagger \sim \varepsilon^2 Q^*$ and $\lambda^\dagger \sim \varepsilon^2\lambda^*$, where $Q^*$ and $\lambda^*$ are $\mathcal{O}(1)$ quantities as $\varepsilon \to 0$. In terms of the rescaled quantities $Q^*$ and $\lambda^*$, the boundary conditions (B.21b), (B.21c) and (B.21e) then become

$$\frac{\partial g_s}{\partial z} = -\varepsilon^2 \frac{Q^*}{D} (g_s - g_b) \quad \text{on} \quad z = 0, \tag{B.23a}$$

$$\frac{\partial g_b}{\partial z} = -\varepsilon^2 Q^* (g_s - g_b) \quad \text{on} \quad z = 0, \tag{B.23b}$$

$$\frac{\partial u}{\partial z} = \varepsilon^2 \lambda^* u \quad \text{on} \quad z = 0. \tag{B.23c}$$

#### B.6 Thin-Film Approximation

We now apply the thin-film (long-wave) approximation to obtain a model to leading order in $\varepsilon$. We expand variables in power series of $\varepsilon^2$,

$$h(x, y, t) \sim h_0(x, y, t) + \varepsilon^2 h_1(x, y, t) + \mathcal{O}(\varepsilon^4), \tag{B.24}$$

and so on, where the expansions for the other variables take the same form as (B.24). To leading order in $\varepsilon$, the governing equations are then

$$\frac{\partial u_0}{\partial x} + \frac{\partial w_0}{\partial z} = \Psi_n \phi_{n0} g_{b0}, \tag{B.25a}$$

$$\frac{\partial \phi_{n0}}{\partial t} + \frac{\partial}{\partial x} (\phi_{n0} u_0) + \frac{\partial}{\partial z} (\phi_{n0} w_0) = \Psi_n \phi_{n0} g_{b0} - \Psi_d \phi_{n0}, \tag{B.25b}$$

$$\frac{\partial^2 g_{s0}}{\partial z^2} = 0, \tag{B.25c}$$

$$\frac{\partial^2 g_{b0}}{\partial z^2} = 0, \tag{B.25d}$$

$$\frac{\partial^2 u_0}{\partial z^2} = 0, \tag{B.25e}$$

$$-\frac{\partial p_0}{\partial z} + \frac{\partial^2 w_0}{\partial z^2} + \frac{1}{3} \frac{\partial}{\partial z} \left( \frac{\partial u_0}{\partial x} + \frac{\partial w_0}{\partial z} \right) = 0, \tag{B.25f}$$

and the leading-order boundary conditions become

$$\frac{\partial g_{s0}}{\partial z} = 0 \quad \text{on} \quad z = -1/\delta, 0, \tag{B.26a}$$

$$\frac{\partial g_{b0}}{\partial z} = 0 \quad \text{on} \quad z = 0, h_0, \tag{B.26b}$$

$$\frac{\partial u_0}{\partial z} = 0 \quad \text{on} \quad z = 0, h_0, \tag{B.26c}$$

$$w_0 = 0 \quad \text{on} \quad z = 0, \tag{B.26d}$$

$$w_0 = \frac{\partial h_0}{\partial t} + u_0 \frac{\partial h_0}{\partial x} \quad \text{on} \quad z = h_0, \tag{B.26e}$$

$$-p_0 - \frac{2}{3} \frac{\partial u_0}{\partial x} + \frac{4}{3} \frac{\partial w_0}{\partial z} = 0 \quad \text{on} \quad z = h_0. \tag{B.26f}$$

As expected for an extensional flow, the systems (B.25) and (B.26) indicate that the leading-order nutrient concentrations $g_{s0}$ and $g_{b0}$, and the leading-order horizontal velocity $u_0$ do not depend on $z$. To proceed, we also define the depth-averaged cell volume fraction

$$\bar{\phi}_0(x,t) = \frac{1}{h_0}\int_0^{h_0} \phi_{n0}(x,z,t)\,\mathrm{d}z. \tag{B.27}$$

On defining (B.27), we exploit the $z$-independence and integrate the mass-balance equations (B.25a) and (B.25b) with respect to $z$ from 0 to $h_0$, yielding

$$\frac{\partial h_0}{\partial t} + \frac{\partial}{\partial x}(u_0 h_0) = \Psi_n \bar{\phi}_0 g_{b0} h_0, \tag{B.28a}$$

$$\frac{\partial}{\partial t}\left(\bar{\phi}_0 h_0\right) + \frac{\partial}{\partial x}\left(\bar{\phi}_0 u_0 h_0\right) = \Psi_n \bar{\phi}_0 g_{b0} h_0 - \Psi_d \bar{\phi}_0 h_0, \tag{B.28b}$$

where we have applied Leibniz's integral rule in obtaining (B.28b).

Since the leading-order equations for $g_{s0}$, $g_{b0}$, and $u_0$ yielded $z$-independence, to obtain governing equations for these variables we consider the higher-order correction terms. Using the dimensionless model and thin-film expansions (B.20), (B.21) and (B.24), the relevant equations at $\mathcal{O}(\varepsilon^2)$ are

$$\frac{\partial^2 g_{s1}}{\partial z^2} = \frac{1}{D}\frac{\partial g_{s0}}{\partial t} - \frac{\partial^2 g_{s0}}{\partial x^2}, \tag{B.29a}$$

$$\frac{\partial^2 g_{b1}}{\partial z^2} = \frac{\partial g_{b0}}{\partial t} + \frac{\partial}{\partial x}(g_{b0}u_0) + \frac{\partial}{\partial z}(g_{b0}w_0) - \frac{\partial^2 g_{b0}}{\partial x^2} + \Upsilon \phi_{n0} g_{b0}, \tag{B.29b}$$

$$\frac{\partial^2 u_1}{\partial z^2} = \frac{\partial p_0}{\partial x} - \frac{\partial^2 u_0}{\partial x^2} - \frac{1}{3}\frac{\partial}{\partial x}\left(\frac{\partial u_0}{\partial x} + \frac{\partial w_0}{\partial z}\right), \tag{B.29c}$$

with boundary conditions

$$\frac{\partial g_{s1}}{\partial z} = 0 \qquad \text{on} \quad z = -1/\delta, \tag{B.30a}$$

$$\frac{\partial g_{s1}}{\partial z} = -\frac{Q^*}{D}(g_{s0} - g_{b0}) \qquad \text{on} \quad z = 0, \tag{B.30b}$$

$$\frac{\partial g_{b1}}{\partial z} = -Q^*(g_{s0} - g_{b0}) \qquad \text{on} \quad z = 0, \tag{B.30c}$$

$$\frac{\partial g_{b1}}{\partial z} = \frac{\partial g_{b0}}{\partial x}\frac{\partial h_0}{\partial x} - g_{b0}\left(u_0\frac{\partial h_0}{\partial x} - w_0\right) \qquad \text{on} \quad z = h_0, \tag{B.30d}$$

$$\frac{\partial u_1}{\partial z} = \lambda^* u_0 \qquad \text{on} \quad z = 0, \tag{B.30e}$$

$$\frac{\partial u_1}{\partial z} = 2\frac{\partial h_0}{\partial x}\left(\frac{\partial u_0}{\partial x} - \frac{\partial w_0}{\partial z}\right) - \frac{\partial w_0}{\partial x} \qquad \text{on} \quad z = h_0. \tag{B.30f}$$

Integrating the correction terms for the nutrient concentration (B.29a) and (B.29b) with respect to $z$ and applying the boundary conditions (B.30a)–(B.30d) yields equations for the leading-order nutrient concentrations,

$$\frac{\partial g_{s0}}{\partial t} = D\frac{\partial^2 g_{s0}}{\partial x^2} - \delta Q^* \left(g_{s0} - g_{b0}\right), \tag{B.31a}$$

$$h_0\frac{\partial g_{b0}}{\partial t} + \frac{\partial}{\partial x}\left(u_0 h_0 g_{b0}\right) = \frac{\partial}{\partial x}\left(h_0\frac{\partial g_{b0}}{\partial x}\right) + Q^*\left(g_{s0} - g_{b0}\right) - \Upsilon\bar{\phi}_0 g_{b0} h_0. \tag{B.31b}$$

A similar depth-integration procedure yields an equation for the leading-order horizontal velocity, $u_0$. First, we rewrite the correction for the $x$-component of momentum (B.29c) using the mass balance (B.25a) to obtain

$$\frac{\partial^2 u_1}{\partial z^2} = \frac{\partial p_0}{\partial x} - 2\frac{\partial^2 u_0}{\partial x^2} - \frac{\partial^2 w_0}{\partial x\,\partial z} + \frac{2\Psi_n}{3}\frac{\partial}{\partial x}\left(\phi_{n0} g_{b0}\right). \tag{B.32}$$

Integrating the correction term for the horizontal velocity (B.32) once with respect to $z$ and applying the boundary conditions (B.30e) and (B.30f) yields, on application of Leibniz's integral rule for pressure and volume fraction terms,

$$\begin{aligned} 2\frac{\partial h_0}{\partial x}\left(\frac{\partial u_0}{\partial x} - \left.\frac{\partial w_0}{\partial z}\right|_{z=h_0}\right) - \left.\frac{\partial w_0}{\partial x}\right|_{z=h_0} - \lambda^* u_0 = \frac{\partial}{\partial x}\left(\int_0^{h_0} p_0 \,\mathrm{d}z\right) - p_0(h_0)\frac{\partial h_0}{\partial x} \\ -2h_0\frac{\partial^2 u_0}{\partial x^2} - \left[\frac{\partial w_0}{\partial x}\right]_0^{h_0} + \frac{2\Psi_n}{3}\frac{\partial}{\partial x}\left(\int_0^{h_0} \phi_{n0} g_{b0}\,\mathrm{d}z\right) - \frac{2\Psi_n}{3}\phi_{n0}(h_0) g_{b0}\frac{\partial h_0}{\partial x}. \end{aligned} \tag{B.33}$$

Derivatives of $w_0$ with respect to $x$ cancel. To solve for the leading-order pressure $p_0$, we integrate (B.25f) to obtain

$$p_0 = \frac{4}{3}\frac{\partial w_0}{\partial z} + \frac{1}{3}\frac{\partial u_0}{\partial x} + C(x). \tag{B.34}$$

Applying the boundary condition (B.26f) for the leading-order pressure on $z = h_0$ determines the constant of integration

$$C(x) = -\frac{\partial u_0}{\partial x} \implies p_0 = -\frac{4}{3}\frac{\partial w_0}{\partial z} - \frac{2}{3}\frac{\partial u_0}{\partial x} = -2\frac{\partial u_0}{\partial x} + \frac{4\Psi_n}{3}\phi_{n0} g_{b0} = 2\frac{\partial w_0}{\partial z} - \frac{2\Psi_n}{3}\phi_{n0} g_{b0}, \tag{B.35}$$

where we have used the mass balance (B.25a) to obtain two alternative forms for $p_0$. Substituting the expression for pressure (B.35) into the integrated $\mathcal{O}(\varepsilon^2)$ momentum equation (B.33) then yields

$$\begin{aligned} 2\frac{\partial h_0}{\partial x}\left(\frac{\partial u_0}{\partial x} - \left.\frac{\partial w_0}{\partial z}\right|_{z=h_0}\right) - \lambda^* u_0 = \frac{\partial}{\partial x}\left(\int_0^{h_0} \frac{4\Psi_n}{3}\phi_{n0} g_{b0} - 2\frac{\partial u_0}{\partial x}\,\mathrm{d}z\right) \\ - \frac{\partial h_0}{\partial x}\left[2\left.\frac{\partial w_0}{\partial z}\right|_{z=h_0} - \frac{2\Psi_n}{3}\phi_{n0}(h_0) g_{b0}\right] \\ -2h_0\frac{\partial^2 u_0}{\partial x^2} + \frac{2\Psi_n}{3}\frac{\partial}{\partial x}\left(\int_0^{h_0} \phi_{n0} g_{b0}\,\mathrm{d}z\right) - \frac{2\Psi_n}{3}\phi_{n0}(h_0) g_{b0}\frac{\partial h_0}{\partial x}. \end{aligned} \tag{B.36}$$

After evaluating the integrals and simplification, we arrive at the leading-order governing equation

$$4\frac{\partial}{\partial x}\left(h_0\frac{\partial u_0}{\partial x}\right) - \lambda^* u_0 = 2\Psi_n\frac{\partial}{\partial x}\left(\bar{\phi}_0 g_{b0} h_0\right). \tag{B.37}$$

Equations (B.28), (B.31) and (B.37) then constitute a system of five spatially one-dimensional PDEs for the leading-order quantities $h_0$, $\bar{\phi}_0$, $g_{s0}$, $g_{b0}$, and $u_0$. Recall that Equations (B.37), (B.28a), (B.28b) and (B.31b) are defined within the colony biofilm $x < S(t)$. The nutrient concentration in the substratum (B.31a) is defined for the Petri dish, $0 < x < 1/\delta$. Since nutrient depletion from the substratum can only occur in regions occupied by the colony biofilm, we modify (B.31a) to write

$$\frac{\partial g_{s0}}{\partial t} = D\frac{\partial^2 g_{s0}}{\partial x^2} - \delta Q^* \left(g_{s0} - g_{b0}\right) H(S(t) - x) \quad \text{on} \quad 0 < x < 1/\delta, \tag{B.38}$$

where $H(\cdot)$ is the Heaviside step function.

It remains to specify the initial and boundary conditions for PDEs (B.28), (B.37), (B.38) and (B.31b). The initial conditions are

$$g_{s0}(x,0) = 1, \tag{B.39a}$$
$$g_{b0}(x,0) = 0, \tag{B.39b}$$
$$\bar{\phi}_0(x,0) = 0, \tag{B.39c}$$
$$h_0(x,0) = H_0\left(1 - x^2\right), \tag{B.39d}$$
$$S(0) = 1. \tag{B.39e}$$

These conditions reflect that there are initially no nutrients or dead cells in the colony biofilm. The substratum nutrient concentration $g_{s0}$ and colony-biofilm half-width $S$ are initially unity because nutrients are scaled by the initial nutrient concentration, and $x$ is scaled by the initial colony-biofilm width. Since the initial cell density in experiments is several orders of magnitude less than the cell density of a mature colony biofilm [20], we expect $H_0$ to be small, even with the thin-film approximation applied.

The boundary conditions for the spatially one-dimensional model consist of no-flux conditions for the nutrients, which become

$$\frac{\partial g_{s0}}{\partial x} = 0 \quad \text{on} \quad x = 0, 1/\delta, \quad \text{and} \quad \frac{\partial g_{b0}}{\partial x} = 0 \quad \text{on} \quad x = 0. \tag{B.40}$$

We also apply the following symmetry conditions at $x = 0$:

$$\frac{\partial h_0}{\partial x} = 0, \quad \frac{\partial \bar{\phi}_0}{\partial x} = 0, \quad u_0 = 0 \quad \text{on} \quad x = 0. \tag{B.41}$$

We obtain the second boundary condition for $g_{b0}$ using the same scaling argument as Tam et al. [4], yielding

$$\frac{\partial g_{b0}}{\partial x} = 0 \quad \text{on} \quad x = S(t). \tag{B.42}$$

The boundary condition for $u_0$ comes from imposing zero normal stress, $\sigma_{xx}$, at $x = S(t)$. Using

$\sigma_{xx} = -p - 2\mu(u_x + w_z)/3 + 2\mu u_x$, we arrive at the condition

$$\frac{\partial u_0}{\partial x} = \frac{1}{2}\Psi_n \bar{\phi}_0 g_{b0} \quad \text{on} \quad x = S(t). \tag{B.43}$$

#### B.7 Summary

The complete thin-film, extensional-flow model addressed in this work then consists of the differential equations (dropping the zero subscript on leading-order quantities and the overbar on averaged quantities)

$$\frac{\partial h}{\partial t} + \frac{\partial}{\partial x}(uh) = \Psi_n \phi g_b h \qquad \text{on} \quad 0 < x < S(t), \tag{B.44a}$$

$$\frac{\partial}{\partial t}(\phi h) + \frac{\partial}{\partial x}(\phi u h) = \Psi_n \phi g_b h - \Psi_d \phi h \qquad \text{on} \quad 0 < x < S(t), \tag{B.44b}$$

$$\frac{\partial g_s}{\partial t} = D\frac{\partial^2 g_s}{\partial x^2} - \delta Q^* (g_s - g_b) H(S(t) - x) \qquad \text{on} \quad 0 < x < 1/\delta, \tag{B.44c}$$

$$h\frac{\partial g_b}{\partial t} + \frac{\partial}{\partial x}(uhg_b) = \frac{\partial}{\partial x}\left(h\frac{\partial g_b}{\partial x}\right) + Q^*(g_s - g_b) - \Upsilon \phi g_b h \qquad \text{on} \quad 0 < x < S(t), \tag{B.44d}$$

$$4\frac{\partial}{\partial x}\left(h\frac{\partial u}{\partial x}\right) - \lambda^* u = 2\Psi_n \frac{\partial}{\partial x}(\phi g_b h) \qquad \text{on} \quad 0 < x < S(t), \tag{B.44e}$$

$$\frac{\mathrm{d}S}{\mathrm{d}t} = u(S(t), t), \tag{B.44f}$$

initial conditions

$$g_s(x, 0) = 1, \tag{B.45a}$$

$$g_b(x, 0) = 0, \tag{B.45b}$$

$$\phi(x, 0) = 0, \tag{B.45c}$$

$$h(x, 0) = H_0\left(1 - x^2\right), \tag{B.45d}$$

$$S(0) = 1, \tag{B.45e}$$

and boundary conditions

$$\frac{\partial h}{\partial x} = 0 \qquad \text{on} \quad x = 0, \tag{B.46a}$$

$$\frac{\partial \phi}{\partial x} = 0, \qquad \text{on} \quad x = 0, \tag{B.46b}$$

$$\frac{\partial g_s}{\partial x} = 0 \qquad \text{on} \quad x = 0, 1/\delta, \tag{B.46c}$$

$$\frac{\partial g_b}{\partial x} = 0 \qquad \text{on} \quad x = 0, S(t), \tag{B.46d}$$

$$u = 0 \qquad \text{on} \quad x = 0, \tag{B.46e}$$

$$\frac{\partial u}{\partial x} = \frac{1}{2}\Psi_n \phi g_b \qquad \text{on} \quad x = S(t). \tag{B.46f}$$

An alternative form of the active-phase mass-balance equation is available by expanding (B.44b) using the product rule and using (B.44a) to obtain

$$\frac{\partial \phi}{\partial t} + u\frac{\partial \phi}{\partial x} = \phi \left[\Psi_n g_b \left(1 - \phi\right) - \Psi_d\right] \quad \text{on} \quad 0 < x < S(t). \tag{B.47}$$

The form (B.47) is useful for the numerical method, which we now outline.

### C Numerical Methods

To solve the model numerically, we map the moving-boundary problem on $0 < x < S(t)$ to the unit interval using the change of variables

$$(\xi, \tau) = \left(\frac{x}{S(t)}, t\right) \implies \frac{\partial}{\partial x} = \frac{1}{S(\tau)}\frac{\partial}{\partial \xi}, \quad \frac{\partial}{\partial t} = \frac{\partial}{\partial \tau} - \frac{\xi S'(\tau)}{S(\tau)}\frac{\partial}{\partial \xi}. \tag{C.1}$$

To circumvent the Heaviside step function, we write Equation (B.44c) for the substratum nutrient concentration in piecewise form, and use the symbol $g_o$, defined for $S(t) < x < 1/\delta$, to denote the nutrient concentration in the region of the Petri dish unoccupied by the colony biofilm. Consequently, $g_s$ is then the nutrient concentration in the substratum, restricted to $0 < x < S(t)$. In distinguishing between $g_s$ and $g_o$, we also impose that the nutrient concentration in the agar defined over the entire Petri dish is a $C^1$ function, such that $g_s(S(t), t) = g_o(S(t), t)$ and $g_s'(S(t), t) = g_o'(S(t), t)$. The transformed model in terms of the new independent variables $\xi$ and $\tau$ to solve numerically is then

$$\frac{\partial h}{\partial \tau} - \frac{\xi u(1,\tau)}{S}\frac{\partial h}{\partial \xi} + \frac{1}{S}\frac{\partial}{\partial \xi}(uh) = \Psi_n \phi g_b h, \tag{C.2a}$$

$$\frac{\partial \phi}{\partial \tau} + \left[\frac{u - \xi u(1,\tau)}{S}\right]\frac{\partial \phi}{\partial \xi} = \phi\left[\Psi_n g_b\left(1-\phi\right) - \Psi_d\right], \tag{C.2b}$$

$$\frac{\partial g_s}{\partial \tau} - \frac{\xi u(1,\tau)}{S}\frac{\partial g_s}{\partial \xi} = \frac{D}{S^2}\frac{\partial^2 g_s}{\partial \xi^2} - \delta Q^*\left(g_s - g_b\right), \tag{C.2c}$$

$$\frac{\partial g_o}{\partial \tau} - \frac{\xi u(1,\tau)}{S}\frac{\partial g_o}{\partial \xi} = \frac{D}{S^2}\frac{\partial^2 g_o}{\partial \xi^2}, \tag{C.2d}$$

$$h\frac{\partial g_b}{\partial \tau} - \frac{\xi u(1,\tau) h}{S}\frac{\partial g_b}{\partial \xi} + \frac{1}{S}\frac{\partial}{\partial \xi}(uhg_b) = \frac{1}{S^2}\frac{\partial}{\partial \xi}\left(h\frac{\partial g_b}{\partial \xi}\right) + Q^*\left(g_s - g_b\right) - \Upsilon \phi g_b h, \tag{C.2e}$$

$$\frac{4}{S^2}\frac{\partial}{\partial \xi}\left(h\frac{\partial u}{\partial \xi}\right) - \lambda^* u = \frac{2\Psi_n}{S}\frac{\partial}{\partial \xi}\left(\phi g_b h\right), \tag{C.2f}$$

$$\frac{\mathrm{d}S}{\mathrm{d}\tau} = u(1,\tau). \tag{C.2g}$$

All PDEs are solved for $0 \leq \tau \leq T$, such that the PDEs (C.2a)–(C.2c), (C.2e) and (C.2f) are defined for $0 < \xi < 1$ and (C.2d) applies for $1 < \xi < 1/\delta S(\tau)$. The corresponding initial conditions are

$$\begin{gathered} g_s(\xi, 0) = 1, \quad g_o(\xi, 0) = 1, \quad g_b(\xi, 0) = 0, \quad \phi(\xi, 0) = 0, \\ h(\xi, 0) = H_0\left(1 - \xi^2\right), \quad S(0) = 1, \end{gathered} \tag{C.3}$$

and the boundary conditions are

$$
\begin{gathered}
\left.\frac{\partial h}{\partial \xi}\right|_{(0,\tau)} = 0, \quad \left.\frac{\partial \phi}{\partial \xi}\right|_{(0,\tau)} = 0, \\
\left.\frac{\partial g_s}{\partial \xi}\right|_{(0,\tau)} = 0, \quad \left.\frac{\partial g_o}{\partial \xi}\right|_{(1/\delta S,\tau)} = 0, \quad \left.\frac{\partial g_b}{\partial \xi}\right|_{(0,\tau)} = \left.\frac{\partial g_b}{\partial \xi}\right|_{(1,\tau)} = 0, \\
g_s(1,\tau) = g_o(1,\tau), \quad \left.\frac{\partial g_s}{\partial \xi}\right|_{(1,\tau)} = \left.\frac{\partial g_o}{\partial \xi}\right|_{(1,\tau)}, \\
u(0,\tau) = 0, \quad \left.\frac{\partial u}{\partial \xi}\right|_{(1,\tau)} = \frac{S(\tau)\Psi_n}{2}\phi(1,\tau)g_b(1,\tau).
\end{gathered}
\tag{C.4}
$$

To describe the numerical scheme, we use the notation $u_j^l = u(\xi_j, \tau_l)$ to represent the value of variables at grid points. We define the temporal grid $\tau_l = (l-1)\Delta\tau$, for $l = 1, \ldots, M$, such that $M$ is the number of time steps, and $\Delta\tau = T/(M-1)$ is the constant time-step size. We define the spatial grid for the colony biofilm to be $\xi_j = (j-1)\Delta\xi$ for $j = 1, \ldots, N$, where $N$ is the number of grid points, and $\Delta\xi = 1/(N-1)$ is the constant grid spacing. In unoccupied regions of the substratum, we define $\xi_j^o = 1 + (j-1)\Delta\xi^o$, where the constant grid spacing $\Delta\xi^o = (1/\delta S - 1)/(N-1)$ is chosen such that $\xi_N^o = 1/\delta S$. Since the grid $\xi^o$ depends on $\tau$, we reinitialise it every time step after updating $S$. We use a Crank–Nicolson method to solve (C.2), similar to Tam [3] and Tam et al. [4]. We use an upwind spatial discretisation for the advection term in (C.2a), and central differences for spatial derivatives arising from applying the change of variables to derivatives with respect to $t$. At interior grid points, this leads to the scheme

$$\frac{h_j^{l+1}-h_j^l}{\Delta\tau}-\frac{\xi_j u_N^l}{S^l}\left(\frac{h_{j+1}^{l+1/2}-h_{j-1}^{l+1/2}}{2\Delta\xi}\right)+\frac{u_j^l h_j^{l+1/2}-u_{j-1}^l h_{j-1}^{l+1/2}}{S^l\Delta\xi}=\Psi_n\phi_j^l {g_b}_j^l h_j^{l+1/2}, \tag{C.5a}$$

$$\frac{\phi_j^{l+1}-\phi_j^l}{\Delta\tau}+\left(\frac{u_j^l-\xi_j u_N^l}{S^l}\right)\left(\frac{\phi_{j+1}^{l+1/2}-\phi_{j-1}^{l+1/2}}{2\Delta\xi}\right)=\phi_j^{l+1/2}\left[\Psi_n {g_b}_j^l\left(1-\phi_j^l\right)-\Psi_d\right], \tag{C.5b}$$

$$\begin{aligned}&\frac{{g_s}_j^{l+1}-{g_s}_j^l}{\Delta\tau}-\frac{\xi_j u_N^l}{S^l}\left(\frac{{g_s}_{j+1}^{l+1/2}-{g_s}_{j-1}^{l+1/2}}{2\Delta\xi}\right)\\&=\frac{D}{{S^l}^2}\left(\frac{{g_s}_{j+1}^{l+1/2}-2{g_s}_j^{l+1/2}+{g_s}_{j-1}^{l+1/2}}{\Delta\xi^2}\right)-\delta Q^*\left({g_s}_j^{l+1/2}-{g_b}_j^l\right),\end{aligned} \tag{C.5c}$$

$$\frac{{g_o}_j^{l+1}-{g_o}_j^l}{\Delta\tau}-\frac{\xi_j^o u_N^l}{S^l}\left(\frac{{g_o}_{j+1}^{l+1/2}-{g_o}_{j-1}^{l+1/2}}{2\Delta\xi^o}\right)=\frac{D}{{S^l}^2}\left(\frac{{g_o}_{j+1}^{l+1/2}-2{g_o}_j^{l+1/2}+{g_o}_{j-1}^{l+1/2}}{{\Delta\xi^o}^2}\right), \tag{C.5d}$$

$$\begin{aligned}&h_j^{l+1}\left(\frac{{g_b}_j^{l+1}-{g_b}_j^l}{\Delta\tau}\right)-\frac{\xi_j u_N^l}{S^l}\left(\frac{{g_b}_{j+1}^{l+1/2}-{g_b}_{j-1}^{l+1/2}}{2\Delta\xi}\right)+\frac{u_{j+1}^l h_{j+1}^{l+1}{g_b}_{j+1}^{l+1/2}-u_{j-1}^l h_{j-1}^{l+1}{g_b}_{j-1}^{l+1/2}}{2\Delta\xi S^l}\\&=\frac{\left(h_{j+1}^{l+1}+h_j^{l+1}\right)\left({g_b}_{j+1}^{l+1/2}-{g_b}_j^{l+1/2}\right)-\left(h_j^{l+1}+h_{j-1}^{l+1}\right)\left({g_b}_j^{l+1/2}-{g_b}_{j-1}^{l+1/2}\right)}{2{S^l}^2\Delta\xi^2}\\&\quad+Q^*\left({g_s}_j^{l+1}-{g_b}_j^{l+1/2}\right)-\Upsilon\phi_j^{l+1}{g_b}_j^{l+1/2}h_j^{l+1},\end{aligned} \tag{C.5e}$$

$$\begin{aligned}&\frac{2}{{S^l}^2}\frac{\left(h_{j+1}^l+h_j^l\right)\left(u_{j+1}^l-u_j^l\right)-\left(h_j^l+h_{j-1}^l\right)\left(u_j^l-u_{j-1}^l\right)}{\Delta\xi^2}-\lambda^* u_j^l\\&=\frac{\Psi_n}{S^l}\left(\frac{\phi_{j+1}^l {g_b}_{j+1}^l h_{j+1}^l-\phi_{j-1}^l {g_b}_{j-1}^l h_{j-1}^l}{\Delta\xi}\right),\end{aligned} \tag{C.5f}$$

$$\frac{S^{l+1}-S^l}{\Delta\tau}=\frac{u_N^{l+1/2}}{2}, \tag{C.5g}$$

for $j=2,\dots,N-1$, and $l=1,\dots M-1$. In (C.5), for compactness we write the scheme with some terms approximated halfway between time steps. In practice, we approximate these terms using centred averages, for example

$$u_j^{l+1/2}=\frac{u_j^{l+1}-u_j^l}{2}, \tag{C.6}$$

and so on for other variables. Since $\xi^o$ depends on $S$, after updating $S$ based on (C.5g) we interpolate linearly to update the nutrient concentrations $g_s$ and $g_o$, to maintain compatibility with the updated colony-biofilm domain.

It remains to specify the numerical schemes for the boundary conditions. By the same argument as Ward and King [11], since $h(S(0),0)=0$ the PDE (B.44a) guarantees that $h(S(t),t)=0$ for all $t$. Therefore, for convenience we impose $h_N=0$ directly. To ensure continuity of the nutrient concentration in the substratum, we apply the Dirichlet conditions $g_s(1,\tau)=g_o(1,\tau)=a$. The correct value of $a$ is such that $\partial_\xi g_s(1,\tau)=\partial_{\xi^o} g_o'(1,\tau)$, and we obtain this value numerically using the Newton–Raphson method, using $g_s$ at $\xi=1$ from the previous time step as the initial guess. Once we have found $a$, the numerical schemes for the boundary conditions read

$$\frac{-3h_1^l + 4h_2^l - h_3^l}{2\Delta\xi} = 0, \tag{C.7a}$$

$$h_N = 0, \tag{C.7b}$$

$$\frac{-3\phi_1^l + 4\phi_2^l - \phi_3^l}{2\Delta\xi} = 0, \tag{C.7c}$$

$$\frac{\phi_N^{l+1} - \phi_N^l}{\Delta\tau} = \phi_N^{l+1/2}\left[\Psi_n {g_b}_N^l \left(1 - \phi_N^l\right) - \Psi_d\right], \tag{C.7d}$$

$$\frac{{g_s}_1^{l+1} - {g_s}_1^l}{\Delta\tau} = \frac{2D}{S^{l2}}\left(\frac{{g_s}_2^{l+1/2} - {g_s}_1^{l+1/2}}{\Delta\xi^2}\right) - \delta Q^*\left({g_s}_1^{l+1/2} - {g_b}_1^l\right), \tag{C.7e}$$

$${g_s}_N^{l+1} = a, \tag{C.7f}$$

$${g_o}_1^{l+1} = a, \tag{C.7g}$$

$$\frac{{g_o}_N^{l+1} - {g_o}_N^l}{\Delta\tau} = \frac{2D}{S^{l2}}\left(\frac{{g_o}_{j-1}^{l+1/2} - {g_o}_j^{l+1/2}}{\Delta\xi^{o2}}\right), \tag{C.7h}$$

$$\begin{aligned} h_1^{l+1}\left(\frac{{g_b}_1^{l+1} - {g_b}_1^l}{\Delta\tau}\right) + \frac{h_1^{l+1}{g_b}_1^{l+1}}{S^l}\left(\frac{-3u_1^l + 4u_2^l - u_3^l}{2\Delta\xi}\right) \\ = \frac{\left(h_2^{l+1} + h_1^{l+1}\right)\left({g_b}_2^{l+1/2} - {g_b}_1^{l+1/2}\right)}{S^{l2}\Delta\xi^2} + Q^*\left({g_s}_1^l - {g_b}_1^{l+1/2}\right) - \Upsilon\phi_1^{l+1}{g_b}_1^{l+1/2}h_1^{l+1}, \end{aligned} \tag{C.7i}$$

$${g_b}_N^{l+1} = Q^*{g_s}_N^{l+1}\left[Q^* + u_N^l\left(\frac{3h_N^{l+1} - 4h_{N-1}^{l+1} + h_{N-2}^{l+1}}{2\Delta\xi}\right)\right]^{-1}, \tag{C.7j}$$

$$u_1^l = 0, \tag{C.7k}$$

$$\frac{3u_N^l - 4u_{N-1}^l + u_{N-2}^l}{2\Delta\xi} = \frac{S^l\Psi_n}{2}\phi_N^l {g_b}_N^l. \tag{C.7l}$$

The expression (C.7j) arises by expanding the PDE (C.2e), applying $h = 0$ and $\partial_\xi g_b = 0$ at $\xi = 1$, and discretising the resulting expression. Together with the initial conditions (C.3), this completes the scheme for solving the full model (B.44)–(B.46) numerically.

#### C.1 Parameter Optimisation Tests

**Repeated Parameter Estimation:** Table C.1 presents parameter-estimation results for five independent runs of the optimisation routine with the mean experimental data. These results confirm that the optimisation procedure converges to a very similar local minimum in $\rho(\theta^*, a)$ each run.

**Numerical Grid Convergence:** Since each evaluation of the objective function in the parameter estimation method requires solving the model numerically once, the parameter estimation procedure is computationally expensive. With $N_\xi = 401$ grid points and $N_\tau = 1601$ time steps, obtaining the optimal parameters for one set of experimental data took approximately 6.5 hours on an Intel Core i7-3770 CPU @ 3.40GHz × 4 running Linux Mint 22.2. Owing to the

Table C.1: Optimal dimensionless parameters $\theta^*(a)$, expressed to four significant figures, for each agar density based on mean experimental data and 5 repetitions of the optimisation routine.

| Run | $a$ | $\Psi_n$ | $\Psi_d$ | $Q^*$ | $\Upsilon$ | $\lambda^*$ | $\rho(\theta^*, a)$ |
|---|---|---|---|---|---|---|---|
| 1 | 0.6 | 0.3030 | 0.007766 | 7.400 | 7.881 | 0.6733 | 0.04374 |
| | 0.8 | 0.3651 | 0.007471 | 6.539 | 7.336 | 1.042 | 0.05364 |
| | 1.2 | 0.4093 | 0.007551 | 4.481 | 6.462 | 1.848 | 0.06360 |
| | 2.0 | 0.3022 | 0.008347 | 3.270 | 5.007 | 2.670 | 0.06254 |
| 2 | 0.6 | 0.3009 | 0.007768 | 7.601 | 7.861 | 0.6693 | 0.04372 |
| | 0.8 | 0.3651 | 0.007473 | 6.533 | 7.332 | 1.042 | 0.05364 |
| | 1.2 | 0.4091 | 0.007552 | 4.480 | 6.459 | 1.847 | 0.06360 |
| | 2.0 | 0.3022 | 0.008346 | 3.271 | 5.009 | 2.670 | 0.06254 |
| 3 | 0.6 | 0.3011 | 0.007765 | 7.602 | 7.867 | 0.6696 | 0.04372 |
| | 0.8 | 0.3650 | 0.007471 | 6.551 | 7.335 | 1.041 | 0.05364 |
| | 1.2 | 0.4095 | 0.007551 | 4.474 | 6.463 | 1.849 | 0.06360 |
| | 2.0 | 0.3023 | 0.008344 | 3.275 | 5.015 | 2.671 | 0.06254 |
| 4 | 0.6 | 0.3013 | 0.007766 | 7.579 | 7.869 | 0.6700 | 0.04372 |
| | 0.8 | 0.3646 | 0.007472 | 6.561 | 7.330 | 1.041 | 0.05364 |
| | 1.2 | 0.4099 | 0.007548 | 4.479 | 6.475 | 1.850 | 0.06360 |
| | 2.0 | 0.3022 | 0.008347 | 3.272 | 5.008 | 2.670 | 0.06254 |
| 5 | 0.6 | 0.3009 | 0.007767 | 7.610 | 7.862 | 0.6692 | 0.04372 |
| | 0.8 | 0.3648 | 0.007473 | 6.550 | 7.330 | 1.041 | 0.05364 |
| | 1.2 | 0.4092 | 0.007553 | 4.477 | 6.457 | 1.848 | 0.06360 |
| | 2.0 | 0.3024 | 0.008344 | 3.273 | 5.014 | 2.672 | 0.06254 |

constraint of computational cost, we use a relatively coarse numerical grid to obtain the results. These numerical solutions will contain numerical error, so we repeat the parameter estimation using different grid spacing and time step sizes, to ensure that our conclusions remain valid as the numerical grid is refined.

Table C.2: Parameter-estimation results for different grid spacing and time-step sizes, and 0.6% agar.

| Parameter | $N_\xi$ \ $N_\tau$ | 101 | 201 | 401 | 801 | 1601 |
|---|---|---|---|---|---|---|
| $\Psi_n$ | 25 | 0.3014 | 0.3033 | 0.3016 | 0.3019 | 0.3009 |
| | 51 | 0.3008 | 0.3029 | 0.3014 | 0.3023 | 0.3012 |
| | 101 | 0.3002 | 0.3023 | 0.3010 | 0.3018 | 0.3009 |
| | 201 | 0.2993 | 0.3018 | 0.3002 | 0.3008 | 0.3002 |
| | 401 | 0.2982 | 0.3009 | 0.2995 | 0.2998 | 0.2992 |
| $\Psi_d$ | 25 | 0.007586 | 0.007662 | 0.007662 | 0.007695 | 0.007699 |
| | 51 | 0.007639 | 0.007721 | 0.007720 | 0.007754 | 0.007754 |
| | 101 | 0.007687 | 0.007768 | 0.007768 | 0.007802 | 0.007802 |
| | 201 | 0.007729 | 0.007811 | 0.007808 | 0.007843 | 0.007844 |
| | 401 | 0.007759 | 0.007844 | 0.007840 | 0.007875 | 0.007876 |
| $Q^*$ | 25 | 11.31 | 10.20 | 10.08 | 9.942 | 9.961 |
| | 51 | 9.316 | 8.549 | 8.455 | 8.291 | 8.349 |
| | 101 | 8.292 | 7.617 | 7.591 | 7.462 | 7.498 |
| | 201 | 7.721 | 7.060 | 7.062 | 6.986 | 6.996 |
| | 401 | 7.413 | 6.757 | 6.744 | 6.692 | 6.704 |
| $\Upsilon$ | 25 | 8.745 | 8.780 | 8.747 | 8.759 | 8.733 |
| | 51 | 8.219 | 8.228 | 8.203 | 8.223 | 8.206 |
| | 101 | 7.890 | 7.882 | 7.862 | 7.879 | 7.867 |
| | 201 | 7.664 | 7.648 | 7.623 | 7.637 | 7.628 |
| | 401 | 7.504 | 7.490 | 7.470 | 7.479 | 7.467 |
| $\lambda^*$ | 25 | 0.6958 | 0.7239 | 0.7300 | 0.7354 | 0.7352 |
| | 51 | 0.6608 | 0.6839 | 0.6900 | 0.6963 | 0.6960 |
| | 101 | 0.6424 | 0.6640 | 0.6694 | 0.6750 | 0.6752 |
| | 201 | 0.6305 | 0.6518 | 0.6569 | 0.6614 | 0.6623 |
| | 401 | 0.6223 | 0.6436 | 0.6492 | 0.6532 | 0.6539 |
| $\rho(\theta^*, a)$ | 25 | 0.03920 | 0.03967 | 0.04041 | 0.04045 | 0.04040 |
| | 51 | 0.04136 | 0.04162 | 0.04234 | 0.04241 | 0.04236 |
| | 101 | 0.04230 | 0.04310 | 0.04372 | 0.04379 | 0.04375 |
| | 201 | 0.04413 | 0.04430 | 0.04485 | 0.04488 | 0.04483 |
| | 401 | 0.04487 | 0.04516 | 0.04573 | 0.04573 | 0.04565 |

Table C.3: Parameter-estimation results for different grid spacing and time-step sizes, and 2.0% agar.

| Parameter | $N_\xi$ \ $N_\tau$ | 101 | 201 | 401 | 801 | 1601 |
|---|---|---|---|---|---|---|
| $\Psi_n$ | 25 | 0.3032 | 0.3059 | 0.3043 | 0.3054 | 0.3042 |
| | 51 | 0.3016 | 0.3042 | 0.3041 | 0.3048 | 0.3044 |
| | 101 | 0.3001 | 0.3023 | 0.3022 | 0.3031 | 0.3028 |
| | 201 | 0.2979 | 0.3002 | 0.2999 | 0.3005 | 0.3003 |
| | 401 | 0.2954 | 0.2979 | 0.2976 | 0.2983 | 0.2979 |
| $\Psi_d$ | 25 | 0.008137 | 0.008201 | 0.008203 | 0.008230 | 0.008238 |
| | 51 | 0.008216 | 0.008281 | 0.008276 | 0.008309 | 0.008308 |
| | 101 | 0.008282 | 0.008352 | 0.008346 | 0.008378 | 0.008377 |
| | 201 | 0.008342 | 0.008411 | 0.008407 | 0.008437 | 0.008439 |
| | 401 | 0.008392 | 0.008460 | 0.008455 | 0.008486 | 0.008488 |
| $Q^*$ | 25 | 4.950 | 4.635 | 4.626 | 4.573 | 4.580 |
| | 51 | 4.063 | 3.813 | 3.767 | 3.716 | 3.726 |
| | 101 | 3.525 | 3.299 | 3.273 | 3.225 | 3.228 |
| | 201 | 3.205 | 2.996 | 2.967 | 2.936 | 2.938 |
| | 401 | 3.028 | 2.810 | 2.789 | 2.760 | 2.768 |
| $\Upsilon$ | 25 | 6.287 | 6.240 | 6.213 | 6.228 | 6.201 |
| | 51 | 5.575 | 5.519 | 5.505 | 5.497 | 5.497 |
| | 101 | 5.102 | 5.015 | 5.010 | 5.001 | 5.000 |
| | 201 | 4.771 | 4.679 | 4.664 | 4.657 | 4.658 |
| | 401 | 4.548 | 4.446 | 4.436 | 4.429 | 4.432 |
| $\lambda^*$ | 25 | 2.990 | 3.088 | 3.098 | 3.124 | 3.117 |
| | 51 | 2.724 | 2.801 | 2.824 | 2.842 | 2.843 |
| | 101 | 2.585 | 2.650 | 2.670 | 2.687 | 2.689 |
| | 201 | 2.495 | 2.556 | 2.574 | 2.587 | 2.590 |
| | 401 | 2.434 | 2.493 | 2.511 | 2.524 | 2.526 |
| $\rho(\theta^*, a)$ | 25 | 0.05883 | 0.05908 | 0.05996 | 0.05978 | 0.05981 |
| | 51 | 0.06039 | 0.06058 | 0.06145 | 0.06128 | 0.06131 |
| | 101 | 0.06161 | 0.06170 | 0.06254 | 0.06237 | 0.06240 |
| | 201 | 0.06253 | 0.06266 | 0.06344 | 0.06327 | 0.06330 |
| | 401 | 0.06322 | 0.06336 | 0.06416 | 0.06397 | 0.06399 |

The convergence results in Tables C.2 and C.3 suggest that $N_\tau$ has no consistent effect on the estimated parameter values when $N_\tau > 201$. Consequently, we consider the results with $N_\tau = 1601$ to be approximately independent of time-step size, $\Delta\tau$. However, the results may not be independent of $\Delta\xi$, because the parameter estimates in Tables C.2 and C.3 vary with $N_\xi$. Doubling the number of grid points and time steps yields small changes in $\Psi_n$, $\Psi_d$, and $\lambda^*$, and up to 10% variation in $Q^*$ and $\Upsilon$. These results align with the sensitivity analyses in the main manuscript, where $Q^*$ was more difficult to identify, and $Q^*$ and $\Upsilon$ tended to vary in concert with each other. The variation in optimal parameters could also be due to randomness inherent in the optimisation procedure, in addition to the grid spacing. As Tables C.2 and C.3 indicate, these variations do not affect the qualitative conclusions of the manuscript.

#### C.2 Heat Maps

See below of additional parameter-pair heat maps indicating how model parameters influence $\rho(\theta, a)$, the distance between the model and experimental results. Figure C.1 shows that the parameter trends on high-density agar are similar to the trends on low-density agar.

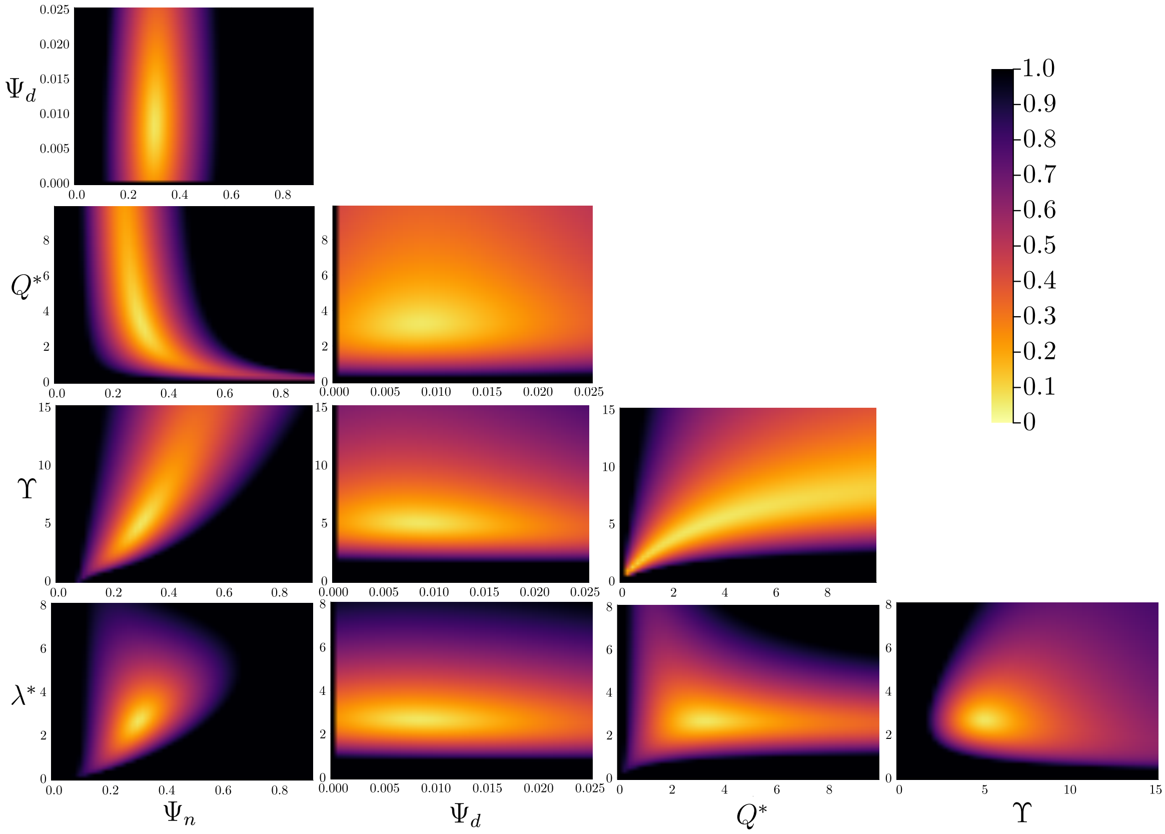

Figure C.1: Parameter-pair heat maps for hard 2.0% agar. Plots represent the distance between the numerical solution and experimental data, $\rho(\theta, a)$, for given parameter combination. Unless otherwise stated, parameters take the optimal values. When varied, each parameter ranges from zero to three times the optimal value.

### D Code and Data Availability

Experimental data, numerical code, and additional results for image processing and solving the thin-film model are available on GitHub.

### References

[1] W. J. Middelhoven, B. Broekhuizen, and J. van Eijk, "Detection, with the Dye Phloxine B, of Yeast Mutants Unable to Utilize Nitrogenous Substances as the Sole Nitrogen Source", *J. Bacteriol.* 128 (1976), pp. 851–852, DOI: 10.1128/jb.128.3.851-852.1976.

[2] J. F. Cannon, J. B. Gibbs, and K. Tatchell, "Supperssors of the Ras2 Mutation of *Saccharomyces Cerevisiae*", *Genetics* 113 (1986), pp. 247–264, DOI: 10.1093/genetics/113.2.247.

[3] A. Tam, "Mathematical Modelling of Pattern Formation in Yeast Biofilms", PhD thesis, The University of Adelaide, 2019, HDL: 2440/122613.

[4] A. Tam, J. E. F. Green, S. Balasuriya, E. L. Tek, J. M. Gardner, J. F. Sundstrom, V. Jiranek, and B. J. Binder, "A Thin-Film Extensional Flow Model for Biofilm Expansion by Sliding Motility", *Proc. Royal Soc. A* 475 (2019), 20190175, DOI: 10.1098/rspa.2019.0175.

[5] A. K. Y. Tam, B. Harding, J. E. F. Green, S. Balasuriya, and B. J. Binder, "Thin-Film Lubrication Model for Biofilm Expansion under Strong Adhesion", *Phys. Rev. E* 105 (2022), 014408, DOI: 10.1103/PhysRevE.105.014408.

[6] S. Srinivasan, C. N. Kaplan, and L. Mahadevan, "A Multiphase Theory for Spreading Microbial Swarms and Films", *eLife* 8 (2019), e42697, DOI: 10.7554/eLife.42697.

[7] D. A. Drew, "Mathematical Modeling of Two-Phase Flow", *Annu. Rev. Fluid Mech.* 15 (1983), pp. 261–291, DOI: 10.1146/annurev.fl.15.010183.001401.

[8] I. Klapper, C. J. Rupp, R. Cargo, B. Purvedorj, and P. Stoodley, "Viscoelastic Fluid Description of Bacterial Biofilm Material Properties", *Biotechnol. Bioeng.* 80 (2002), pp. 289–296, DOI: 10.1002/bit.10376.

[9] S. J. Franks, H. M. Byrne, J. R. King, J. C. E. Underwood, and C. E. Lewis, "Modelling the Early Growth of Ductal Carcinoma in Situ of the Breast", *J. Math. Biol.* 47 (2003), pp. 424–452, DOI: 10.1007/s00285-003-0214-x.

[10] R. D. O'Dea, S. L. Waters, and H. M. Byrne, "A Multiphase Model for Tissue Construct Growth in a Perfusion Bioreactor", *Math. Med. Biol.* 27 (2010), pp. 95–127, DOI: 10.1093/imammb/dqp003.

[11] J. P. Ward and J. R. King, "Thin-Film Modelling of Biofilm Growth and Quorum Sensing", *J. Eng. Math.* 73 (2012), pp. 71–92, DOI: 10.1007/s10665-011-9490-4.

[12] G. Lemon, J. R. King, H. M. Byrne, O. E. Jensen, and K. M. Shakesheff, "Mathematical Modelling of Engineered Tissue Growth Using a Multiphase Porous Flow Mixture Theory", *J. Math. Biol.* 52 (2006), pp. 571–594, DOI: 10.1007/s00285-005-0363-1.

[13] D. A. Drew and L. A. Segel, "Averaged Equations for Two-Phase Flows", *Stud. Appl. Math.* 50 (1971), pp. 205–231, DOI: 10.1002/sapm1971503205.

[14] G. Forgacs, R. A. Foty, Y. Shafrir, and M. S. Steinberg, "Viscoelastic Properties of Living Embryonic Tissues: A Quantitative Study", *Biophys. J.* 74 (1998), pp. 2227–2234, DOI: 10.1016/S0006-3495(98)77932-9.

[15] R. M. Harshey, "Bacterial Motility on a Surface: Many Ways to a Common Goal", *Annu. Rev. Microbiol.* 57 (2003), pp. 249–273, DOI: 10.1146/annurev.micro.57.030502.091014.

[16] J. Recht, A. Martínez, S. Torello, and R. Kolter, "Genetic Analysis of Sliding Motility in *Mycobacterium smegmatis*", *J. Bacteriol.* 182 (2000), pp. 4348–4351, DOI: 10.1128/JB.182.15.4348-4351.2000.

[17] R. D. O'Dea, S. L. Waters, and H. M. Byrne, "A Two-Fluid Model for Tissue Growth within a Dynamic Flow Environment", *Eur. J. Appl. Math.* 19 (2008), pp. 607–634, DOI: 10.1017/S0956792508007687.

[18] S. J. Franks and J. R. King, "Interactions between a Uniformly Proliferating Tumour and Its Surroundings: Uniform Material Properties", *Math. Med. Biol.* 20 (2003), pp. 47–89, DOI: 10.1093/imammb/20.1.47.

[19] S. R. Lubkin and T. L. Jackson, "Multiphase Mechanics of Capsule Formation in Tumors", *J. Biomech. Eng.* 124 (2002), pp. 237–243, DOI: 10.1115/1.1427925.

[20] A. Tam, J. E. F. Green, S. Balasuriya, E. L. Tek, J. M. Gardner, J. F. Sundstrom, V. Jiranek, and B. J. Binder, "Nutrient-Limited Growth with Non-Linear Cell Diffusion as a Mechanism for Floral Pattern Formation in Yeast Biofilms", *J. Theor. Biol.* 448 (2018), pp. 122–141, DOI: 10.1016/j.jtbi.2018.04.004.
